# Parasite and pathogen prevalence in our closest animal companions is determined by accessibility of sanitation services

**DOI:** 10.1101/2022.03.08.483406

**Authors:** Kayleigh Chalkowski, Christopher A. Lepczyk, Alan E. Wilson, Kim Valenta, Breanna Sipley, Joi Brownlee, Abbey M. Morgan, Scott R. Santos, Janna R. Willoughby, Sarah Zohdy

## Abstract

Despite the critical role parasites play in ecosystem functioning and their considerable influence on human society, little is known about their variations in abundance on a global scale. This gap in knowledge is amplified by a lack of holistic understanding on how the abundance of parasites of wildlife and humans varies across environmental and socioeconomic gradients, despite a need to integrate study of parasites across social and environmental spheres. Free-roaming companion animals (e.g., domestic cats (*Felis catus*) and dogs (*Canis lupus familiaris*)) share pathogens and have frequent contact with humans and wildlife. Thus, they are an effective model to understand how parasite and pathogen prevalence of humans and wildlife varies across environmental and socioeconomic gradients. Through a global systematic review and analysis of socioeconomic and environmental variables, including per capita GDP, income disparity, sanitation, biodiversity, island habitation, and latitude, we find that sanitation and island habitation best explained free-roaming companion animal parasite and pathogen prevalence. Sanitation was significantly associated with parasite and pathogen prevalence in free-roaming companion animals, such that for every 10% increase in the proportion of the human population with improved sanitation access, parasite and pathogen prevalence in free-roaming companion animals decreased by 12% (5-19%, 95% C.L.; *p* = 0.0023). Since companion animals share many parasites with humans and wildlife, these results suggest that actionable interventions to improve sanitation access could reduce parasite and pathogen exposure risks from companion animals to humans and wildlife.

**Significance Statement:** In addition to playing a critical role in ecosystem functioning, parasites also influence human health, behavior, and society. Further, parasites are also impacted by human activities, as much as by ecological phenomena in natural environments. Despite these dualities, little is known about their variations in abundance on a global scale across environmental and socioeconomic gradients. Using free-roaming companion animals (e.g., domestic cats (*Felis catus*) and dogs (*Canis lupus familiaris*)) as a model system, we find that access to safely managed sanitation services is strongly associated with parasite and pathogen prevalence. This finding underscores improvements to sanitation as an actionable One Health intervention that could reduce parasite and pathogen exposure risks from companion animals to humans and wildlife.

## Introduction

Nature has had far-reaching impacts on many aspects of human society including human health, psychology, the economy, and policy (Carleton and Hsiang 2016, Bratman et al. 2019, Soga and Gaston 2020). Likewise, human society also has considerable influence on the natural world including impacts on climate (Yuan et al. 2019, Madakumbura et al. 2021, Kishore et al. 2022) and biological communities, from microscopic organisms to charismatic megafauna (Burney and Flannery 2005, Cavicchioli et al. 2019, Hill et al. 2020, Adhikari et al. 2022). Parasitic organisms (i.e., parasites and pathogens) are one such component of ecosystems that have had considerable influence on human culture, behavior, and even the very course of human evolution (Lafferty 2006, Vannier-Santos and Lenzi 2011), and likewise may be affected by human society in return (Budria and Candolin 2013). Yet, we currently have only a vague understanding of how parasites and pathogens in humans and wildlife are distributed across environmental and socioeconomic gradients (Pappalardo et al. 2019). This is problematic, because in addition to their influence on human society, they also play key roles in population regulation as well as ecosystem functioning and resilience (Hudson et al. 2006, Wood and Johnson 2015, Frainer et al. 2018). This lack of understanding is also troublesome since a growing need to address planetary health problems with a One Health approach (i.e., the idea that the health of humans and animals are interlinked (Schwabe 1984)) necessitates integrated understanding of parasite and pathogen distribution across both environmental and socioeconomic gradients (Jenkins et al. 2015, Sleeman et al. 2019).

Environmental gradients can drastically alter parasite and pathogen distribution and abundance (Guernier et al. 2004, Jones et al. 2008, Rohr et al. 2020). In fact, gradients such as latitude, biodiversity, and inhabitation of islands have been well-explored in wildlife with some inconsistencies in direction of effect due to nuances in parasite and pathogen biology, scale, and habitat (Table 1). However, these results are difficult to integrate with respect to parasites infecting humans given that such studies have often focused on disease burdens and emergence rather than metrics of parasite abundance (i.e., prevalence, infection intensity). For example, our understanding of whether biodiversity is positively (i.e., amplification effect hypothesis) or negatively (i.e., dilution effect hypothesis) associated with parasite and pathogen abundance has been well studied in wildlife (Keesing et al. 2006, Rohr et al. 2020) but findings in human populations conflict with these results (Wood et al. 2017, Rohr et al. 2020). This difference is likely due to the difficulty with isolating underlying drivers between human and wildlife studies due in part to the use of different metrics (e.g., disability-affected life years vs. parasite prevalence or abundance; Rohr et al. 2020). Another example of an important environmental gradient is summarized by the island syndrome hypothesis, which predicts greater parasite prevalence on islands (Dybing et al. 2018). While this pattern has been observed in wildlife (Literák et al. 2015, Dybing et al. 2018) and there is evidence that parasites in humans have reduced species richness on islands (Jean et al. 2015), the effect of islands on parasite prevalence has not yet been investigated in humans. Thus, further study of the relationship between environmental factors and human parasites and pathogens is critically needed to adequately address public health concerns.

**Table 1.**
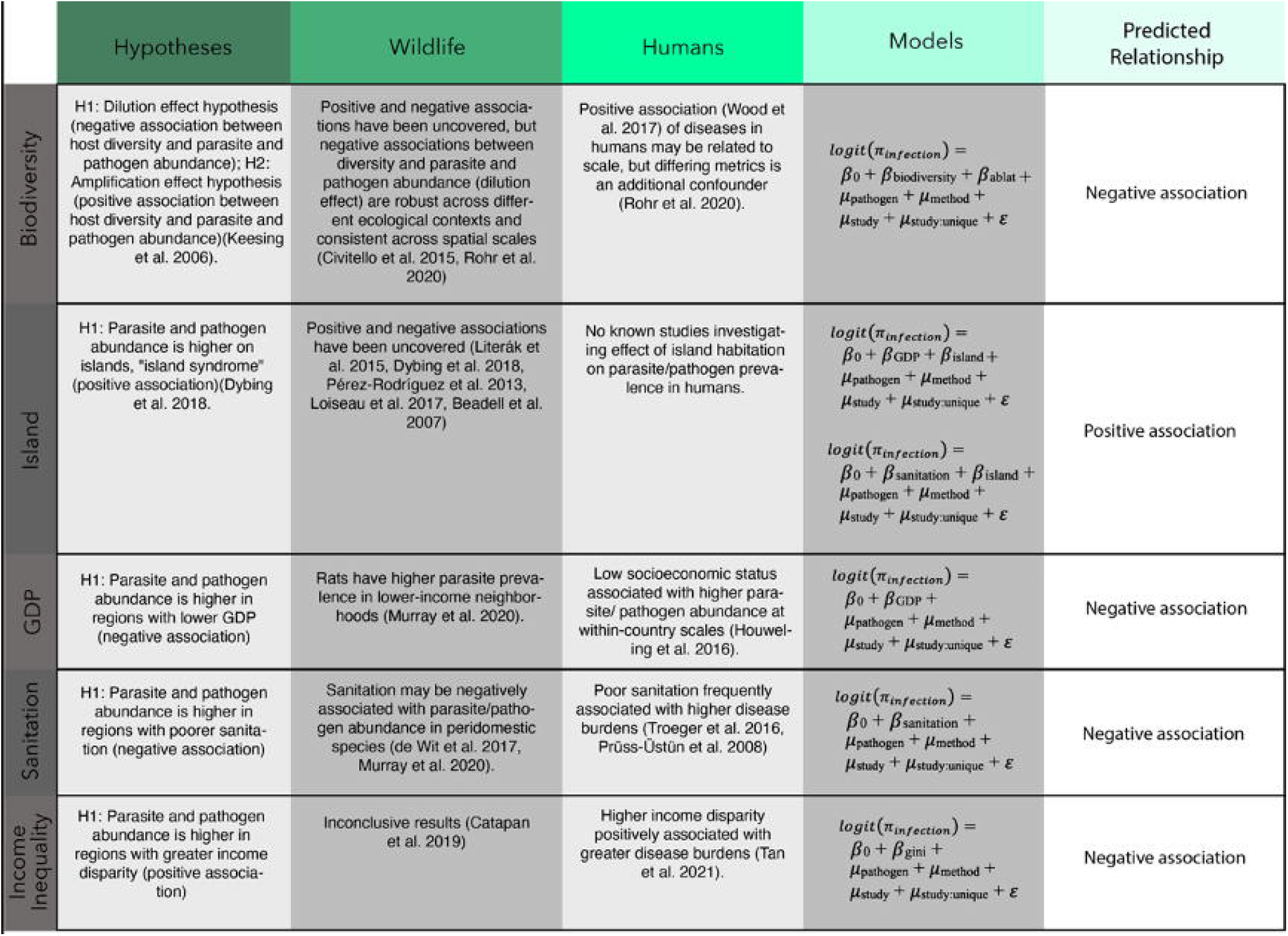
Summary of hypotheses; parasite and pathogen abundance patterns across socioeconomic and environmental gradients in previous research on wildlife and humans; our models tested in the present study; and the corresponding predicted relationships. Logit(π_infection_)-probability of infection; *β*- fixed effects; *β*_0_- intercept; *µ*- random effect; *ε*- residual error; pathogen- pathogen species; method- detection method used to identify pathogen in prevalence study; study-unique identifier for each study; study: unique-each unique pathogen prevalence value nested within each study (because some studies evaluated multiple pathogens); sanitation-proportion of population with access to safely managed sanitation services; island-whether or not free-roaming cat dog was on an island; gdp-per capita gross domestic product of country where each study was conducted; biodiversity-national biodiversity index of the country where each study was conducted; ablat-absolute latitude based on center-point of the country where each study was conducted; gini-gini coefficient of income inequality of the country where each study was conducted.

Socioeconomic gradients such as gross domestic product (GDP), sanitation, and income disparity have been well-explored in humans, with the general consensus that low GDP, poor sanitation, and high income disparity is associated with greater disease burdens (Hotez 2002, Sachs and Malaney 2002, Pru□ss-U□stu□n and World Health Organization 2008, Pru□ss-U□stu□n et al. 2016, Troeger et al. 2018, Tan et al. 2021; Table 1). While there has been markedly less attention on how socioeconomic factors influence wildlife health, some findings suggest similar associations as in humans while others have been relatively unexplored (Table 1). For example, low-income neighborhoods are associated with greater parasite prevalence in rats (Murray et al. 2020), and parasite prevalence in peri-domestic species (i.e., species associated with humans) on islands may be greater in areas with poor sanitation (de Wit et al. 2017). However, previous research on the impact of income disparity on animal health has had inconclusive results (Catapan 2019), and there have been no previous studies on associations between economic wealth (i.e., per capita GDP) and parasite and pathogen prevalence in wildlife. Thus, while some of these studies suggest that parasite and pathogen prevalence in wildlife varies with socioeconomic factors, others require further study, and little is known about which of these socioeconomic factors are most important. This lack of resolution, combined with use of different metrics between studies of how parasite and pathogen prevalence in humans and wildlife varies with socioeconomic factors, poses a notable challenge in understanding and integrating parasite distributional variation in humans and wildlife.

Companion animals like cats (*Felis catus*) and dogs (*Canis lupus familiaris*) are globally ubiquitous, landscape integrators that share parasites and pathogens with both humans and wildlife (Day 2011). Both animals occur globally and have co-existed with humans for thousands of years (Driscoll et al. 2007, Shannon et al. 2015). They share close contact with humans, frequently living in homes and even sleeping in peoples’ beds (Chomel and Sun 2011), occur in high densities in urban, rural, and natural areas around the world (Tiwari et al. 2019, McDonald and Skillings 2021), and commonly interact with wildlife (Medina et al. 2011, Young et al. 2011, Hughes and Macdonald 2013, Loss et al. 2013, Plaza et al. 2018). Additionally, many parasites and pathogens found in cats and dogs are well-studied due to their relevance to veterinary and human health (Bowser and Anderson 2018), and the ability of parasites to infect humans are best predicted by ability to infect companion animals (Majewska et al. 2021). Lastly, cats and dogs are both *Carnivora*, a taxa that contains among the greatest numbers of zoonotic pathogens (Keesing and Ostfeld 2021). For these reasons, companion animals are a useful model for understanding the ecology of parasites and pathogens across both environmental and socioeconomic gradients.

By evaluating associations among socioeconomic and environmental gradients and parasite and pathogen prevalence in model species that utilizes both natural and anthropogenic habitat, and has contact with both humans and wildlife, we analyze how these gradients differ in relative strengths of association with parasite and pathogen abundance at this One Health interface. In accordance with previous research on parasite abundance in wildlife (Table 1), we hypothesize that parasite distribution and abundance in companion animals is negatively correlated with biodiversity and greater on islands (i.e., island syndrome; Table 1). Furthermore, in accordance with previous research on socioeconomic factors and disease burdens in humans (Table 1), we hypothesize that parasite and pathogen prevalence will be negatively correlated with GDP and sanitation, and positively associated with income inequality (Table 1).

## Results

Parasite and pathogen prevalence in free-roaming companion animals was related to sanitation, GDP, and island habitation (Table 2, Figure 1a,1b). Parasite prevalence was best explained by two models, which were within < 2 delta AICc and thus considered equally explanatory. These two models included one with sanitation and the other containing sanitation and island (Table 3). In the full averaged model, for every 10% increase in sanitation, infectious agent prevalence in free-roaming cats and dogs decreased by 12% (5%-19% 95% C.L.; *p* = 0.002). Additionally, parasite and pathogen prevalence in free-roaming cats and dogs on islands was 12% higher than on continents, but this effect was not significant (−2%-76% 95% C.L.; *p* = 0.378).

**Table 2.** Model parameters for all considered models evaluating the effect of socioeconomic (GDP-per capita gross domestic product of country, sanitation-proportion of population with access to safely managed sanitation services, gini-gini coefficient index of income inequality of country) and environmental (biodiversity index of country, island-whether or not free-roaming cat dog was on an island, absolute latitude-based on center point of country) variables on infectious agent prevalence of free-roaming cats and dogs from compiled studies evaluated in the present work.

| Model | Parameter | Effect size | confidence limit | p value |
| --- | --- | --- | --- | --- |
| **Null** |  |  |  |  |
|  | Intercept | 0.15 | 0.12–0.20 | <0.001 |
| **sanitation** |  |  |  |  |
|  | Intercept | 0.40 | 0.19–0.85 | **0.015** |
|  | sanitation | 0.89 | 0.83–0.96 | **0.004** |
| **sanitation + island** |  |  |  |  |
|  | Intercept | 0.46 | 0.21–0.98 | **0.039** |
|  | sanitation | 0.87 | 0.83–0.96 | **0.001** |
|  | island [Y] | 1.34 | 0.95-1.90 | 0.154 |
| **GDP** |  |  |  |  |
|  | Intercept | 0.18 | 0.12–0.25 | <0.001 |
|  | GDP | 0.90 | 0.82–0.99 | **0.03** |
| **GDP + island** |  |  |  |  |
|  | Intercept | 0.18 | 0.12–0.25 | <0.001 |
|  | GDP | 0.89 | 0.80–0.98 | **0.02** |
|  | island [Y] | 1.27 | 0.90–1.80 | 0.17 |
| **biodiversity index + absolute latitude** |  |  |  |  |
|  | Intercept | 0.32 | 0.14–0.73 | **0.01** |
|  | biodiversity index | 0.54 | 0.24–1.20 | 0.12 |
|  | absolute latitude | 0.99 | 0.98–1.00 | **0.031** |
| **gini** |  |  |  |  |
|  | Intercept | 0.11 | 0.05–0.23 | <0.001 |
|  | gini | 1.01 | 0.99–1.02 | 0.36 |

**Table 3.**
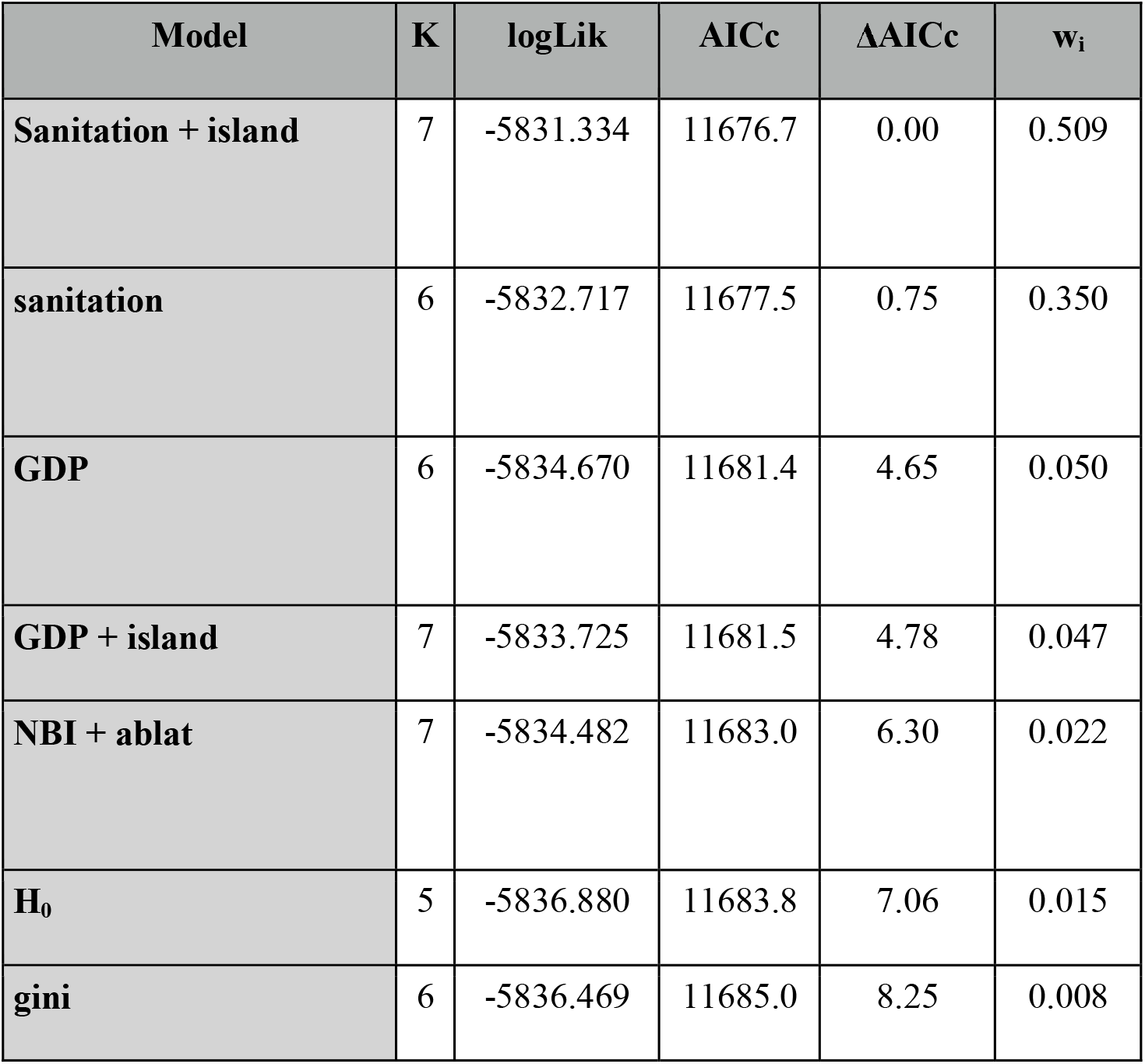
Model results of the association between socioeconomic and environmental variables and infectious agent prevalence of free-roaming cats and dogs. AICc values, delta (difference from lowest AICc score), weights, log likelihoods (logLik) and degrees of freedom (K). Sanitation is the proportion of a country’s population with access to safely managed sanitation services, island- is whether or not the study evaluating free-roaming cat/dog infectious agent prevalence was conducted on an island, GDP is the per capita gross domestic product of country, NBI is the national biodiversity index of country, ablat is the absolute latitude of center point of country, H_0_ is the null model, gini is the gini coefficient depicting income inequality of country.

**Figure 1.**
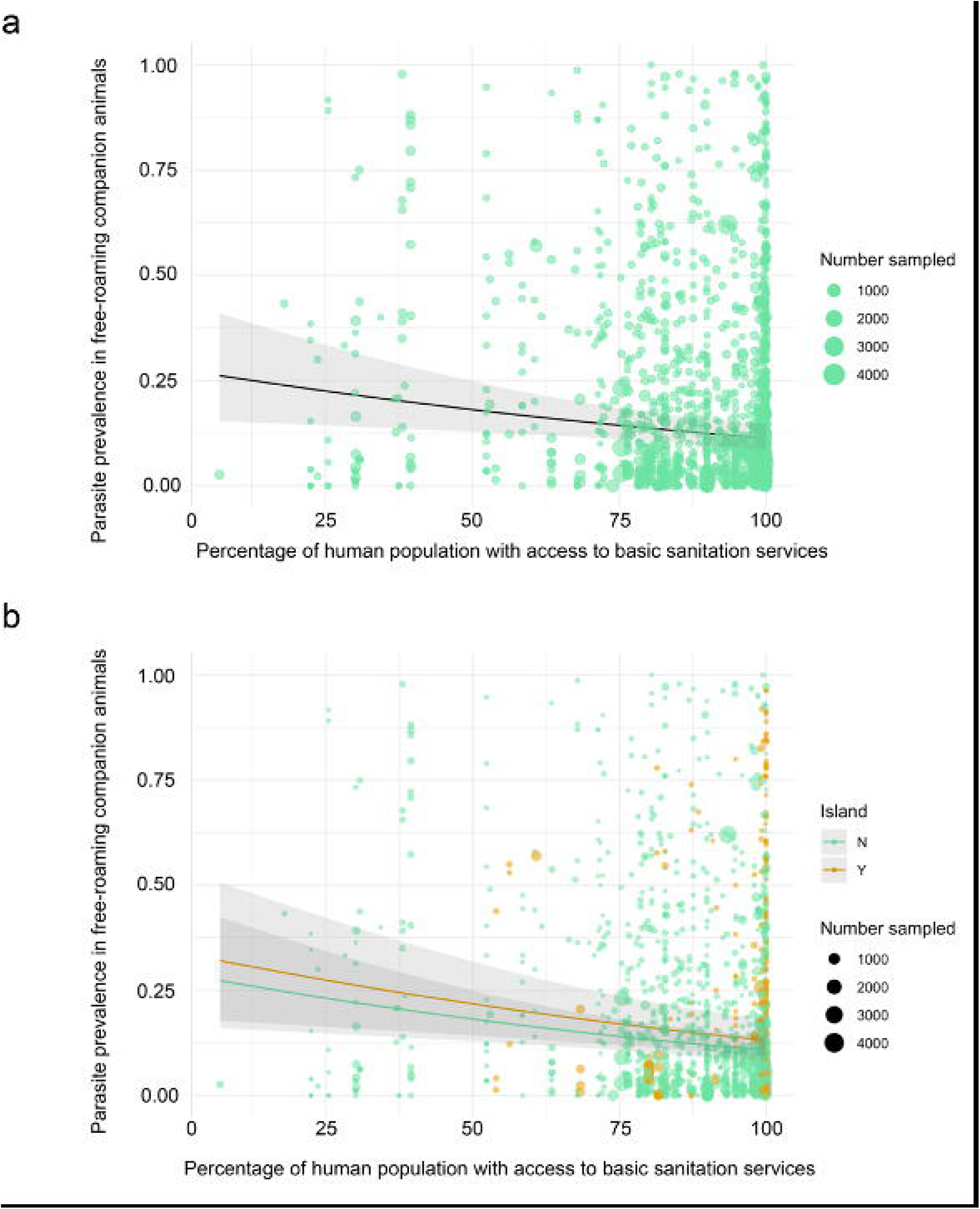
Scatter plots for the two top (ranked by AICc) multilevel GLMM with 95% confidence interval (in gray) evaluating the associations between socioeconomic and ecological variables and parasite prevalence in free-roaming cats and dogs. a) sanitation (percentage of country population with access to safely managed sanitation services); b) sanitation and island (whether or not the parasite was detected in a free-roaming cat/dog on an island).

According to the ICC results on the null model, each included random variable accounted for differing levels of heterogeneity (Supplementary Figure 1a, 1b). For example, “infectious agent detection method” accounted for the least amount of variation in the data (9.4%) while “infectious agent species” and “study” accounted for 21.8% and 18.6% of the variation, respectively. The greatest amount of variation was captured by “unique sample ID”, nested within “study” (61.4%). The results of the leave one out analysis indicated that no single study made the effect of sanitation significant on infectious agent prevalence in free-roaming cats and dogs (Supplementary Figure 2). Regression analysis revealed that sample size was significantly associated with parasite and pathogen prevalence in free-roaming companion animals (p < 0.001) such that for every 100 dogs added to the sample size of a given study, infectious agent prevalence decreased by 7.8% (4.4%-11% C.L., *p* = <0.001; Figure 2). Sample size was not significantly correlated with any of the independent variables considered in this study.

**Figure 2.**
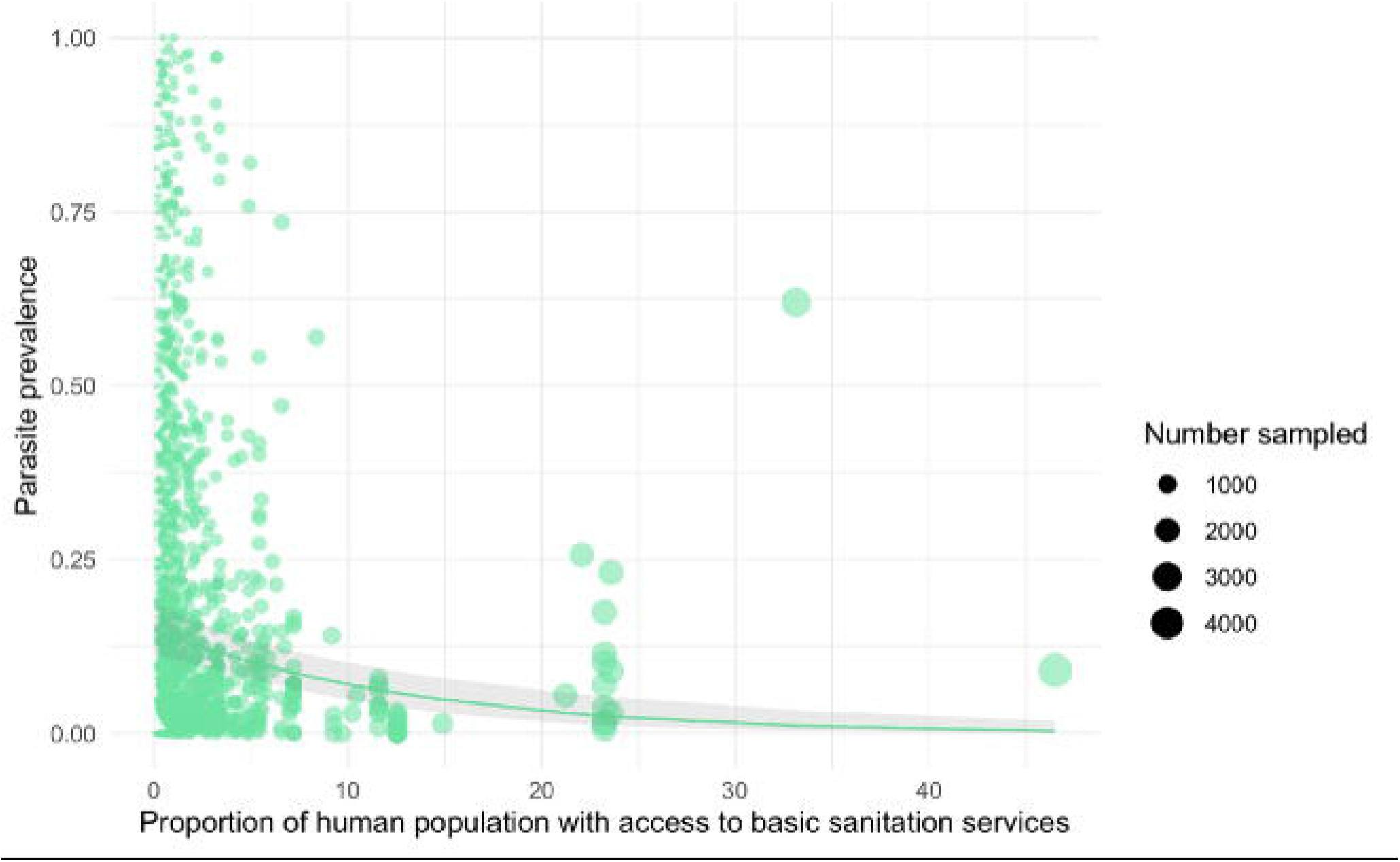
Scatter plot with multilevel GLMM trendline with 95% confidence interval (gray) demonstrating the relationship between sample size (number of free-roaming cats/dogs sampled in each study for each parasite) and infectious agent prevalence.

## Discussion

Companion animals live across a selection of environmental gradients, and thus can be used as efficient models to characterize parasite prevalence along socioeconomic and environmental gradients as well as serve as proxies for parasite and pathogen prevalence in humans and wildlife. We found support for our hypothesis that sanitation was negatively correlated with parasite and pathogen prevalence, and partial support that parasite and pathogen prevalence was greater on islands. These were the only parameters in the top models, and thus may be most important to consider in terms of factors affecting parasite prevalence in these hosts. GDP was also a significant socioeconomic variable associated with parasite and pathogen prevalence, but redundance with sanitation suggests that it is more weakly associated with parasite prevalence than the latter.

Sanitation being in the top two models suggests that socioeconomic factors may play a larger role in companion animal health than previously appreciated. Better sanitation is known to be a cost-effective intervention towards improving overall human health (Haller et al. 2007) and has been the focus of much attention at reducing disease burdens in humans around the world (World Health Organization 2019, Nelson et al. 2021). However, the effect of these interventions on animals, or human exposure to animal feces, have not been well studied (Prendergast et al. 2019), but a couple investigations on specific parasites or geographic locations have indicated that improved sanitation is associated with reduced parasite prevalence in wildlife (de Wit et al. 2017, Murray et al. 2020). Importantly, this reduction in parasite/pathogen prevalence of companion animals is not inconsequential for human health. In fact, animal feces may be an important health consequence of poor sanitation infrastructure in general (Berendes et al. 2018), where infected animals can shed contaminated feces into the environment and are thereby capable of infecting humans. Companion animals have strong associations with both humans (Chomel and Sun 2011, Zhang et al. 2019) and wildlife (Medina et al. 2011, Young et al. 2011, Hughes and Macdonald 2013, Loss et al. 2013, Plaza et al. 2018), making them an important bridge host (i.e., a link that transmits pathogens between a reservoir host population and some target host population; Caron et al. 2015) for many diseases between humans and wildlife (Ellwanger and Chies 2019). Parasite and pathogen vectoring from companion animals to wildlife has also received much attention, and many species of conservation concern are at risk for health problems from parasites and pathogens of companion animals (Salb et al. 2008, Day 2011, Wilson et al. 2021). Thus, in addition to reducing exposure of parasites and pathogens to humans, improving sanitation could be a sound conservation strategy for reducing exposure of wildlife to infectious agents in the environment. In fact, one sanitation intervention reported reduced ruminant fecal contamination in the soil, implying improved sanitation access can reduce environmental fecal contamination from animals as well as humans (Boehm et al. 2016). Taken together, improved sanitation is a One Health intervention worthy of further exploration.

Parasite and pathogen prevalence on islands was higher than that of continents. While not significantly associated with parasite and pathogen prevalence, island habitation was included in one of the two top models, and thus may be important in predicting infectious agent prevalence in free-roaming companion animals. Increased parasite and pathogen prevalence on islands is consistent with the island syndrome hypothesis (Dybing et al. 2018). Although island habitation in our analysis was not significantly associated with parasite and pathogen prevalence, previous studies at smaller scales did find significant associations, with either positive and negative effects (Beadell et al. 2007, Pérez-Rodríguez et al. 2013, Literák et al. 2015, Loiseau et al. 2017, Dybing et al. 2018). Higher parasite and pathogen prevalence on islands is of conservation concern since a large proportion of island species are endemics (Kier et al. 2009) that can be particularly susceptible to introduced parasites and pathogens due to reduced immunological investments (Lindström et al. 2003).

Biodiversity was unrelated to parasite and pathogen prevalence in free-roaming cats and dogs, suggesting it is less predictive of parasite and pathogen prevalence than sanitation or island habitation. These results corroborate previous findings that biodiversity is less influential at larger scales (Rohr et al. 2020), perhaps due to stronger emergent factors at global scales (Johnson et al. 2015, Rohr et al. 2020), such as climate (Rohr et al. 2020). Recent findings also indicate that biodiversity loss, rather than biodiversity itself, may be a stronger predictor of parasite and pathogen prevalence (Halliday et al. 2020).

With respect to our findings that prevalence was significantly correlated with sample size, the latter was not correlated with any of the independent variables we evaluated, and thus do not believe this relationship affected our results. The relationship between prevalence and sample size is still notable since it may reflect a lower likelihood of publishing studies with small sample sizes that find a low prevalence of the infectious agent of interest. Such publication bias can be harmful to our greater knowledge of diseases in companion animals and efforts ought to be made to publish such studies regardless of prevalence findings, given the lack of health surveillance for companion animals (Radford et al. 2021).

Parasites and pathogens carried by free-roaming companion animals are associated with both socioeconomic and environmental gradients, particularly with sanitation and island habitation. As such, future research should target how these associations may be altered by aspects of parasite ecology. For example, are some of these associations more important for parasites and pathogens with different transmission types? While sanitation interventions typically target parasites spread through the environment (i.e., soil-borne), improved sanitation in the form of refuse management may decrease the prevalence of parasites and pathogens that are directly transmitted, due to reduced aggregation of animal hosts around resources (Becker and Hall 2014, Becker et al. 2015). Furthermore, previous research indicates that urbanization gradients may reduce parasite and pathogen prevalence, but many competing hypotheses exist and results across host and parasite taxa have been inconsistent (Bradley and Altizer 2007, Werner and Nunn 2020). In addition to these potential avenues for future research and given the breadth of parasites and pathogens infecting free-roaming cats and dogs (including many of which that also infect humans and wildlife (Salb et al. 2008, Day 2011), stronger management of free-roaming cat and dog populations, especially on islands and in regions with poor sanitation infrastructure, may be a particularly important One Health intervention.

## Materials and Methods

### Systematic Review

To build the database of studies pertaining to infectious agent prevalence in feral cats and dogs, we conducted two searches in Web of Science Core Collection, following PRISMA guidelines (Page et al. 2021; Figure 3). The first search was conducted on January 11^th^ 2018 and returned 500 results from the following search terms: TOPIC: (‘feral cat’ OR ‘feral dog*’) AND (‘infect*’ OR ‘parasit*’ OR ‘disease*’ OR ‘virus*’) NOT TITLE: (review). The second search, done on December 23^rd^, 2018, returned 1,308 results from the search terms: TOPIC: (feral OR stray OR free-roaming OR ‘free roaming’ OR free-ranging OR ‘free ranging’) AND TOPIC: (dog OR canis familiaris) AND TOPIC: (infect* OR parasit* OR disease* OR pathogen*) NOT TITLE: (review).

**Figure 3.**
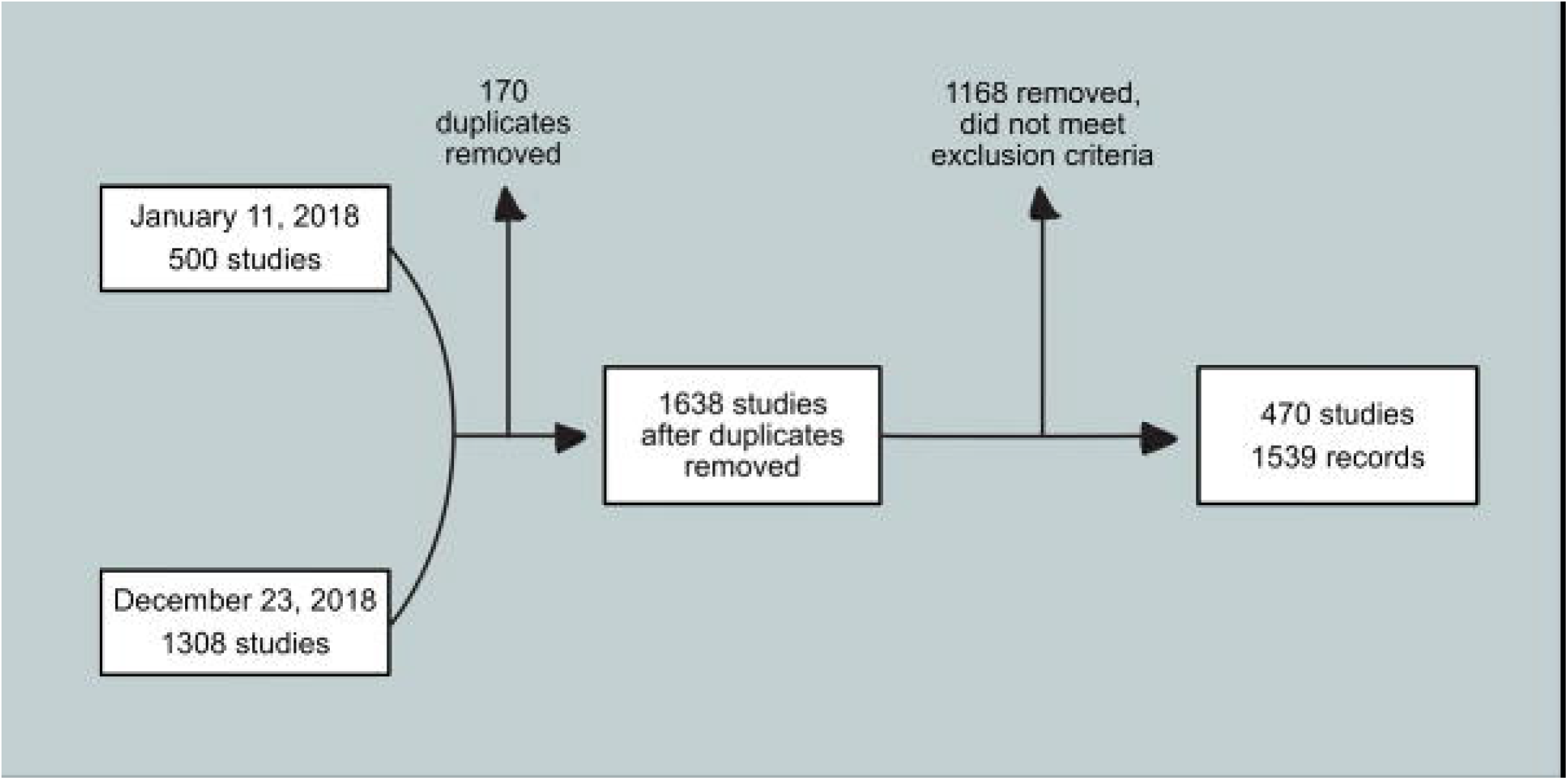
PRISMA (Page et al. 2021) flow-chart detailing the systematic review conducted to identify studies investigating prevalence in free-roaming cats and dogs. Two searches were conducted on 11 January 2018 and 23 December 2018 each returning 500 and 1,308 studies, respectively. After 170 duplicates were removed, 1,638 studies remained. Studies were then examined manually and 1,168 studies that did not meet the exclusion criteria were removed. This process resulted in a database of 470 studies with 1,539 records (because some studies investigated multiple parasites or conducted research in multiple regions).

We removed 170 duplicates from the 1,808 identified between the two searches, resulting in a total of 1,638 studies. We then manually sorted the studies, and 1,168 that did not meet the following exclusion criteria were removed: unclear parasite sampling method; infection prevalence of specific parasite not given (only overall prevalence); animal described as “semiferal” without further description of lifestyle; prevalence values unclear (only gave confidence interval); feces collected from ground or litter tray; free-roaming designation included “shelter animals” but not clear if this designation only included previously free-roaming animals (or if they included owner drop-offs); seroprevalence values duplicated from another study; no information provided on how feces were collected. This filtering resulted in 470 studies with 1,539 effect sizes, due to single studies detecting multiple parasites and pathogens or being conducted across multiple regions (Figure 4), used in the present analysis (Supplementary Figure 3). We also removed records from countries for which we were unable to find supplemental data, resulting in a database of 445 studies with 1,480 records. For the final database, we recorded a number of additional moderating factors from each study: species (cat or dog sampled), parasite species or virus name; tissue where parasite detected; parasite detection method; total number of animals sampled in the study; number of positive infections; country where study was conducted; whether the study was conducted on an island; parasite type (i.e., helminth, bacteria, protozoan, virus); and whether or not the parasite was zoonotic. Lastly, we assigned a unique identification number to each record.

**Figure 4.**
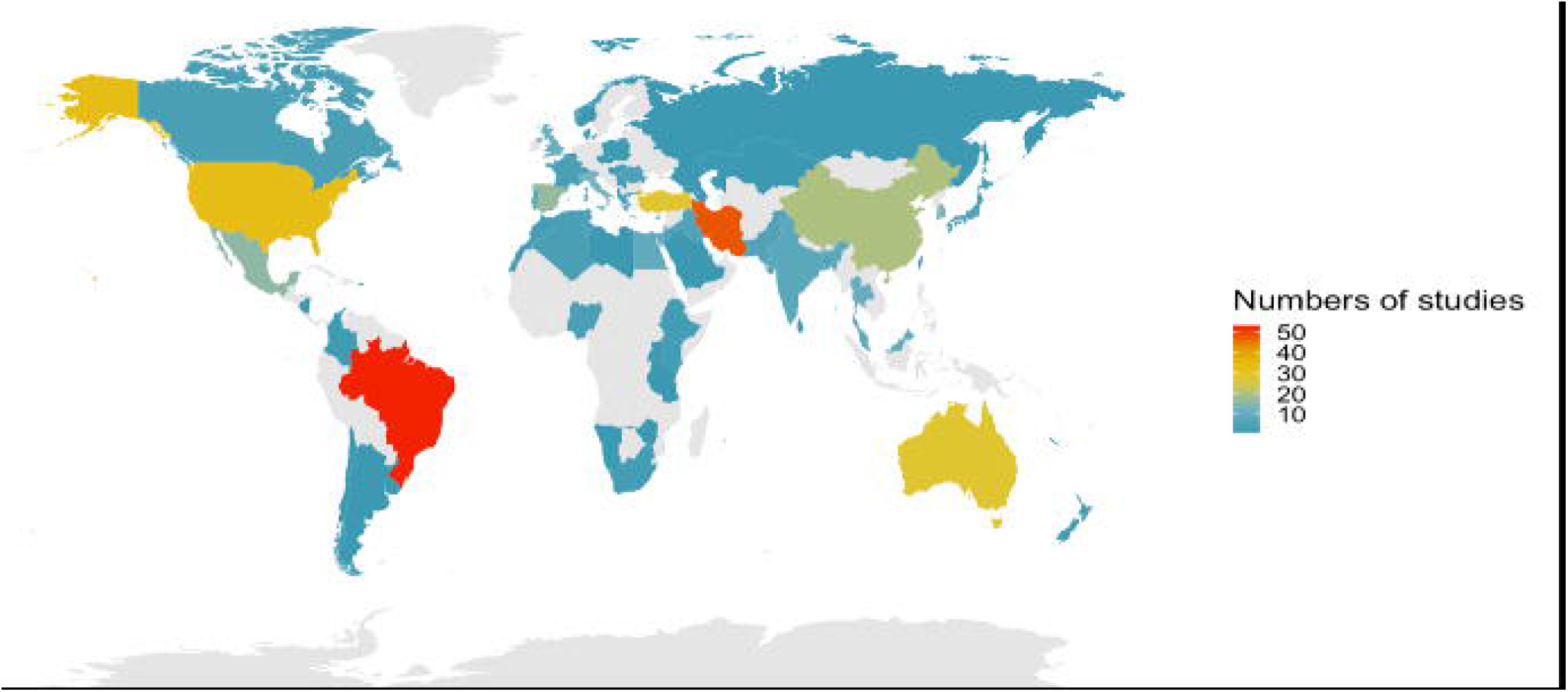
Heat map depicting numbers of studies by country compiled to analyze socioeconomic, and environmental variables influencing parasite prevalence in free-roaming cats and dogs.

### Environmental and Socioeconomic Data

The dependent variable across all models was the prevalence of each infectious agent in each study (number infected/total sampled). We used the following sources for country-level ecological and socioeconomic data (see Supplementary Figure 4): per capita gross domestic product in U.S. Dollars (GDP; Lindgren 2019); income inequality score (gini index; World Bank 2020a); national biodiversity index (NBI; Convention on Biological Diversity 2011); % of population using at least basic sanitation services (hereafter referred to as % sanitation; World Bank 2020b); and absolute latitude (the distance from the equator, obtained using the center-point latitude of the country). We linked these databases to the free-roaming companion animal parasite and pathogen database using R version 4.0.4 and R packages dplyr, tidyverse, Hmisc, rgeos, and rworldmap (South 2011, Wickham et al. 2019, Bivand and Rundel 2020, Harrell Jr. and Dupont 2021, Wickham et al. 2021). We then removed samples that did not have associated values for GDP, gini, sanitation, NBI, or latitude given that model comparison requires use of the same dataset. Summaries and mapping of numbers of studies as well as totaled prevalence across studies was conducted using the R packages dplyr and ggplot2 (Wickham 2019, Wickham et al. 2021).

### Model Selection

In order to develop our list of candidate models, we conducted exploratory moderator analyses with the goal of identifying which of the included variables may be colinear, redundant, or confounding in effect on infectious agent prevalence. To test for collinearity, we calculated variance inflation score (VIF) for each of the variables. Typically, if VIF > 5, high collinearity is likely (Craney and Surles 2007). In this exploratory analysis, the VIF for all variables was < 2.

We identified redundant variables using a Pearson correlation for comparison between continuous variables (all variables except island) and polyserial correlation between continuous and categorical variables (island with all other variables); conducted using the polycor package (Fox 2019). Variables with polyserial scores between +/- 0.50 and +/- 1, +/- 0.30 and +/- 0.49 and below +/- 0.30 were classified as highly, moderately and lowly correlated (Olsson and Drasgow 1982), respectively. Following this polyserial correlation, all variables with moderate to high correlation with each other were tested in a series of weighted multilevel logit generalized linear mixed models to identify redundant variables. Redundant variables were characterized as those that changed considerably when another correlated variable was added to the model (Supplementary Figure 5) and are therefore less explanatory. We identified confounding variables where both changed considerably between univariable and multivariable models. Variables found to be redundant with another were not included in multivariable models together in the final list of models (Table 1).

The calculated VIF score was < 2 for all independent variables, indicating no collinearity. There were several variables, however, with moderate to high correlation according to polyserial or Pearson correlations (Supplementary Figure 5). Of the correlated variables, we identified several instances of redundancy or confounding which resulted in our considered models (Supplementary Figure 5, Table 1).

### Bias Estimation

A regression test was carried out to identify bias between sample size and prevalence in each study. We conducted this regression using sample size in each study as the independent variable and prevalence as the dependent variable in a weighted, nested, univariable multilevel logit generalized mixed linear model (GLMM). We then performed a series of polyserial and Pearson correlation tests (Olsson and Dragow 1982) between sample size and the other independent variables to identify potential confounding or redundancy between these variables with sample size. We also conducted a leave one out analysis on the univariable GLMM containing sanitation to determine if the results of any one study were overly influencing our results, because this was the simpler of the top two models according to the AICc results (Hurvich and Tsai 1989; Supplementary Figure 2).

### Statistical Analysis

We modeled infectious agent prevalence as a function of latitude, island, biodiversity index, per capita GDP, gini coefficient, and sanitation using a series of weighted, nested univariable multilevel logit GLMMs using the lme4 package (Bates et al. 2015) in R. Study (to account for variation across studies) and a unique identification number (each parasite sampled within each study was given a unique id) were both used as nested random variables to account for the structure of the data (because some studies sampled multiple parasites and pathogens). We also included parasite and pathogen species and parasite and pathogen detection method (i.e., via necropsy, microscopy, polymerase chain reaction) as random variables to account for additional heterogeneity. We then compared these models using the corrected Akaike’s Information Criterion (AICc) using the MuMin package (Bartoń 2020). The model with the lowest AICc score was considered the best model for explaining parasite prevalence in free-roaming cats and dogs. We considered models with delta AICc <2 equally explanatory, and used the full average of these models to get the final best explanatory model. Variables within each top model with a p-value < 0.05 were considered significant. Lastly, we calculated the intra-class correlation coefficients (ICC; Hofmann 1997) for the null model to characterize the level of heterogeneity captured by each random variable.

## Acknowledgments

We thank Dr. Wolfgang Viechtbauer and Dr. Todd Steury for guidance on our statistical analysis, and Dr. Patricia Hartman for guidance in conducting the systematic literature search.

