## Supplementary Figure 1 for "Parasite and pathogen prevalence in our closest animal companions is determined by accessibility of sanitation services"

**a)** Intra-class correlation coefficients (ICC) for random effects of null model from analysis evaluating the effect of socioeconomic and environmental variables on infectious agent prevalence of free-roaming cats and dogs from compiled studies evaluated in the present work. Group- random effect; pathogen- pathogen species; method- detection method used to identify pathogen in prevalence study; study- unique identifier for each study; study: unique- each unique pathogen prevalence value nested within each study (because some studies evaluated multiple pathogens).

| **Group** | **ICC** |
| --- | --- |
| study:uniq | 0.614 |
| study | 0.186 |
| pathogen | 0.218 |
| method | 0.094 |

**b)** Caterpillar plots depicting random intercepts (blue dots) and conditional standard deviations (error bars) for each of the random variables included in the analysis evaluating the effect of socioeconomic and environmental variables on infectious agent prevalence of free-roaming cats and dogs from compiled studies evaluated in the present work. Method- detection method used to identify pathogen in prevalence study; study- unique identifier for each study; study: unique- each unique pathogen prevalence value nested within each study (because some studies evaluated multiple pathogens); Country- country where the study was conducted.

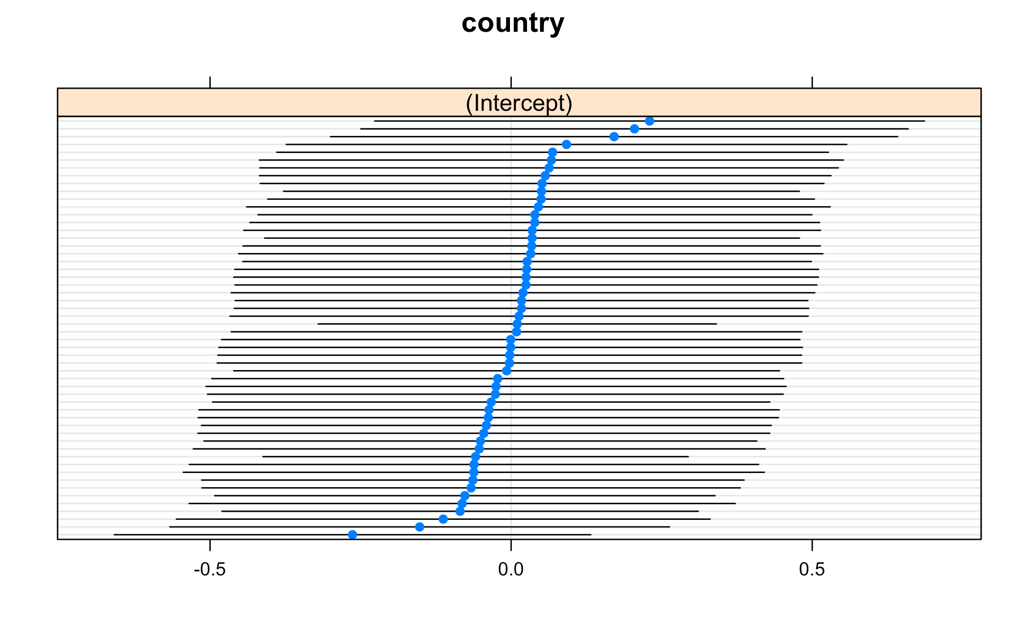

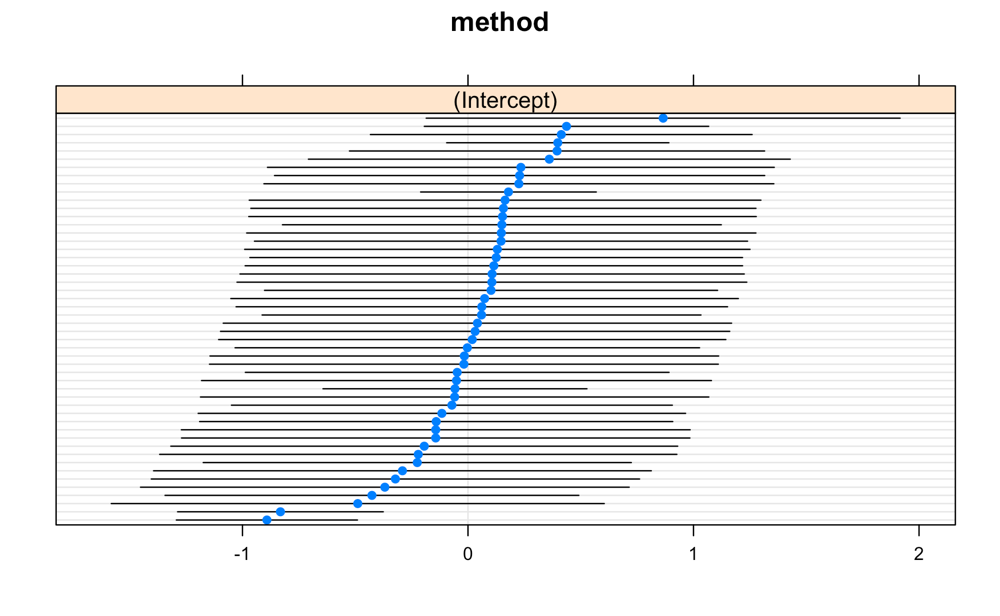

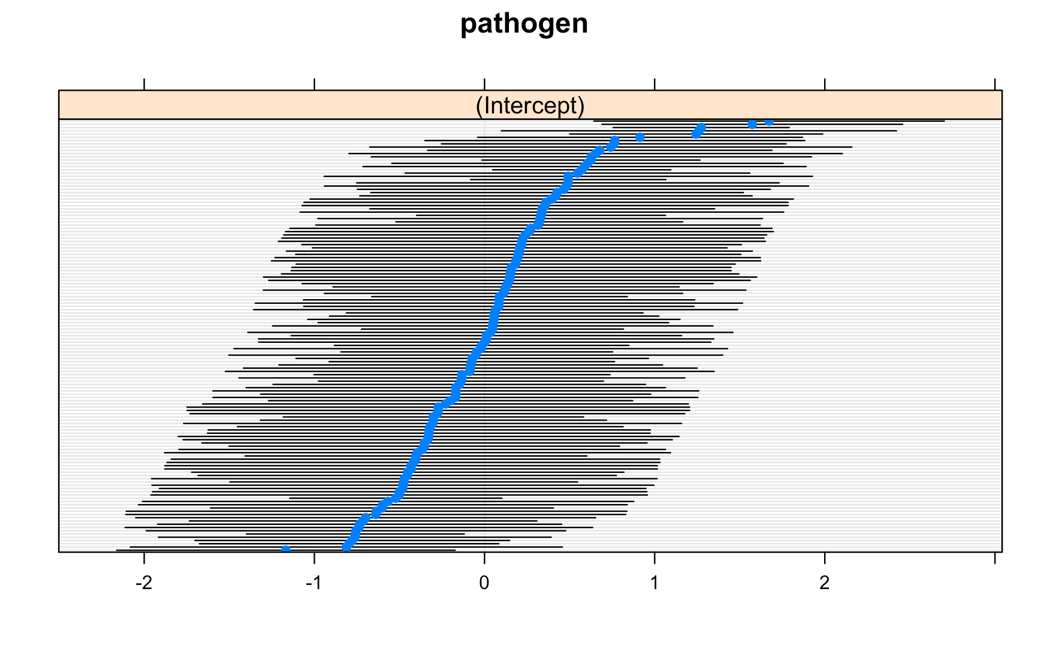

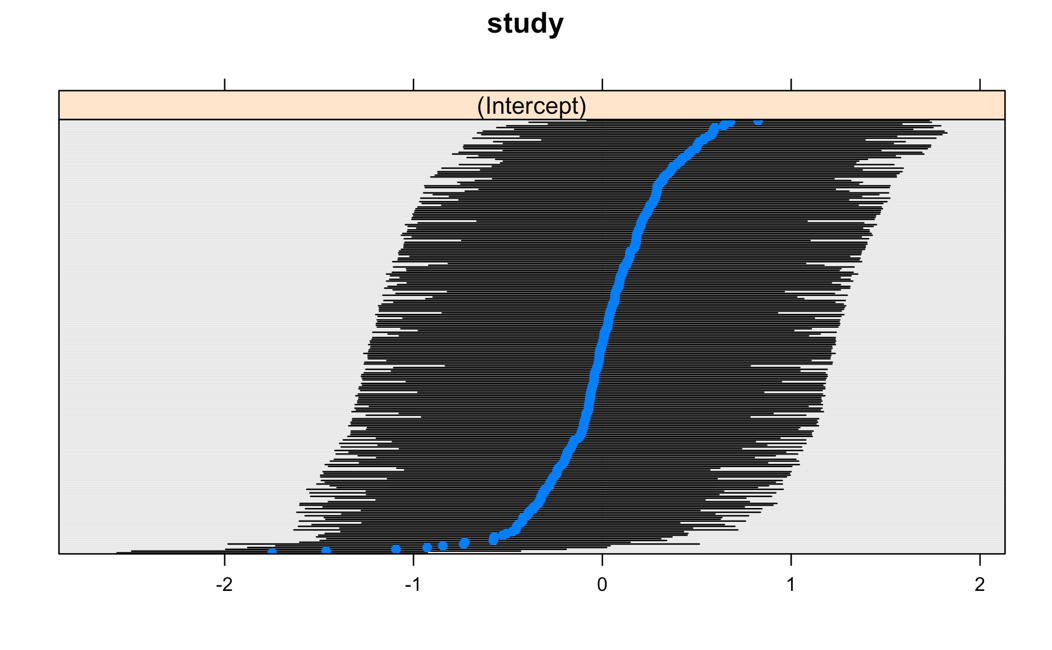

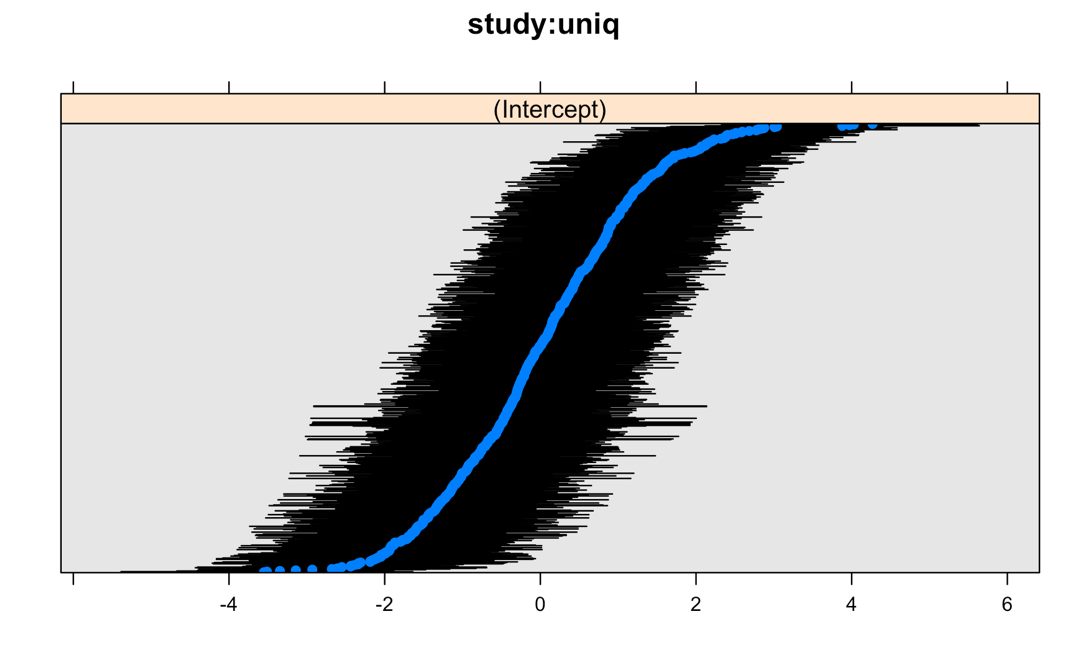
