## Supplementary Figure 3 for "Parasite and pathogen prevalence in our closest animal companions is determined by accessibility of sanitation services"

**Supplementary Figure 3:** List of studies included in the database that evaluate prevalence of parasites in free-roaming cats (*Felis catus*) or dogs (*Canis lupus familiaris*)

el-Ahraf A WBW Tacal JV, Sobih M, Amin M, Lawrence W. 1991. Prevalence of cryptosporidiosis in dogs and human beings in San Bernardino County, California. J Am Vet Med Assoc. 198(4):631–634.

Abbaszadeh Afshar MK, Sharifi II, Bamorovat M, Mohebali M, Bahreini MS, Naderi A. 2018. Canine visceral leishmaniasis; a seroepidemiological survey in Jiroft district, southern Kerman province, Southeastern Iran in 2015. Iran J Parasitol. 13(1):67–71.

Abdi J, Taherikalani M, Asadolahi K, Emaneini M. 2013. Echinococcosis/Hydatidosis in Ilam Province, Western Iran. Iran J Parasitol. 8(3):417–422.

Abdulsalam J. 1986. Intestinal helminth-parasites of stray dogs in Kuwait. Arab Gulf J Sci Res. 4(2):659–663.

Abu-Madi MA, Behnke JM. 2014. Feline patent *Toxoplasma*-like coccidiosis among feral cats (Felis catus) in Doha city, Qatar and its immediate surroundings. Acta Parasitol. 59(3):390–397. doi:10.2478/s11686-014-0254-y.

Abu-Madi MA, Behnke JM, Prabhaker KS, Al-Ibrahim R, Lewis JW. 2010. Intestinal helminths of feral cat populations from urban and suburban districts of Qatar. Vet Parasitol. 168(3–4):284–292. doi:10.1016/j.vetpar.2009.11.027.

Abu-Madi MA, Pal P, Al-Thani A, Lewis JW. 2008. Descriptive epidemiology of intestinal helminth parasites from stray cat populations in Qatar. J Helminthol. 82(1):59–68. doi:10.1017/S0022149X07870830.

Adams PJ, Elliot AD, Algar D, Brazell RI. 2008. Gastrointestinal parasites of feral cats from Christmas Island. Aust Vet J. 86(1–2):60–63. doi:10.1111/j.1751-0813.2007.00246.x.

Adanir R, Sezer K, Köse O. 2013. The prevalence of *Dirofilaria immitis* in dogs with different breed, ages and sex. Vet Fak Derg. 60:241–244. doi:10.1501/Vetfak_0000002586.

Adesiyun AA, Hull-Jackson C, Mootoo N, Halsall S, Bennett R, Clarke NR, Whittington CU, Seepersadsingh N. 2006. Sero-epidemiology of canine leptospirosis in Trinidad: serovars, implications for vaccination and public health. J Vet Med B Infect Dis Vet Public Health. 53(2):91–99. doi:[10.1111/j.1439-0450.2006.00922.x](https://doi.org/10.1111/j.1439-0450.2006.00922.x).

Adinezadeh A, Kia EB, Mohebali M, Shojaee S, Rokni MB, Zarei Z, Mowlavi G. 2013. Endoparasites of Stray Dogs in Mashhad, Khorasan Razavi Province, Northeast Iran with Special Reference to Zoonotic Parasites. Iran J Parasitol. 8(3):459–466.

Afonso E, Thulliez P, Gilot-Fromont E. 2006. Transmission of *Toxoplasma gondii* in an urban population of domestic cats (*Felis catus*). Int J Parasitol. 36(13):1373–1382. doi:[10.1016/j.ijpara.2006.07.010](https://doi.org/10.1016/j.ijpara.2006.07.010).

Afonso E, Thulliez P, Pontier D, Gilot-Fromont E. 2007. Toxoplasmosis in prey species and consequences for prevalence in feral cats: not all prey species are equal. Parasitology. 134(Pt.14):1963–1971. doi:[10.1017/S0031182007003320](https://doi.org/10.1017/S0031182007003320).

Ahmad N, Barazandeh M. 2010. A Survey on *Dirofilaria immitis* Occurence in Stray Dogs of Tabriz (Iran). Acta Veterinaria Brno. 79:449–451. doi:[10.2754/avb201079030449](https://doi.org/10.2754/avb201079030449).

Ahmad W, Mahmood F, Li Y, Duan M, Guan Z, Zhang M, Ali M, Liu Z. 2016. Immunopathological studies of canine rabies in Faisalabad, Pakistan. J Anim Plant Sci. 26:1–20.

Akhtardanesh B, Sharifi I, Mohammadi A, Mostafavi M, Hakimmipour M, Pourafshar NG. 2017. Feline visceral leishmaniasis in Kerman, southeast of Iran: Serological and molecular study. J Vector Borne Dis. 54(1):96–102.

Aktas M, Dumanlı N, Özübek S S, Altay K K, Balkaya I, Utuk AE AE, Kırbas A, Şimsek S. 2015. A molecular and parasitological survey of *Hepatozoon canis* in domestic dogs in Turkey. Vet Parasitol. 209(3–4):264–267. doi:[10.1016/j.vetpar.2015.02.015](https://doi.org/10.1016/j.vetpar.2015.02.015).

Aktas M, Ozubek S. 2018. A molecular survey of hemoplasmas in domestic dogs from Turkey. Vet Microbiol. 221:94–97. doi:[10.1016/j.vetmic.2018.06.004](https://doi.org/10.1016/j.vetmic.2018.06.004).

Aktas M, Ozübek S, Ipek DNS. 2013. Molecular investigations of Hepatozoon species in dogs and developmental stages of *Rhipicephalus sanguineus*. Parasitol Res. 112(6):2381–2385. doi:[10.1007/s00436-013-3403-6](https://doi.org/10.1007/s00436-013-3403-6).

Akter S, Alam MZ, Nakao R, Yasin G, Kato H, Katakura K. 2016. Molecular and Serological Evidence of *Leishmania* Infection in Stray Dogs from Visceral Leishmaniasis–Endemic Areas of Bangladesh. Am J Trop Med Hyg. 95(4):795–799. doi:[10.4269/ajtmh.16-0151](https://doi.org/10.4269/ajtmh.16-0151).

Akucewich LH GEC Philman K, Clark A, Gillespie J, Kunkle G, Nicklin CF. 2002. Prevalence of ectoparasites in a population of feral cats from north central Florida during the summer. Vet Parasitol. 109(1–2):129–139. doi:[10.1016/s0304-4017(02)00205-4](https://doi.org/10.1016/s0304-4017(02)00205-4).

Alam MA, Maqbool A, Nazir MM, Lateef M, Khan MS, Lindsay DS. 2015. *Entamoeba* infections in different populations of dogs in an endemic area of Lahore, Pakistan. Vet Parasitol. 207(3–4):216–219. doi:[10.1016/j.vetpar.2014.12.001](https://doi.org/10.1016/j.vetpar.2014.12.001).

Alanazi AD. 2018. Parasitological and Molecular Detection of Canine Trypanosomiasis From Riyadh Province, Saudi Arabia. J Parasitol. 104(5):539–543. doi:[10.1645/18-16](https://doi.org/10.1645/18-16).

Ali CN, Harris JA, Watkins JD, Adesiyun AA. 2003. Seroepidemiology of *Toxoplasma gondii* in dogs in Trinidad and Tobago. Vet Parasitol. 113(3):179–187. doi:[10.1016/S0304-4017(03)00075-X](https://doi.org/10.1016/S0304-4017(03)00075-X).

Al-Kappany YM, Lappin MR, Kwok OCH, Abu-Elwafa SA, Hilali M, Dubey JP. 2011. Seroprevalence of *Toxoplasma gondii* and concurrent *Bartonella* spp., feline immunodeficiency virus, feline leukemia virus, and *Dirofilaria immitis* infections in Egyptian cats. J Parasitol. 97(2):256–258. doi:[10.1645/GE-2654.1](https://doi.org/10.1645/GE-2654.1).

Al-Kappany YM, Rajendran C, Ferreira LR, Kwok OCH, Abu-Elwafa SA, Hilali M, Dubey JP. 2010. High prevalence of toxoplasmosis in cats from Egypt: isolation of viable *Toxoplasma gondii*, tissue distribution, and isolate designation. J Parasitol. 96(6):1115–1118. doi:[10.1645/GE-2554.1](https://doi.org/10.1645/GE-2554.1).

de Almeida Curi NH, Araújo AS, Campos FS, Lobato ZIP, Gennari SM, Marvulo MFV, Silva JCR, Talamoni SA. 2010. Wild canids, domestic dogs and their pathogens in Southeast Brazil: disease threats for canid conservation. Biodivers Conserv. 19(12):3513–3524. doi:[10.1007/s10531-010-9911-0](https://doi.org/10.1007/s10531-010-9911-0).

Al-Qaoud KM, Abdel-Hafez SK, Craig PS. 2003. Canine echinococcosis in northern Jordan: increased prevalence and dominance of sheep/dog strain. Parasitol Res. 90(3):187–191. doi:[10.1007/s00436-002-0793-2](https://doi.org/10.1007/s00436-002-0793-2).

Amoli A, Razmi G, Khoshnegah J. 2012. A preliminary parasitological survey of *Hepatozoon* spp. Infection in dogs in Mashhad, Iran. Iran J Parasitol. 7(4):99–103.

Amouei A, Jahandar H, Daryani A, Sharif M, Sarvi S, Mizani A, Hosseini SA, Sarafrazi M, Siyadatpanah A, Gohardieh S, et al. 2018. Carnivores as important reservoirs of intestinal helminthic infections in Mazandaran Province, Northern Iran. Iran J Parasitol. 13(2):251–257.

An D-J, Jeoung H-Y, Jeong W, Park J-Y, Lee M-H, Park B-K. 2011. Prevalence of Korean cats with natural feline coronavirus infections. Virol J. 8(1):455. doi:[10.1186/1743-422X-8-455](https://doi.org/10.1186/1743-422X-8-455).

Anderson TC, Foster GW, Forrester DJ. 2003. Hookworms of feral cats in Florida. Vet Parasitol. 115(1):19–24. doi:[10.1016/s0304-4017(03)00162-6](https://doi.org/10.1016/s0304-4017(03)00162-6).

Araujo IC, Mota SB, de Aquino MHC, Ferreira AMR. 2010. *Helicobacter* species detection and histopathological changes in stray cats from Niterói, Brazil. J Feline Med Surg. 12(6):509–511. doi:[10.1016/j.jfms.2010.01.008](https://doi.org/10.1016/j.jfms.2010.01.008).

Arbabi M, Hooshyar H. 2006. Survey of echinococcosis and hydatidosis in Kashan region, Central Iran. Iranian J Publ Health. 35:75–81.

Asgarali Z, Pargass I, Adam J, Mutani A, Ezeokoli C. 2012. Haematological parameters in stray dogs seropositive and seronegative to Ehrlichia canis in North Trinidad. Ticks Tick Borne Dis. 3(4):207–211. doi:[10.1016/j.ttbdis.2012.03.006](https://doi.org/10.1016/j.ttbdis.2012.03.006).

Asl A, Jamshidi, Mohammadi M, Soroush Barhaghi M, Bahadori A. 2009. Assessment of antimicrobial resistance of cultivable Helicobacter-like organisms in asymptomatic dogs. Iran J Vet Res. 10(3): 241-249.

Asl AS, Amanizad H, Partovi A, Jalili M, Nezhad DM, Soroush M, Barzegari A, Babazadeh D. 2014. Assessment of Gastric *Helicobacter* spp. in Fresh Gastric Samples of Naturally Infected Dogs by Scanning Electron Microscopy. Kafkas Univ Vet Fak Derg 20(5): 663-669. doi:[10.9775/KVFD.2014.10778](https://doi.org/10.9775/KVFD.2014.10778).

Asl AS, Jamshidi S, Mohammadi M, Soroush MH, Bahadori A, Oghalaie A. 2010. Detection of atypical cultivable canine gastric *Helicobacter* strain and its biochemical and morphological characters in naturally infected dogs. Zoonoses Public Health. 57(4):244–248. doi:[10.1111/j.1863-2378.2008.01219.x](https://doi.org/10.1111/j.1863-2378.2008.01219.x).

Aslantaş O, Ozdemir V, Kiliç S, Babür C. 2005. Seroepidemiology of leptospirosis, toxoplasmosis, and leishmaniosis among dogs in Ankara, Turkey. Vet Parasitol. 129(3–4):187–191. doi:[10.1016/j.vetpar.2004.11.037](https://doi.org/10.1016/j.vetpar.2004.11.037).

Ataş A, Altay K, Alim A, Özkan E. 2018. Survey of *Dirofilaria immitis* in dogs from Sivas Province in the Central Anatolia Region of Turkey. Turkish J Vet Anim. 42:130-134. doi:[10.3906/VET-1707-93](https://doi.org/10.3906/VET-1707-93).

Atasevan VS, Ucar H, Akca Y. 2005. Canine coronavirus antibodies in stray dogs. Indian Vet J. 82:782–783.

Atasoy A, Paşa S, Toz SÖ, Ertabaklar H. 2010. Seroprevalence of canine visceral leishmaniasis around the Aegean Coast of Turkey. Kafkas Univ Vet Fak Derg. 16(1):1-6.

Attipa C, Papasouliotis K, Solano-Gallego L, Baneth G, Nachum-Biala Y, Sarvani E, Knowles TG, Mengi S, Morris D, Helps C, et al. 2017. Prevalence study and risk factor analysis of selected bacterial, protozoal and viral, including vector-borne, pathogens in cats from Cyprus. Parasit Vectors. 10: 130. doi:[10.1186/s13071-017-2063-2](https://doi.org/10.1186/s13071-017-2063-2).

Awadallah MAI, Salem LMA. 2015. Zoonotic enteric parasites transmitted from dogs in Egypt with special concern to *Toxocara canis* infection. Vet World. 8(8):946–957. doi:[10.14202/vetworld.2015.946-957](https://doi.org/10.14202/vetworld.2015.946-957).

Aydin MF, Sevinc F, Sevinc M. 2015. Molecular detection and characterization of *Hepatozoon* spp. in dogs from the central part of Turkey. Ticks Tick Borne Dis. 6(3):388–392. doi:[10.1016/j.ttbdis.2015.03.004](https://doi.org/10.1016/j.ttbdis.2015.03.004).

Azzag N BHJ Petit E, Gandoin C, Bouillin C, Ghalmi F, Haddad N. 2015. Prevalence of select vector-borne pathogens in stray and client-owned dogs from Algiers. Comp Immunol Microbiol Infect Dis. 38:1–7. doi:[10.1016/j.cimid.2015.01.001](https://doi.org/10.1016/j.cimid.2015.01.001).

Babuadze G, Alvar J, Argaw D, de Koning HP, Iosava M, Kekelidze M, Tsertsvadze N, Tsereteli D, Chakhunashvili G, Mamatsashvili T, et al. 2014. Epidemiology of Visceral Leishmaniasis in Georgia. PLoS Negl Trop Dis. 8(3):e2725. doi:[10.1371/journal.pntd.0002725](https://doi.org/10.1371/journal.pntd.0002725).

Bai Y, Kosoy MY, Boonmar S, Sawatwong P, Sangmaneedet S, Peruski LF. 2010. Enrichment culture and molecular identification of diverse Bartonella species in stray dogs. Vet Microbiol. 146(3–4):314–319. doi:[10.1016/j.vetmic.2010.05.017](https://doi.org/10.1016/j.vetmic.2010.05.017).

Baker J, Barton MD, Lanser J. 1999. Campylobacter species in cats and dogs in South Australia. Aust Vet J. 77(10):662–666. doi:[10.1111/j.1751-0813.1999.tb13159.x](https://doi.org/10.1111/j.1751-0813.1999.tb13159.x).

Bakirci S, Bilgic H, Köse O, Aksulu ayça, Hacılarlıoğlu S, Erdogan H, Karagenc T. 2016. Molecular and seroprevalence of canine visceral leishmaniasis in West Anatolia, Turkey. Turkish J Vet Anim Sci. 40:637–644. doi:[10.3906/vet-1508-73](https://doi.org/10.3906/vet-1508-73).

Balakrishnan N, Musulin S, Varanat M, Bradley JM, Breitschwerdt EB. 2014. Serological and molecular prevalence of selected canine vector borne pathogens in blood donor candidates, clinically healthy volunteers, and stray dogs in North Carolina. Parasit Vectors. 7:116. doi:[10.1186/1756-3305-7-116](https://doi.org/10.1186/1756-3305-7-116).

Balan LU, Yerbes IM, Piña MAN, Balmes J, Pascual A, Hernández O, Lopez R, Monteón V. 2011. Higher seroprevalence of *Trypanosoma cruzi* infection in dogs than in humans in an urban area of Campeche, Mexico. Vector Borne Zoonotic Dis. 11(7):843–844. doi:[10.1089/vbz.2010.0039](https://doi.org/10.1089/vbz.2010.0039).

Balkaya I, Avcioglu H. 2011. Gastro-Intestinal Helminths Detected by Coprological Examination in Stray Dogs in the Erzurum Province -Turkey. Kafkas Univ Vet Fak Derg. 17:S43–S46.

Bamorovat M, Sharifi I, Dabiri S, Mohammadi MA, Fasihi Harandi M, Mohebali M, Aflatoonian MR, Keyhani A. 2015. *Leishmania tropica* in Stray Dogs in Southeast Iran. Iran J Public Health. 44(10):1359–1366.

Bamorovat M, Sharifi I, Mohammadi MA, Fasihi Harandi M, Mohebali M, Malekpour Afshar R, Babaei Z, Ziaali N, Aflatoonian MR. 2014. Canine Visceral Leishmaniasis in Kerman, Southeast of Iran: A seroepidemiological, histopathological and molecular study. Iran J Parasitol. 9(3):342–349.

Baneth G, Waner T, Koplah A, Weinstein S, Keysary A. 1996. Survey of *Ehrlichia canis* antibodies among dogs in Israel. Vet Rec. 138(11):257–259. doi:[10.1136/vr.138.11.257](https://doi.org/10.1136/vr.138.11.257).

Barati A, Razmi G. 2018. A parasitologic and molecular survey of *Hepatozoon canis* infection in stray dogs in Northeastern Iran. J Parasitol. 104(4):413–417. doi:[10.1645/17-105](https://doi.org/10.1645/17-105).

Barker EN, Langton DA, Helps CR, Brown G, Malik R, Shaw SE, Tasker S. 2012. Haemoparasites of free-roaming dogs associated with several remote Aboriginal communities in Australia. BMC Vet Res. 8(1):55. doi:[10.1186/1746-6148-8-55](https://doi.org/10.1186/1746-6148-8-55).

Barnes A, Bell SC, Isherwood DR, Bennett M, Carter SD. 2000. Evidence of *Bartonella henselae* infection in cats and dogs in the United Kingdom. Vet Rec. 147(24):673–677.

Batamuzi EK, Kassuku AA, Agger JF. 1992. Risk factors associated with canine transmissible venereal tumour in Tanzania. Prev Vet Med. 13(1):13–17. doi:[10.1016/0167-5877(92)90031-A](https://doi.org/10.1016/0167-5877(92)90031-A).

Beard CB, Pye G, Steurer FJ, Rodriguez R, Campman R, Peterson AT, Ramsey J, Wirtz RA, Robinson LE. 2003. Chagas disease in a domestic transmission cycle, southern Texas, USA. Emerg Infect Dis. 9(1):103–105. doi:[10.3201/eid0901.020217](https://doi.org/10.3201/eid0901.020217).

Beiromvand M RE Akhlaghi L, Fattahi Massom SH, Meamar AR, Motevalian A, Oormazdi H. 2013. Prevalence of zoonotic intestinal parasites in domestic and stray dogs in a rural area of Iran. Prev Vet Med. 109(1–2):162–167. doi:[10.1016/j.prevetmed.2012.09.009](https://doi.org/10.1016/j.prevetmed.2012.09.009).

Belkhiria J FTB Chomel BB, Ben Hamida T, Kasten RW, Stuckey MJ, Fleischman DA, Christopher MM, Boulouis HJ. 2017. Prevalence and Potential Risk Factors for Bartonella Infection in Tunisian Stray Dogs. Vector Borne Zoonotic Dis. 17(6):388–397. doi:[10.1089/vbz.2016.2039](https://doi.org/10.1089/vbz.2016.2039).

Bell ET, Toribio J a. LML, White JD, Malik R, Norris JM. 2006. Seroprevalence study of feline coronavirus in owned and feral cats in Sydney, Australia. Aust Vet J. 84(3):74–81. doi:[10.1111/j.1751-0813.2006.tb12231.x](https://doi.org/10.1111/j.1751-0813.2006.tb12231.x).

Belsare AV, Vanak AT, Gompper ME. 2014. Epidemiology of viral pathogens of free-ranging dogs and Indian foxes in a human-dominated landscape in central India. Transbound Emerg Dis. 61 Suppl 1:78–86. doi:[10.1111/tbed.12265](https://doi.org/10.1111/tbed.12265).

Benacer D, Thong KL, Ooi PT, Souris M, Lewis JW, Ahmed AA, Mohd Zain SN. 2017. Serological and molecular identification of *Leptospira* spp. in swine and stray dogs from Malaysia. Trop Biomed. 34(1):89–97.

Bergh K, Bevanger L, Hanssen I, Løseth K. 2002. Low prevalence of *Bartonella henselae* infections in Norwegian domestic and feral cats. APMIS. 110(4):309–314. doi:[10.1034/j.1600-0463.2002.100405.x](https://doi.org/10.1034/j.1600-0463.2002.100405.x).

Bessas A, Leulmi H, Bitam I, Zaidi S, Ait-Oudhia K, Raoult D, Parola P. 2016. Molecular evidence of vector-borne pathogens in dogs and cats and their ectoparasites in Algiers, Algeria. Comp Immunol Microbiol Infect Dis. 45:23–28. doi:[10.1016/j.cimid.2016.01.002](https://doi.org/10.1016/j.cimid.2016.01.002).

Bigdeli M, Namavari M, Moazeni-Ju F, Sadeghzade S, Mirzaei A. 2011. First Study Prevalence of Brucellosis in Stray and Herding Dogs South of Iran. Journal of Animal and Veterinary Advances. 10:1322–1326. doi:[10.3923/javaa.2011.1322.1326](https://doi.org/10.3923/javaa.2011.1322.1326).

Birkenheuer AJ, Levy MG, Stebbins M, Poore M, Breitschwerdt E. 2003. Serosurvey of anti-*Babesia* antibodies in stray dogs and American pit bull terriers and American staffordshire terriers from North Carolina. J Am Anim Hosp Assoc. 39(6):551–557. doi:[10.5326/0390551](https://doi.org/10.5326/0390551).

Blum Domínguez SDC, Chi Dzib MY, Maldonado Velázquez MG, Nuñez Oreza LA, Gómez Solano MI, Caballero Poot RI, Tamay Segovia P. 2013. Detection of reactive canines to *Leptospira* in Campeche City, Mexico. Rev Argent Microbiol. 45(1):34–38.

Bogićević N, Radovanović M, Vasic A, Manić M, Marić J, Vojinovic D, Rogožarski D, Gligić A, Valčić M. 2016. Seroprevalence of *Ehrlichia canis* infection in stray dogs from Serbia. Maced Vet Rev. 40(1):37-42. doi:[10.1515/macvetrev-2016-0096](https://doi.org/10.1515/macvetrev-2016-0096).

Borji H, Sadeghi H, Razmi G, Pozio E, La Rosa G. 2012. Trichinella infection in wildlife of northeast of iran. Iran J Parasitol. 7(4):57–61.

Borthakur SK, Deka DK, Islam S, Sarmah PC. 2015. Occult dirofilariosis in dogs of north eastern region in India. J Arthropod Borne Dis. 10(1):92–97.

Boughattas S, Behnke J, Sharma A, Abu-Madi M. 2017. Seroprevalence of *Toxoplasma gondii* infection in feral cats in Qatar. BMC Veterinary Research. 13(1):26. doi:[10.1186/s12917-017-0952-4](https://doi.org/10.1186/s12917-017-0952-4).

Branley J BR Wolfson C, Waters P, Gottlieb T. 1996. Prevalence of *Bartonella henselae* bacteremia, the causative agent of cat scratch disease, in an Australian cat population. Pathology. 28(3):262–265. doi:[10.1080/00313029600169124](https://doi.org/10.1080/00313029600169124).

Brown GK, Canfield PJ, Dunstan RH, Roberts TK, Martin AR, Brown CS, Irving R. 2006. Detection of *Anaplasma platys* and *Babesia canis vogeli* and their impact on platelet numbers in free-roaming dogs associated with remote Aboriginal communities in Australia. Aust Vet J. 84(9):321–325. doi:[10.1111/j.1751-0813.2006.00029.x](https://doi.org/10.1111/j.1751-0813.2006.00029.x).

Brown J, Blue JL, Wooley RE, Dreesen DW. 1976. *Brucella canis* infectivity rates in stray and pet dog populations. Am J Public Health. 66(9):889–891.

Buishi I, Njoroge E, Zeyhle E, Rogan MT, Craig PS. 2006. Canine echinococcosis in Turkana (north-western Kenya): a coproantigen survey in the previous hydatid-control area and an analysis of risk factors. Ann Trop Med Parasitol. 100(7):601–610. doi:[10.1179/136485906X118503](https://doi.org/10.1179/136485906X118503).

Buishi IE, Njoroge EM, Bouamra O, Craig PS. 2005. Canine echinococcosis in northwest Libya: assessment of coproantigen ELISA, and a survey of infection with analysis of risk-factors. Vet Parasitol. 130(3–4):223–232. doi:[10.1016/j.vetpar.2005.03.004](https://doi.org/10.1016/j.vetpar.2005.03.004).

Cantó GJ A-TG Guerrero RI, Olvera-Ramírez AM, Milián F, Mosqueda J. 2013. Prevalence of fleas and gastrointestinal parasites in free-roaming cats in central Mexico. PLoS One. 8(4):e60744. doi:[10.1371/journal.pone.0060744](https://doi.org/10.1371/journal.pone.0060744).

Cantó GJ, García MP, García A, Guerrero MJ, Mosqueda J. 2011. The prevalence and abundance of helminth parasites in stray dogs from the city of Queretaro in central Mexico. J Helminthol. 85(3):263–269. doi:[10.1017/S0022149X10000544](https://doi.org/10.1017/S0022149X10000544).

Capelli G, Poglayen G, Bertotti F, Giupponi S, Martini M. 1996. The host-parasite relationship in canine heartworm infection in a hyperendemic area of Italy. Vet Res Commun. 20(4):320–330. doi:[10.1007/BF00366538](https://doi.org/10.1007/BF00366538).

Carlos RSA, Albuquerque GR, Bezerra RA, Sicupira PML, Munhoz AD, Lopes CWG. 2010. Ocorrência de anticorpos anti-*Toxoplasma gondii* e principais fatores de risco associados à infecção canina na região de Ilhéus-Itabuna, estado da Bahia. Bras J Vet Med. 32(2):115–121.

Carpenter MA, Brown EW, MacDonald DW, O’brien SJ. 1998. Phylogeographic patterns of feline immunodeficiency virus genetic diversity in the domestic cat. Virology. 251(2):234–243. doi:[10.1006/viro.1998.9402](https://doi.org/10.1006/viro.1998.9402).

Case JB, Chomel B, Nicholson W, Foley JE. 2006. Serological survey of vector-borne zoonotic pathogens in pet cats and cats from animal shelters and feral colonies. J Feline Med Surg. 8(2):111–117. doi:[10.1016/j.jfms.2005.10.004](https://doi.org/10.1016/j.jfms.2005.10.004).

Castanheira P, Duarte A, Gil S, Cartaxeiro C, Malta M, Vieira S, Tavares L. 2014. Molecular and serological surveillance of canine enteric viruses in stray dogs from Vila do Maio, Cape Verde. BMC Vet Res. 10:91. doi:[10.1186/1746-6148-10-91](https://doi.org/10.1186/1746-6148-10-91).

Causapé AC del CE Quílez J, Sánchez-Acedo C. 1996. Prevalence of intestinal parasites, including *Cryptosporidium parvum*, in dogs in Zaragoza city, Spain. Vet Parasitol. 67(3–4):161–167. doi:[10.1016/s0304-4017(96)01033-3](https://doi.org/10.1016/s0304-4017(96)01033-3).

Cave TA, Golder MC, Simpson J, Addie DD. 2004. Risk factors for feline coronavirus seropositivity in cats relinquished to a UK rescue charity. J Feline Med Surg. 6(2):53–58. doi:[10.1016/j.jfms.2004.01.003](https://doi.org/10.1016/j.jfms.2004.01.003).

Cedillo-Peláez C, Díaz-Figueroa ID, Jiménez-Seres MI, Sánchez-Hernández G, Correa D. 2012. Frequency of antibodies to *Toxoplasma gondii* in stray dogs of Oaxaca, México. J Parasitol. 98(4):871–872. doi:[10.1645/GE-3095.1](https://doi.org/10.1645/GE-3095.1).

Celebi B, Carhan A, Kilic S, Babur C. 2010. Detection and genetic diversity of *Bartonella vinsonii subsp. berkhoffii* strains isolated from dogs in Ankara, Turkey. J Vet Med Sci. 72(8):969–973. doi:[10.1292/jvms.09-0466](https://doi.org/10.1292/jvms.09-0466).

Celebi B, Taylan Ozkan A, Kilic S, Akca A, Koenhemsi L, Pasa S, Yildiz K, Mamak N, Guzel M. 2010. Seroprevalence of *Bartonella vinsonii* subsp. berkhoffii in urban and rural dogs in Turkey. J Vet Med Sci. 72(11):1491–1494. doi:[10.1292/jvms.10-0188](https://doi.org/10.1292/jvms.10-0188).

Chattha M, Aslam A, Rehman Z ur, Khan J, Avais M. 2009. Prevalence of *Toxocara Canis* infection in dogs and its effects on various blood parameters in Lahore (Pakistan). J Anim Plant Sci. 19.

Chee J-H, Kwon J-K, Cho H-S, Cho K-O, Lee Y-J, Abd El-Aty AM, Shin S-S. 2008. A survey of ectoparasite infestations in stray dogs of Gwang-ju City, Republic of Korea. Korean J Parasitol. 46(1):23–27. doi:[10.3347/kjp.2008.46.1.23](https://doi.org/10.3347/kjp.2008.46.1.23).

Chikweto A, Bhaiyat MI, Tiwari KP, de Allie C, Sharma RN. 2012. Spirocercosis in owned and stray dogs in Grenada. Vet Parasitol. 190(3–4):613–616. doi:[10.1016/j.vetpar.2012.07.006](https://doi.org/10.1016/j.vetpar.2012.07.006).

Chikweto A, Kumthekar S, Chawla P, Tiwari KP, Perea LM, Paterson T, Sharma RN. 2014. Seroprevalence of *Trypanosoma cruzi* in stray and pet dogs in Grenada, West Indies. Trop Biomed. 31(2):347–350.

Childs JE, Rooney JA, Cooper JL, Olson JG, Regnery RL. 1994. Epidemiologic observations on infection with *Rochalimaea* species among cats living in Baltimore, Md. J Am Vet Med Assoc. 204(11):1775–1778.

Chinyoka S, Dhliwayo S, Marabini L, Dutlow K, Matope G, Pfukenyi DM. 2014. Serological survey of *Brucella canis* in dogs in urban Harare and selected rural communities in Zimbabwe. J S Afr Vet Assoc. 85(1):e1–e5. doi:[10.4102/jsava.v85i1.1087](https://doi.org/10.4102/jsava.v85i1.1087).

Chou C-H, Yeh T-M, Lu Y-P, Shih W-L, Chang C-D, Chien C-H, Liu S-S, Wu H-Y, Tsai F-J, Huang HH, et al. 2014. Prevalence of zoonotic pathogens by molecular detection in stray dogs in Central Taiwan. Thai J Vet Med. 44(3):363–375.

Ciucă L., Genchi M, Kramer L, Mangia C, Miron LD, Prete LD, Maurelli MP, Cringoli G, Rinaldi L. 2016. Heat treatment of serum samples from stray dogs naturally exposed to *Dirofilaria immitis* and *Dirofilaria repens* in Romania. Vet Parasitol. 225:81–85. doi:[10.1016/j.vetpar.2016.05.032](https://doi.org/10.1016/j.vetpar.2016.05.032).

Ciucă Lavinia, Musella V, Miron LD, Maurelli MP, Cringoli G, Bosco A, Rinaldi L. 2016. Geographic distribution of canine heartworm (*Dirofilaria immitis*) infection in stray dogs of eastern Romania. Geospat Health. 11(3):499. doi:[10.4081/gh.2016.499](https://doi.org/10.4081/gh.2016.499).

van der Colf BE, van Zyl GU, Noden BH, Ntirampeba D. 2020. Seroprevalence of *Toxoplasma gondii* infection among pregnant women in Windhoek, Namibia, in 2016. S Afr J Infect Dis. 35(1):25. doi:[10.4102/sajid.v35i1.25](https://doi.org/10.4102/sajid.v35i1.25).

Collantes-Fernández E, Gómez-Bautista M, Miró G, Alvarez-García G, Pereira-Bueno J, Frisuelos C, Ortega-Mora LM. 2008. Seroprevalence and risk factors associated with *Neospora caninum* infection in different dog populations in Spain. Vet Parasitol. 152(1–2):148–151. doi:[10.1016/j.vetpar.2007.12.005](https://doi.org/10.1016/j.vetpar.2007.12.005).

Coman BJ. 1972. A survey of the gastro-intestinal parasites of the feral cat in Victoria. Aust Vet J. 48(4):133–136. doi:[10.1111/j.1751-0813.1972.tb09260.x](https://doi.org/10.1111/j.1751-0813.1972.tb09260.x).

Coman BJ, Jones EH, Driesen MA. 1981. Helminth parasites and arthropods of feral cats. Aust Vet J. 57(7):324–327. doi:[10.1111/j.1751-0813.1981.tb05837.x](https://doi.org/10.1111/j.1751-0813.1981.tb05837.x).

Coman BJ, Jones EH, Westbury HA. 1981. Protozoan and viral infections of feral cats. Aust Vet J. 57:319–323.

Cornell HN, O’Neal PR, Wong VM, Noah DL. 2017. Survey of intestinal parasitism in dogs in the Phoenix metropolitan area. J Am Vet Med Assoc. 251(5):539–543. doi:[10.2460/javma.251.5.539](https://doi.org/10.2460/javma.251.5.539).

Courchamp F, Say L, Pontier D. 2000. Transmission of Feline Immunodeficiency Virus in a population of cats (*Felis catus*). Wildl Res. 27(6):603–611. doi:[10.1071/wr99049](https://doi.org/10.1071/wr99049).

Cruz-Chan JV, Bolio-González M, Colín-Flores R, Ramirez-Sierra MJ, Quijano-Hernandez I, Dumonteil E. 2009. Immunopathology of natural infection with *Trypanosoma cruzi* in dogs. Vet Parasitol. 162(1–2):151–155. doi:[10.1016/j.vetpar.2009.02.024](https://doi.org/10.1016/j.vetpar.2009.02.024).

Cui Y, Yan Y, Wang X, Cao S, Zhang Y, Jian F, Zhang L, Wang R, Shi K, Ning C. 2017. First molecular evidence of mixed infections of *Anaplasma* species in dogs in Henan, China. Ticks Tick Borne Dis. 8(2):283–289. doi:[10.1016/j.ttbdis.2016.12.001](https://doi.org/10.1016/j.ttbdis.2016.12.001).

Curi NH de A, Massara RL, de Oliveira Paschoal AM, Soriano-Araújo A, Lobato ZIP, Demétrio GR, Chiarello AG, Passamani M. 2016. Prevalence and risk factors for viral exposure in rural dogs around protected areas of the Atlantic forest. BMC Vet Res. 12:21. doi:[10.1186/s12917-016-0646-3](https://doi.org/10.1186/s12917-016-0646-3).

Curi NH de A, Miranda I, Talamoni SA. 2006. Serologic evidence of *Leishmania* infection in free-ranging wild and domestic canids around a Brazilian National Park. Mem Inst Oswaldo Cruz. 101:99–101. doi:[10.1590/S0074-02762006000100019](https://doi.org/10.1590/S0074-02762006000100019).

Curi NH de A, Paschoal AM de O, Massara RL, Marcelino AP, Ribeiro AA, Passamani M, Demétrio GR, Chiarello AG. 2014. Factors associated with the seroprevalence of Leishmaniasis in dogs living around Atlantic forest fragments. Plos One. 9(8):e104003. doi:[10.1371/journal.pone.0104003](https://doi.org/10.1371/journal.pone.0104003).

Dada BJ, Adegboye DS, Mohammed AN. 1979. A survey of gastro intestinal helminth parasites of stray dogs in Zaria, Nigeria. Vet Rec. 104(7):145–146. doi:[10.1136/vr.104.7.145](https://doi.org/10.1136/vr.104.7.145).

Dağ S, Sözmen M, Cihan M, Tunca R, Kurt B, Devrim AK, Özen H. 2016. Gastric Helicobacter-like organisms in stray cats: identification, prevalence, and pathologic association. Pak Vet J. 36:199-203.

Dakkak A, El Berbri I, Petavy AF, Boué F, Bouslikhane M, Fassi Fihri O, Welburn S, Ducrotoy MJ. 2017. *Echinococcus granulosus* infection in dogs in Sidi Kacem Province (North-West Morocco). Acta Trop. 165:26–32. doi:[10.1016/j.actatropica.2016.07.007](https://doi.org/10.1016/j.actatropica.2016.07.007).

Dalimi A, Motamedi G, Hosseini M, Mohammadian B, Malaki H, Ghamari Z, Ghaffari Far F. 2002. Echinococcosis/Hydatidosis in western Iran. Vet Parasitol. 105(2):161–171. doi:[10.1016/s0304-4017(02)00005-5](https://doi.org/10.1016/s0304-4017(02)00005-5).

Dalimi A, Sattari A, Motamedi Gh. 2006. A study on intestinal helminthes of dogs, foxes and jackals in the western part of Iran. Vet Parasitol. 142(1):129–133. doi:[10.1016/j.vetpar.2006.06.024](https://doi.org/10.1016/j.vetpar.2006.06.024).

Dalimi A, Shamsi M, Khosravi A, Ghaffarifar F. 2017. Genotyping *Echinococcus granulosus* from canine isolates in Ilam Province, West of Iran. Iran J Parasitol. 12(4):614–621.

Danesi P, Furnari C, Granato A, Schivo A, Otranto D, Capelli G, Cafarchia C. 2014. Molecular identity and prevalence of *Cryptococcus* spp. nasal carriage in asymptomatic feral cats in Italy. Med Mycol. 52(7):667–673. doi:[10.1093/mmy/myu030](https://doi.org/10.1093/mmy/myu030).

Danner RM, Goltz DM, Hess SC, Banko PC. 2007. Evidence of Feline Immunodeficiency Virus, Feline Leukemia Virus, and *Toxoplasma gondii* in feral cats on Mauna Kea, Hawaii. J Wildl Dis. 43(2):315–318. doi:[10.7589/0090-3558-43.2.315](https://doi.org/10.7589/0090-3558-43.2.315).

Delahay RJ, Daniels MJ, Macdonald DW, McGuire K, Balharry D. 1998. Do patterns of helminth parasitism differ between groups of wild-living cats in Scotland? J Zool. 245(2):175–183. doi:[10.1111/j.1469-7998.1998.tb00085.x](https://doi.org/10.1111/j.1469-7998.1998.tb00085.x).

Deplazes P, Guscetti F, Wunderlin E, Bucklar H, Skaggs J, Wolff K. 1995. [Endoparasite infection in stray and abandoned dogs in southern Switzerland]. Schweiz Arch Tierheilkd. 137(5):172–179.

Di Francesco A, Donati M, Battelli G, Cevenini R, Baldelli R. 2004. Seroepidemiological survey for *Chlamydophila felis* among household and feral cats in northern Italy. Vet Rec. 155(13):399–400. doi:[10.1136/vr.155.13.399](https://doi.org/10.1136/vr.155.13.399).

Diakou A, Papadopoulos E, Lazarides K. 2009. Specific anti-*Leishmania* spp. antibodies in stray cats in Greece. J Feline Med Surg. 11(8):728–730. doi:[10.1016/j.jfms.2008.01.009](https://doi.org/10.1016/j.jfms.2008.01.009).

Dos-Santos W, Jesus EE, Paranhos-Silva M, Pereira A, Santos JC, Baleeiro CO, Nascimento EG, Moreira E, Oliveira GG, Pontes-de-carvalho L. 2008. Associations among immunological, parasitological and clinical parameters in canine visceral leishmaniasis: Emaciation, spleen parasitism, specific antibodies and leishmanin skin test reaction. Vet Immunol Immunopathol. 123:251-259. doi:[10.1016/j.vetimm.2008.02.004](https://doi.org/10.1016/j.vetimm.2008.02.004).

Duarte A, Castro I, Pereira da Fonseca IM, Almeida V, Madeira de Carvalho LM, Meireles J, Fazendeiro MI, Tavares L, Vaz Y. 2010. Survey of infectious and parasitic diseases in stray cats at the Lisbon Metropolitan Area, Portugal. J Feline Med Surg. 12(6):441–446. doi:[10.1016/j.jfms.2009.11.003](https://doi.org/10.1016/j.jfms.2009.11.003).

Duarte A, Fernandes M, Santos N, Tavares L. 2012. Virological Survey in free-ranging wildcats (*Felis silvestris*) and feral domestic cats in Portugal. Vet Microbiol. 158(3–4):400–404. doi:[10.1016/j.vetmic.2012.02.033](https://doi.org/10.1016/j.vetmic.2012.02.033).

Dubey JP, Darrington C, Tiao N, Ferreira LR, Choudhary S, Molla B, Saville WJA, Tilahun G, Kwok OCH, Gebreyes WA. 2013. Isolation of viable *Toxoplasma gondii* from tissues and feces of cats from Addis Ababa, Ethiopia. J Parasitol. 99(1):56–58. doi:[10.1645/GE-3229.1](https://doi.org/10.1645/GE-3229.1).

Dubey JP, Gennari SM, Sundar N, Vianna MCB, Bandini LM, Yai LEO, Kwok CH, Suf C. 2007. Diverse and atypical genotypes identified in Toxoplasma gondii from dogs in São Paulo, Brazil. J Parasitol. 93(1):60–64. doi:[10.1645/GE-972R.1](https://doi.org/10.1645/GE-972R.1).

Dubey JP, Lappin MR, Kwok OCH, Mofya S, Chikweto A, Baffa A, Doherty D, Shakeri J, Macpherson CNL, Sharma RN. 2009. Seroprevalence of *Toxoplasma gondii* and concurrent *Bartonella* spp., feline immunodeficiency virus, and feline leukemia virus infections in cats from Grenada, West Indies. J Parasitol. 95(5):1129–1133. doi:[10.1645/GE-2114.1](https://doi.org/10.1645/GE-2114.1).

Dubey JP, López-Torres HY, Sundar N, Velmurugan GV, Ajzenberg D, Kwok OCH, Hill R, Dardé ML, Su C. 2007. Mouse-virulent *Toxoplasma gondii* isolated from feral cats on Mona Island, Puerto Rico. J Parasitol. 93(6):1365–1369. doi:[10.1645/GE-1409.1](https://doi.org/10.1645/GE-1409.1).

Dubey JP, Moura L, Majumdar D, Sundar N, Velmurugan GV, Kwok OCH, Kelly P, Krecek RC, Su C. 2009. Isolation and characterization of viable *Toxoplasma gondii* isolates revealed possible high frequency of mixed infection in feral cats ( Felis domesticus) from St Kitts, West Indies. Parasitology. 136(6):589–594. doi:[10.1017/S0031182009006015](https://doi.org/10.1017/S0031182009006015).

Dubey JP SC Rajapakse RP, Wijesundera RR, Sundar N, Velmurugan GV, Kwok OC. 2007. Prevalence of *Toxoplasma gondii* in dogs from Sri Lanka and genetic characterization of the parasite isolates. Vet Parasitol. 146(3–4):341–346. doi:[10.1016/j.vetpar.2007.03.009](https://doi.org/10.1016/j.vetpar.2007.03.009).

Dubey JP, Tilahun G, Boyle JP, Schares G, Verma SK, Ferreira LR, Oliveira S, Tiao N, Darrington C, Gebreyes WA. 2013. Molecular and biological characterization of first isolates of *Hammondia hammondi* from cats from Ethiopia. J Parasitol. 99(4):614–618. doi:[10.1645/12-51.1](https://doi.org/10.1645/12-51.1).

Dubey JP, Tiwari K, Chikweto A, Deallie C, Sharma R, Thomas D, Choudhary S, Ferreira LR, Oliveira S, Verma SK, et al. 2013. Isolation and RFLP genotyping of *Toxoplasma gondii* from the domestic dogs (*Canis familiaris*) from Grenada, West Indies revealed high genetic variability. Vet Parasitol. 197(3–4):623–626. doi:[10.1016/j.vetpar.2013.07.029](https://doi.org/10.1016/j.vetpar.2013.07.029).

Dubinský P, Havasiová-Reiterová K, Petko B, Hovorka I, Tomasovicová O. 1995. Role of small mammals in the epidemiology of Toxocariasis. Parasitology. 110 ( Pt 2):187–193. doi:[10.1017/s0031182000063952](https://doi.org/10.1017/s0031182000063952).

Dybing NA, Jacobson C, Irwin P, Algar D, Adams PJ. 2016. *Bartonella* species identified in rodent and feline hosts from island and mainland Western Australia. Vector-Borne and Zoonotic Dis. 16(4):238–244. doi:[10.1089/vbz.2015.1902](https://doi.org/10.1089/vbz.2015.1902).

Dybing NA, Jacobson C, Irwin P, Algar D, Adams PJ. 2017. *Leptospira* species in feral cats and black rats from Western Australia and Christmas Island. Vector Borne Zoonotic Dis. 17(5):319–324. doi:[10.1089/vbz.2016.1992](https://doi.org/10.1089/vbz.2016.1992).

Dybing NA, Jacobson C, Irwin P, Algar D, Adams PJ. 2018. Challenging the dogma of the ‘Island Syndrome’: a study of helminth parasites of feral cats and black rats on Christmas Island. Australas J Environ Manag. 25(1):99–118. doi:[10.1080/14486563.2017.1417165](https://doi.org/10.1080/14486563.2017.1417165).

Ebrahimzade E, Fattahi R, Ahoo MB. 2016. Ectoparasites of stray dogs in Mazandaran, Gilan and Qazvin Provinces, north and center of Iran. J Arthropod Borne Dis. 10(3):364–369.

Eguía-Aguilar P, Cruz-Reyes A, Martínez-Maya JJ. 2005. Ecological analysis and description of the intestinal helminths present in dogs in Mexico City. Vet Parasitol. 127(2):139–146. doi:[10.1016/j.vetpar.2004.10.004](https://doi.org/10.1016/j.vetpar.2004.10.004).

El Behairy AM, Choudhary S, Ferreira LR, Kwok OCH, Hilali M, Su C, Dubey JP. 2013. Genetic characterization of viable *Toxoplasma gondii* isolates from stray dogs from Giza, Egypt. Vet Parasitol. 193(1–3):25–29. doi:[10.1016/j.vetpar.2012.12.007](https://doi.org/10.1016/j.vetpar.2012.12.007).

El-Gayar AK. 2007. Studies on some trematode parasites of stray dogs in Egypt with a key to the identification of intestinal trematodes of dogs. Vet Parasitol. 144(3–4):360–365. doi:[10.1016/j.vetpar.2006.09.043](https://doi.org/10.1016/j.vetpar.2006.09.043).

El-Shehabi FS, Abdel-Hafez SK, Kamhawi SA. 1999. Prevalence of intestinal helminths of dogs and foxes from Jordan. Parasitol Res. 85(11):928–934. doi:[10.1007/s004360050660](https://doi.org/10.1007/s004360050660).

Erol N, Pasa S. 2013. An Investigation of the Feline Immunodeficiency Virus (FIV) and Feline Leukemia Virus (FeLV) Infections in cats in western Turkey. Acta Scientiae Veterinari. 41:1166.

Eslami A, Ranjbar-Bahadori S, Meshgi B, Dehghan M, Bokaie S. 2010. Helminth Infections of Stray Dogs from Garmsar, Semnan Province, Central Iran. Iran J Parasitol. 5(4):37–41.

Fancourt BA, Jackson RB. 2014. Regional seroprevalence of *Toxoplasma gondii* antibodies in feral and stray cats (*Felis catus*) from Tasmania. Aust J Zool. 62(4):272–283. doi:[10.1071/ZO14015](https://doi.org/10.1071/ZO14015).

Fang F, Li J, Huang T, Guillot J, Huang W. 2015. Zoonotic helminths parasites in the digestive tract of feral dogs and cats in Guangxi, China. BMC Vet Res. 11(1):211. doi:[10.1186/s12917-015-0521-7](https://doi.org/10.1186/s12917-015-0521-7).

Feng T-H, Chou C-C, Yeh T-M, Su Y-C, Lu Y-P, Shih W-L, Chiang C-H, Chang C-D, Liu S-S, Wu H-Y, et al. 2015. Molecular prevalence of zoonotic pathogens in pet and stray dogs in southern Taiwan. Thai J Vet Med. 45(4):509–522.

Fernandez C SRN Chikweto A, Mofya S, Lanum L, Flynn P, Burnett JP, Doherty D. 2010. A serological study of *Dirofilaria immitis* in feral cats in Grenada, West Indies. J Helminthol. 84(4):390–393. doi:[10.1017/s0022149x10000027](https://doi.org/10.1017/s0022149x10000027).

Fernández H, Martin R. 1991. *Campylobacter* intestinal carriage among stray and pet dogs. Rev Saude Publica. 25(6):473–475. doi:[10.1590/s0034-89101991000600009](https://doi.org/10.1590/s0034-89101991000600009).

Fiorello CV, Straub MH, Schwartz LM, Liu J, Campbell A, Kownacki AK, Foley JE. 2017. Multiple-host pathogens in domestic hunting dogs in Nicaragua’s Bosawás Biosphere Reserve. Acta Trop. 167:183–190. doi:[10.1016/j.actatropica.2016.12.020](https://doi.org/10.1016/j.actatropica.2016.12.020).

Fontana FF, dos Santos CTB, Esteves FM, Rocha A, Fernandes GF, do Amaral CC, Domingues MA, De Camargo ZP, Silva-Vergara ML. 2010. Seroepidemiological survey of paracoccidioidomycosis infection among urban and rural dogs from Uberaba, Minas Gerais, Brazil. Mycopathologia. 169(3):159–165. doi:[10.1007/s11046-009-9241-5](https://doi.org/10.1007/s11046-009-9241-5).

Fonzar UJV, Langoni H. 2012. Geographic analysis on the occurrence of human and canine leptospirosis in the city of Maringá, state of Paraná, Brazil. Rev Soc Bras Med Trop. 45:100–105. doi:[10.1590/S0037-86822012000100019](https://doi.org/10.1590/S0037-86822012000100019).

Fraga DBM, Solcà MS, Silva VMG, Borja LS, Nascimento EG, Oliveira GGS, Pontes-de-Carvalho LC, Veras PST, dos-Santos WLC. 2012. Temporal distribution of positive results of tests for detecting *Leishmania* infection in stray dogs of an endemic area of visceral leishmaniasis in the Brazilian tropics: a 13 years survey and association with human disease. Vet Parasitol. 190(3–4):591–594.

Fridlund-Plugge N, Montiani-Ferreira F, Richartz RRTB, Dal Pizzol J, Machado PC, Patrício LFL, Rosinelli AS, Locatelli-Dittrich R. 2008. Frequency of antibodies against *Neospora caninum* in stray and domiciled dogs from urban, periurban and rural areas from Paraná State, Southern Brazil. Rev Bras Parasitol Vet. 17(4):222–226. doi:[10.1590/s1984-29612008000400010](https://doi.org/10.1590/s1984-29612008000400010).

Fromont E, Courchamp F, Pontier D, Artois M. 1997. Infection strategies of retroviruses and social grouping of domestic cats. Can J Zool. 75(12):1994–2002. doi:[10.1139/z97-832](https://doi.org/10.1139/z97-832).

Fromont E, Morvilliers L, Artois M, Pontier D. 2001. Parasite richness and abundance in insular and mainland feral cats: insularity or density? Parasitology. 123(Pt 2):143–151. doi:[10.1017/s0031182001008277](https://doi.org/10.1017/s0031182001008277).

Gauss CBL, Almería S, Ortuño A, Garcia F, Dubey JP. 2003. Seroprevalence of *Toxoplasma gondii* antibodies in domestic cats from Barcelona, Spain. J Parasitol. 89(5):1067–1068. doi:[10.1645/GE-114](https://doi.org/10.1645/GE-114).

Gay N, Soupé-Gilbert M-E, Goarant C. 2014. Though not reservoirs, dogs might Transmit Leptospira in New Caledonia. Int J Environ Res Public Health. 11(4):4316–4325. doi:[10.3390/ijerph110404316](https://doi.org/10.3390/ijerph110404316).

Gencay Göksu A, Oncel T, Karaoglu T, Sancak AA, Demir Altas A, Ozkul A. 2004. Antibody prevalence to Canine Distemper Virus (CDV) in stray dogs in Turkey. Rev Vet Med. 155:432–434.

Gennari SM, Yai LEO, D’Auria SNR, Cardoso SMS, Kwok OCH, Jenkins MC, Dubey JP. 2002. Occurrence of *Neospora caninum* antibodies in sera from dogs of the city of São Paulo, Brazil. Vet Parasitol. 106(2):177–179. doi:[10.1016/s0304-4017(02)00052-3](https://doi.org/10.1016/s0304-4017(02)00052-3).

Ghalmi F, China B, Kaidi R, Losson B. 2009. First epidemiological study on exposure to *Neospora caninum* in different canine populations in the Algiers District (Algeria). Parasitol Int. 58(4):444–450. doi:[10.1016/j.parint.2009.08.008](https://doi.org/10.1016/j.parint.2009.08.008).

Ghil H-M, Yoo J-H, Jung W-S, Chung T-H, Youn H-Y, Hwang C-Y. 2009. Survey of *Helicobacter* infection in domestic and feral cats in Korea. J Vet Sci. 10(1):67–72. doi:[10.4142/jvs.2009.10.1.67](https://doi.org/10.4142/jvs.2009.10.1.67).

Giacomelli M, Follador N, Coppola LM, Martini M, Piccirillo A. 2015. Survey of *Campylobacter* spp. in owned and unowned dogs and cats in Northern Italy. Vet J. 204(3):333–337. doi:[10.1016/j.tvjl.2015.03.017](https://doi.org/10.1016/j.tvjl.2015.03.017).

Giorgobiani E, Chitadze N, Chanturya G, Grdzelidze M, Jochim RC, Machablishvili A, Tushishvili T, Zedginidze Y, Manjgaladze MK, Iashvili N, et al. 2011. Epidemiologic aspects of an emerging focus of Visceral Leishmaniasis in Tbilisi, Georgia. PLOS Negl Trop Dis. 5(12):e1415. doi:[10.1371/journal.pntd.0001415](https://doi.org/10.1371/journal.pntd.0001415).

Gomes L de A, Moraes PHG, do Nascimento L de CS, O’Dwyer LH, Nunes MRT, Rossi ADRP, Aguiar DCF, Gonçalves EC. 2016. Molecular analysis reveals the diversity of *Hepatozoon* species naturally infecting domestic dogs in a northern region of Brazil. Ticks Tick Borne Dis. 7(6):1061–1066. doi:[10.1016/j.ttbdis.2016.09.008](https://doi.org/10.1016/j.ttbdis.2016.09.008).

Gonçalves D, Moura R, Dreer M, Nascimento D, Rodrigues G, Caetano I, Vidotto O, FReitas J, Vieira M. 2015. First record of *Borrelia burgdorferi* sensu lato antibodies in stray dogs in the northwest region of Parana State, Brazil. Semin Cienc Agrar. 36:2641–2648. doi:[10.5433/1679-0359.2015v36n4p2641](https://doi.org/10.5433/1679-0359.2015v36n4p2641).

Goncharuk MS RVV Kerley LL, Naidenko SV. 2012. Prevalence of seropositivity to pathogens in small carnivores in adjacent areas of Lazovskii Reserve. Biol Bull Russ Acad Sci. 39(8):708–713. doi:[10.1134/s1062359012080067](https://doi.org/10.1134/s1062359012080067).

Goni MD. 2017. Occurrence of *Campylobacter* in dogs and cats in Selangor Malaysia and the associated risk factors. Malays J Microbiol. 13(3):164–171.

Gordy JT, Jones CA, Rue J, Crawford PC, Levy JK, Stallknecht DE, Tripp RA, Tompkins SM. 2012. Surveillance of feral cats for influenza A virus in North Central Florida. Influenza Other Respir Viruses. 6(5):341–347. doi:[10.1111/j.1750-2659.2011.00325.x](https://doi.org/10.1111/j.1750-2659.2011.00325.x).

Gregory GG, Munday BL. 1976. Internal parasites of feral cats from the Tasmanian Midlands and King Island. Aust Vet J. 52(7):317–320. doi:[10.1111/j.1751-0813.1976.tb02396.x](https://doi.org/10.1111/j.1751-0813.1976.tb02396.x).

Gruntmeir JM, Adolph C, Thomas JE, Reichard M, Blagburn B, Little S. 2017. Increased detection of *Dirofilaria immitis* antigen in cats after heat pretreatment of samples. J Feline Med Surg. 19(10):1013-1016. doi:[10.1177/1098612X16670562](https://doi.org/10.1177/1098612X16670562).

Guernier V, Lagadec E, Cordonin C, Minter GL, Gomard Y, Pagès F, Jaffar-Bandjee M-C, Michault A, Tortosa P, Dellagi K. 2016. Human Leptospirosis on Reunion Island, Indian Ocean: are rodents the (only) ones to blame? PLOS Negl Trop Dis. 10(6):e0004733. doi:[10.1371/journal.pntd.0004733](https://doi.org/10.1371/journal.pntd.0004733).

Guven E, Avcioglu H, Cengiz S, Hayirli A. 2017. Vector-Borne pathogens in stray dogs in northeastern Turkey. Vector Borne Zoonotic Dis. 17(8):610–617. doi:[10.1089/vbz.2017.2128](https://doi.org/10.1089/vbz.2017.2128).

Haddadzade H, Fattahi R, Mohebali M, Akhoundi B, Ebrahimzade E. 2013. Seroepidemiologcal investigation of Visceral leishmaniasis in stray and owned dogs in Alborz Province, central Iran using direct agglutination test. Iran J Parasitol. 8(1):152–157.

Hafemann DCM, Merlini LS, Gonçalves DD, Fortes MS, Navarro IT, Chiderolli RT, Freitas JC, Gonçalves APP, Rosa G, Sposito PH. 2018. Detection of anti-*Leptospira* spp., anti-*Brucella* spp., and anti-*Toxoplasma* gondii antibodies in stray dogs. Sem Ciec Agrar. 39(1):167–176.

Hamel D, Silaghi C, Lescai D, Pfister K. 2012. Epidemiological aspects on vector-borne infections in stray and pet dogs from Romania and Hungary with focus on *Babesia* spp. Parasitol Res. 110(4):1537–1545. doi:[10.1007/s00436-011-2659-y](https://doi.org/10.1007/s00436-011-2659-y).

Hamidinejat H, Mosalanejad B, Avizeh R, Razi Jalali MH, Ghorbanpoor M, Namavari M. 2011. *Neospora caninum* and *Toxoplasma gondii* antibody prevalence in Ahvaz feral cats, Iran. Jundishapur J Microbiol. 4(4):217-222.

Hammond-Aryee K, Esser M, van Helden L, van Helden P. 2015. A high seroprevalence of *Toxoplasma gondii* antibodies in a population of feral cats in the Western Cape province of South Africa. S Afr J Infect Dis. 30(4):141–144. doi:[10.1080/23120053.2015.1107295](https://doi.org/10.1080/23120053.2015.1107295).

Havas K, Burkman K. 2011. A comparison of the serological evidence of *Coxiella burnetii* exposure between military working dogs and feral canines in Iraq. Mil Med. 176(10):1101–1103. doi:[10.7205/milmed-d-11-00025](https://doi.org/10.7205/milmed-d-11-00025).

Hayward JJ, Taylor J, Rodrigo AG. 2007. Phylogenetic analysis of Feline Immunodeficiency Virus in feral and companion domestic cats of New Zealand. J Virol. 81(6):2999–3004. doi:[10.1128/JVI.02090-06](https://doi.org/10.1128/JVI.02090-06).

Headley SA, Gillen MA, Sanches AWD, Satti MZ. 2012. *Platynosomum fastosum*-induced chronic intrahepatic cholangitis and *Spirometra* spp. infections in feral cats from Grand Cayman. J Helminthol. 86(2):209–214. doi:[10.1017/S0022149X11000265](https://doi.org/10.1017/S0022149X11000265).

Hellard E, Fouchet D, Santin-Janin H, Tarin B, Badol V, Coupier C, Leblanc G, Poulet H, Pontier D. 2011. When cats’ ways of life interact with their viruses: a study in 15 natural populations of owned and unowned cats (*Felis silvestris catus*). Prev Vet Med. 101(3–4):250–264. doi:[10.1016/j.prevetmed.2011.04.020](https://doi.org/10.1016/j.prevetmed.2011.04.020).

Henn JB, VanHorn BA, Kasten RW, Kachani M, Chomel BB. 2006. Short report: Antibodies to *Bartonella vinsonii* *subsp. berkhoffii* in Moroccan dogs. Am J Trop Med. 74(2):222–223.

Herrera HM, Rocha FL, Lisboa CV, Rademaker V, Mourão GM, Jansen AM. 2011. Food web connections and the transmission cycles of *Trypanosoma cruzi* and *Trypanosoma evansi* (Kinetoplastida, Trypanosomatidae) in the Pantanal Region, Brazil. Trans R Soc Trop Med Hyg. 105(7):380–387. doi:[10.1016/j.trstmh.2011.04.008](https://doi.org/10.1016/j.trstmh.2011.04.008).

Hoff B, McEwen B, Peregrine AS. 2008. A survey for infection with *Dirofilaria immitis*, *Ehrlichia canis*, *Borrelia burgdorferi*, and *Babesia canis* in feral and client-owned dogs in the Turks and Caicos Islands, British West Indies. Can Vet J. 49(6):593–594.

Hoida G, Greenberg Z, Furth M, Malsha Y, Craig PS, Schantz PM, Sneir R, el-On J. 1998. An epidemiological survey of *Echinococcus granulosus* and other helminths in animal populations in northern Israel. J Helminthol. 72(2):127–131. doi:[10.1017/s0022149x00016308](https://doi.org/10.1017/s0022149x00016308).

Hornok S, Edelhofer R, Fok E, Berta K, Fejes P, Répási A, Farkas R. 2006. Canine Neosporosis in Hungary: screening for seroconversion of household, herding and stray dogs. Vet Parasitol. 137(3–4):197–201. doi:[10.1016/j.vetpar.2006.01.030](https://doi.org/10.1016/j.vetpar.2006.01.030).

Hornok S, Edelhofer R, Joachim A, Farkas R, Berta K, Répási A, Lakatos B. 2008. Seroprevalence of *Toxoplasma gondii* and *Neospora caninum* infection of cats in Hungary. Acta Vet Hung. 56(1):81–88. doi:[10.1556/AVet.56.2008.1.8](https://doi.org/10.1556/AVet.56.2008.1.8).

Hosseininejad M, Malmasi A, Hosseini F, Selk-Ghaffari M, Khorrami N, Mohebali M, Shojaee S, Mirani A, Azizzadeh M, Mirshokraei P, et al. 2011. Seroprevalence of *Toxoplasma gondii* infection in dogs in Tehran, Iran. Iran J Parasitol. 6(1):81–85.

Hosseininejad M, Mohebali M, Hosseini F, Karimi S, Sharifzad S, Akhoundi B. 2012. Seroprevalence of canine Visceral Leishmaniasis in asymptomatic dogs in Iran. Iran J Vet Res. 13:54–57.

Hou H, Cao L, Ren W, Wang D, Ding H, You J, Yao X, Dong H, Guo Y, Yuan S, et al. 2017. Seroprevalence of *Dirofilaria immitis* in cats from Liaoning Province, Northeastern China. Korean J Parasitol. 55(6):673–677. doi:[10.3347/kjp.2017.55.6.673](https://doi.org/10.3347/kjp.2017.55.6.673).

Hwang J, Gottdenker N, Min M-S, Lee H, Chun M-S. 2016. Evaluation of biochemical and haematological parameters and prevalence of selected pathogens in feral cats from urban and rural habitats in South Korea. J Feline Med Surg. 18(6):443–451. doi:[10.1177/1098612X15587572](https://doi.org/10.1177/1098612X15587572).

Hwang J, Gottdenker N, Oh D-H, Lee H, Chun M-S. 2017. Infections by pathogens with different transmission modes in feral cats from urban and rural areas of Korea. J Vet Sci. 18(4):541–545. doi:[10.4142/jvs.2017.18.4.541](https://doi.org/10.4142/jvs.2017.18.4.541).

Hwang J, Kang J-G, Oh S-S, Chae J-B, Cho Y-K, Cho Y-S, Lee H, Chae J-S. 2017. Molecular detection of Severe Fever With Thrombocytopenia Syndrome Virus (SFTSV) in feral cats from Seoul, Korea. Ticks Tick Borne Dis. 8(1):9–12. doi:[10.1016/j.ttbdis.2016.08.005](https://doi.org/10.1016/j.ttbdis.2016.08.005).

Inangolet FO, Biffa D, Opuda-Asibo J, Oloya J, Skjerve E. 2010. Distribution and intensity of *Echinococcus granulosus* infections in dogs in Moroto District, Uganda. Trop Anim Health Prod. 42(7):1451–1457. doi:[10.1007/s11250-010-9574-6](https://doi.org/10.1007/s11250-010-9574-6).

Inokuma H, Nane G, Uechi T, Yonahara Y, Brouqui P, Okuda M, Onishi T. 2001. Survey of tick infestation and tick-borne ehrlichial infection of dogs in Ishigaki Island, Japan. J Vet Med Sci. 63(11):1225–1227. doi:[10.1292/jvms.63.1225](https://doi.org/10.1292/jvms.63.1225).

Inokuma H, Yoshizaki Y, Matsumoto K, Okuda M, Onishi T, Nakagome K, Kosugi R, Hirakawa M. 2004. Molecular survey of *Babesia* infection in dogs in Okinawa, Japan. Vet Parasitol. 121(3–4):341–346. doi:[10.1016/j.vetpar.2004.03.012](https://doi.org/10.1016/j.vetpar.2004.03.012).

Inoue K JS Maruyama S, Kabeya H, Kawanami K, Yanai K, Jitchum S. 2009. Prevalence of *Bartonella* infection in cats and dogs in a metropolitan area, Thailand. Epidemiol Infect. 137(11):1568–1573. doi:[10.1017/s095026880900257x](https://doi.org/10.1017/s095026880900257x).

Iorio R, Cafarchia C, Capelli G, Fasciocco D, Otranto D, Giangaspero A. 2007. Dermatophytoses in cats and humans in central Italy: epidemiological aspects. Mycoses. 50(6):491–495. doi:[10.1111/j.1439-0507.2007.01385.x](https://doi.org/10.1111/j.1439-0507.2007.01385.x).

Iraqi W. 2017. Canine echinococcosis: the predominance of immature eggs in adult tapeworms of *Echinococcus granulosus* in stray dogs from Tunisia. J Helminthol. 91(3):380–383. doi:[10.1017/s0022149x16000341](https://doi.org/10.1017/s0022149x16000341).

Islam A, Rahman M, Islam S, Debnath P, Alam M, Hassan M. 2017. Sero-prevalence of visceral leishmaniasis (VL) among dogs in VL endemic areas of Mymensingh distict, Bangladesh. Journal of Advanced Veterinary and Animal Research. 4(3):241-248. doi:[10.5455/javar.2017.d217](https://doi.org/10.5455/javar.2017.d217).

Ito J, Papasarathorn T, Tongkoom B. 1962. An investigation of parasitic helminths of stray dogs in Bangkok. Jpn J Med Sci Biol. 15:53–60. doi:[10.7883/yoken1952.15.53](https://doi.org/10.7883/yoken1952.15.53).

Jadoon A, Akhtar T, Maqbool A, Anjum A, Ajmal A. 2009. Seroprevalence of *Toxoplasma gondii* in canines. J Anim Plant Sci. 19(4):179-181.

Jimenez-Coello M, Ortega-Pacheco A, Guzman-Marin E, Guiris-Andrade DM, Martinez-Figueroa L, Acosta-Viana KY. 2010. Stray dogs as reservoirs of the zoonotic agents *Leptospira interrogans*, *Trypanosoma cruzi*, and *Aspergillus* spp. in an urban area of Chiapas in southern Mexico. Vector Borne Zoonotic Dis. 10(2):135–141. doi:[10.1089/vbz.2008.0170](https://doi.org/10.1089/vbz.2008.0170).

Jiménez-Coello M, Pérez-Osorio C, Vado-Solís I, Rodríguez-Buenfil JC, Ortega-Pacheco A. 2009. Serological survey of *Ehrlichia canis* in stray dogs from Yucatan, Mexico, using two different diagnostic tests. Vector Borne Zoonotic Dis. 9(2):209–212. doi:[10.1089/vbz.2008.0039](https://doi.org/10.1089/vbz.2008.0039).

Jimenez-Coello M, Poot-Cob M, Ortega-Pacheco A, Guzman-Marin E, Ramos-Ligonio A, Sauri-Arceo CH, Acosta-Viana KY. 2008. American Trypanosomiasis in dogs from an urban and rural area of Yucatan, Mexico. Vector Borne Zoonotic Dis. 8(6):755–761. doi:[10.1089/vbz.2007.0224](https://doi.org/10.1089/vbz.2007.0224).

Jittapalapong S MS Nimsupan B, Pinyopanuwat N, Chimnoi W, Kabeya H. 2007. Seroprevalence of *Toxoplasma gondii* antibodies in stray cats and dogs in the Bangkok metropolitan area, Thailand. Vet Parasitol. 145(1–2):138–141. doi:[10.1016/j.vetpar.2006.10.021](https://doi.org/10.1016/j.vetpar.2006.10.021).

Jittapalapong S, Rungphisutthipongse O, Maruyama S, Schaefer JJ, Stich RW. 2006. Detection of *Hepatozoon canis* in stray dogs and cats in Bangkok, Thailand. Ann N Y Acad Sci. 1081:479–488. doi:[10.1196/annals.1373.071](https://doi.org/10.1196/annals.1373.071).

Jittapalapong S, Sittisan P, Sakpuaram T, Kabeya H, Maruyama S, Inpankaew T. 2009. Coinfection of *Leptospira* spp and *Toxoplasma gondii* among stray dogs in Bangkok, Thailand. Southeast Asian J Trop Med Public Health. 40(2):247–252.

Johnson EM RMV Nagamori Y, Duncan-Decocq RA, Whitley PN, Ramachandran A. 2017. Prevalence of *Alaria* infection in companion animals in north central Oklahoma from 2006 through 2015 and detection in wildlife. J Am Vet Med Assoc. 250(8):881–886. doi:[10.2460/javma.250.8.881](https://doi.org/10.2460/javma.250.8.881).

Johnston J, Gasser R. 1993. Copro-Parasitological survey of dogs in southern Victoria. Aust Vet Pract. 23(3):127–131.

Jorge RSP, Ferreira F, Ferreira Neto JS, Vasconcellos S de A, Lima E de S, Morais ZM de, Souza GO de. 2011. Exposure of free-ranging wild carnivores, horses and domestic dogs to *Leptospira* spp in the northern Pantanal, Brazil. Mem Inst Oswaldo Cruz. 106:441–444. doi:[10.1590/S0074-02762011000400009](https://doi.org/10.1590/S0074-02762011000400009).

Kamani J, Harrus S, Nachum-Biala Y, Salant H, Mumcuoglu KY, Baneth G. 2018. Pathogenic and endosymbiont apicomplexans in *Ctenocephalides felis* (Siphonaptera: Pulicidae) from cats in Jerusalem, Israel. Comp Immunol Microbiol Infect Dis. 57:29–33. doi:[10.1016/j.cimid.2018.03.002](https://doi.org/10.1016/j.cimid.2018.03.002).

Kang Y-H, Cong W, Qin S-Y, Shan X-F, Gao Y-H, Wang C-F, Qian A-D. 2016. First report of *Toxoplasma gondii*, *Dirofilaria immitis*, and *Chlamydia felis* infection in stray and companion cats in northeastern and eastern China. Vector Borne Zoonotic Dis. 16(10):654–658. doi:[10.1089/vbz.2016.1993](https://doi.org/10.1089/vbz.2016.1993).

Karakuş M, Aykur M, Özbel Y, Töz S, Dağcı H. 2016. Molecular detection and genotyping of *Acanthamoeba* spp. among stray dogs using conjunctival swab sampling. Acta Trop. 164:23–26. doi:[10.1016/j.actatropica.2016.08.011](https://doi.org/10.1016/j.actatropica.2016.08.011).

Karatepe B, Babür C, Karatepe M, Kiliç S, Dündar B. 2008. Prevalence of *Toxoplasma gondii* antibodies and intestinal parasites in stray cats from Nigde, Turkey. Ital J Anim Sci. 7(1):113–118. doi:[10.4081/ijas.2008.113](https://doi.org/10.4081/ijas.2008.113).

Karim MR, Dong H, Yu F, Jian F, Zhang L, Wang R, Zhang S, Rume FI, Ning C, Xiao L. 2014. Genetic Diversity in *Enterocytozoon bieneusi* isolates from dogs and cats in china: host specificity and public health implications. J Clin Microbiol. 52(9):3297–3302. doi:[10.1128/JCM.01352-14](https://doi.org/10.1128/JCM.01352-14).

Kasempimolporn S SV Sichanasai B, Saengseesom W, Puempumpanich S, Chatraporn S. 2007. Prevalence of rabies virus infection and rabies antibody in stray dogs: a survey in Bangkok, Thailand. Prev Vet Med. 78(3–4):325–332. doi:[10.1016/j.prevetmed.2006.11.003](https://doi.org/10.1016/j.prevetmed.2006.11.003).

Katagiri S O-STC. 2008. Prevalence of dog intestinal parasites and risk perception of zoonotic infection by dog owners in São Paulo State, Brazil. Zoonoses Public Health. 55(8–10):406–413. doi:[10.1111/j.1863-2378.2008.01163.x](https://doi.org/10.1111/j.1863-2378.2008.01163.x).

Kelly PJ, Köster L, Li J, Zhang J, Huang K, Branford GC, Marchi S, Vandenplas M, Wang C. 2017. Survey of vector-borne agents in feral cats and first report of *Babesia gibsoni* in cats on St Kitts, West Indies. BMC Vet Res. 13:331. doi:[10.1186/s12917-017-1230-1](https://doi.org/10.1186/s12917-017-1230-1).

Kelly PJ, Moura L, Miller T, Thurk J, Perreault N, Weil A, Maggio R, Lucas H, Breitschwerdt E. 2010. Feline Immunodeficiency Virus, Feline Leukemia Virus and *Bartonella* species in stray cats on St Kitts, West Indies. J Feline Med Surg. 12(6):447–450. doi:[10.1016/j.jfms.2009.12.015](https://doi.org/10.1016/j.jfms.2009.12.015).

Ketzis JK, Shell L, Chinault S, Pemberton C, Pereira MM. 2015. The prevalence of *Trichuris* spp. infection in indoor and outdoor cats on St. Kitts. J Infect Dev Ctries. 9(1):111–113. doi:[10.3855/jidc.5778](https://doi.org/10.3855/jidc.5778).

Khalafalla R. 2011. A survey study on gastrointestinal parasites of stray cats in northern region of Nile delta, Egypt. PLoS One. 6(7):e20283. doi:[10.1371/journal.pone.0020283](https://doi.org/10.1371/journal.pone.0020283).

Khamesipour F, Doosti A, Emadi MF, Awosile B. 2014. Detection of *Brucella* sp. and *Leptospira* sp. in dogs using conventional polymerase chain reaction. J Vet Res. 58(4):527–531. doi:[10.2478/bvip-2014-0081](https://doi.org/10.2478/bvip-2014-0081).

Khodabakhsh M, Malmasi A, Mohebali M, Zarei Z, Kia EB, Azarm A. 2016. Feline Dirofilariosis Due to *Dirofilaria immitis* in Meshkin Shahr District, northwestern Iran. Iran J Parasitol. 11(2):269–273.

Kim Y, Seo K, Lee J, Choi E, Lee H, Hwang C, Shin N, Youn H, Youn HY. 2009. Prevalence of *Bartonella henselae* and *Bartonella clarridgeiae* in cats and dogs in Korea. J Vet Sci. 10(1):85–87. doi:[10.4142/jvs.2009.10.1.85](https://doi.org/10.4142/jvs.2009.10.1.85).

Kimura YH Morishima Y, Nagahama S, Horikoshi T, Edagawa A, Kawabuchi-Kurata T, Sugiyama H Kimura, Morishima Y, Nagahama S, Horikoshi T, Edagawa A, Kawabuchi-Kurata T, Sugiyama H, Yamasaki H. 2013. A coprological survey of intestinal helminthes in stray dogs captured in Osaka Prefecture, Japan. J Vet Med Sci. 75(10):1409–1411. doi:[10.1292/jvms.12-0499](https://doi.org/10.1292/jvms.12-0499).

King JS, Brown GK, Jenkins DJ, Ellis JT, Fleming PJS, Windsor PA, Slapeta J. 2012. Oocysts and high seroprevalence of *Neospora caninum* in dogs living in remote Aboriginal communities and wild dogs in Australia. Vet Parasitol. 187(1–2):85–92. doi:[10.1016/j.vetpar.2011.12.027](https://doi.org/10.1016/j.vetpar.2011.12.027).

Kipp EJ, Mariscal J, Armijos RX, Weigel M, Waldrup K. 2016. Genetic evidence of enzootic Leishmaniasis in a stray canine and Texas mouse from sites in west and central Texas. Mem Inst Oswaldo Cruz. 111:652–654. doi:[10.1590/0074-02760160225](https://doi.org/10.1590/0074-02760160225).

Kirby DR. 1979. Prevalence of patent and occult filarial infections in stray dogs from the coastal bend area of Texas. Southwestern veterinarian. 32(2):121-123.

Klimpel S, Heukelbach J, Pothmann D, Rückert S. 2010. Gastrointestinal and ectoparasites from urban stray dogs in Fortaleza (Brazil): high infection risk for humans? Parasitol Res. 107(3):713–719. doi:[10.1007/s00436-010-1926-7](https://doi.org/10.1007/s00436-010-1926-7).

Kocabiyik AL, Cetin C, Dedicova D. 2006. Detection of *Salmonella* spp. in stray dogs in Bursa Province, Turkey: first isolation of *Salmonella corvallis* from dogs. J Vet Med B Infect Dis Vet Public Health. 53(4):194–196. doi:[10.1111/j.1439-0450.2006.00932.x](https://doi.org/10.1111/j.1439-0450.2006.00932.x).

Koh FX, Panchadcharam C, Tay ST. 2016. Vector-Borne diseases in stray dogs in peninsular malaysia and molecular detection of *Anaplasma* and *Ehrlichia* spp. from *Rhipicephalus sanguineus* (Acari: Ixodidae) ticks. J Med Entomol. 53(1):183–187. doi:[10.1093/jme/tjv153](https://doi.org/10.1093/jme/tjv153).

Konto M, Tukur SM, Watanabe M, Abd-Rani P a. M, Sharma RSK, Fong LS. 2017. Molecular and serological prevalence of *Anaplasma* and *Ehrlichia* sp. among stray dogs in East Malaysia. Trop Biomed. 34(3):570–575.

Kozan E, Sevimli F, Birdane FM. 2007. Incidence of Dirofilaria sp. in stray dogs in the Afyonkarahisar and Eskisehir provinces. Veteriner Fakültesi dergisi. 54:117–119.

Kvac M, Hofmannova L, Ortega Y, Holubova N, Horcickova M, Kicia M, Hlaskova L, Kvetonova D, Sak B, McEvoy J. 2017. Stray cats are more frequently infected with zoonotic protists than pet cats. Folia Parasitol (Praha). 64:2017.034. doi:[10.14411/fp.2017.034](https://doi.org/10.14411/fp.2017.034).

Labarthe N, Serrão ML, Ferreira AMR, Almeida NKO, Guerrero J. 2004. A survey of gastrointestinal helminths in cats of the metropolitan region of Rio de Janeiro, Brazil. Vet Parasitol. 123(1–2):133–139. doi:[10.1016/j.vetpar.2004.06.002](https://doi.org/10.1016/j.vetpar.2004.06.002).

Lahmar S, Boufana BS, Lahmar S, Inoubli S, Guadraoui M, Dhibi M, Bradshaw H, Craig PS. 2009. Echinococcus in the wild carnivores and stray dogs of northern Tunisia: the results of a pilot survey. Ann Trop Med Parasitol. 103(4):323–331. doi:[10.1179/136485909X440836](https://doi.org/10.1179/136485909X440836).

Lahmar S, Kilani M, Torgerson PR. 2001. Frequency distributions of *Echinococcus granulos*us and other helminths in stray dogs in Tunisia. Ann Trop Med Parasitol. 95(1):69–76. doi:[10.1080/00034983.2001.11813616](https://doi.org/10.1080/00034983.2001.11813616).

Lan D, Ji W, Yu D, Chu J, Wang C, Yang Z, Hua X. 2011. Serological evidence of West Nile Virus in dogs and cats in China. Arch Virol. 156(5):893–895. doi:[10.1007/s00705-010-0913-8](https://doi.org/10.1007/s00705-010-0913-8).

Lappin MR, Hawley J. 2009. Presence of *Bartonella* species and *Rickettsia* species DNA in the blood, oral cavity, skin and claw beds of cats in the United States. Vet Dermatol. 20(5–6):509–514. doi:[10.1111/j.1365-3164.2009.00800.x](https://doi.org/10.1111/j.1365-3164.2009.00800.x).

Lau SKP, Woo PCY, Yeung HC, Teng JLL, Wu Y, Bai R, Fan RYY, Chan K-H, Yuen K-Y. 2012. Identification and characterization of bocaviruses in cats and dogs reveals a novel feline bocavirus and a novel genetic group of canine bocavirus. J Gen Virol. 93(Pt 7):1573–1582. doi:[10.1099/vir.0.042531-0](https://doi.org/10.1099/vir.0.042531-0).

Lechner ES CR Crawford PC, Levy JK, Edinboro CH, Dubovi EJ. 2010. Prevalence of protective antibody titers for Canine Distemper Virus and Canine Parvovirus in dogs entering a Florida animal shelter. J Am Vet Med Assoc. 236(12):1317–1321. doi:[10.2460/javma.236.12.1317](https://doi.org/10.2460/javma.236.12.1317).

Lee SE LJH Kim NH, Chae HS, Cho SH, Nam HW, Lee WJ, Kim SH. 2011. Prevalence of *Toxoplasma gondii* infection in feral cats in Seoul, Korea. J Parasitol. 97(1):153–155. doi:[10.1645/ge-2455.1](https://doi.org/10.1645/ge-2455.1).

Lefkaditis M, Pastiu A, Rodi-Buriel A, Sossidou A, Panorias A, Eleftheriadis T, Vasile C, Mihalca A. 2014. Helminth burden in stray cats from Thessaloniki, Greece. Helminthologia. 51:73–76. doi:[10.2478/s11687-014-0211-1](https://doi.org/10.2478/s11687-014-0211-1).

Lefkaditis MA, Athanasiou LV, Ionicã AM, Koukeri SE, Panorias A, Eleftheriadis TG, Boutsini S. 2016. Ectoparasite infestations of urban stray dogs in Greece and their zoonotic potential. Trop Biomed. 33(2):226–230.

Lefkaditis MA, Sossidou AV, Panorias AH, Koukeri SE, Paştiu AI, Athanasiou LV. 2015. Urban stray cats infested by ectoparasites with zoonotic potential in Greece. Parasitol Res. 114(10):3931–3934. doi:[10.1007/s00436-015-4688-4](https://doi.org/10.1007/s00436-015-4688-4).

Lehmann C, Lehmann W. 2004. Giardia: Infection and vaccination in an animal shelter. Tierarztliche Umschau. 59:337–340.

Liberato CD, Berrilli F, Odorizi L, Scarcella R, Barni M, Amoruso C, Scarito A, Filippo MMD, Carvelli A, Iacoponi F, et al. 2018. Parasites in stray dogs from Italy: prevalence, risk factors and management concerns. Acta Parasitol. 63(1):27–32. doi:[10.1515/ap-2018-0003](https://doi.org/10.1515/ap-2018-0003).

Lima DCV de, Magalhães FJR, Andrade MR, Silva JG da, Morais EGF de, Filho CDF de L, Porto WJN, Mota RA. 2018. Anti-*Neospora caninum* antibodies in feral cats on the Island of Fernando de Noronha, Brazil. Acta Parasitol. 63(3):645–646. doi:[10.1515/ap-2018-0074](https://doi.org/10.1515/ap-2018-0074).

Lima VFS, Ramos RAN, Lepold R, Borges JCG, Ferreira CD, Rinaldi L, Cringoli G, Alves LC. 2017. Gastrointestinal parasites in feral cats and rodents from the Fernando de Noronha Archipelago, Brazil. Rev Bras Parasitol Vet. 26:521–524. doi:[10.1590/S1984-29612017066](https://doi.org/10.1590/S1984-29612017066).

Little SE. 2005. Feline immunodeficiency virus testing in stray, feral, and client-owned cats of Ottawa. Can Vet J. 46(10):898–901.

Liu J, Song KH, Lee SE, Lee JY, Lee JI, Hayasaki M, You MJ, Kim DH. 2005. Serological and molecular survey of *Dirofilaria immitis* infection in stray cats in Gyunggi province, South Korea. Vet Parasitol. 130(1–2):125–129. doi:[10.1016/j.vetpar.2005.03.026](https://doi.org/10.1016/j.vetpar.2005.03.026).

Liu M, Ruttayaporn N, Saechan V, Jirapattharasate C, Vudriko P, Moumouni PFA, Cao S, Inpankaew T, Ybañez AP, Suzuki H, et al. 2016. Molecular survey of canine vector-borne diseases in stray dogs in Thailand. Parasitol Int. 65(4):357–361. doi:[10.1016/j.parint.2016.04.011](https://doi.org/10.1016/j.parint.2016.04.011).

Liu Y, He G, Cheng Z, Qi Y, Liu J, Zhang H, Liu G, Shi D, Yang D, Wang S, et al. 2012. Seroprevalence of *Toxoplasma gondii* in dogs in Shandong, Henan, and Heilongjiang Provinces, and in the Xinjiang Uygur Autonomous Region, People’s Republic of China. J Parasitol. 98(1):211–212. doi:[10.1645/GE-2892.1](https://doi.org/10.1645/GE-2892.1).

Lonardoni MVC, Bernal FHZ, Silveira TGV, Antunes V, Teodoro U, Jorge FA, Zanzarini PD. 2006. Comparison between indirect immunofluorescence and direct agglutination for the serologic diagnosis of American Cutaneous Leishmaniasis in stray dogs. Arq Bras Med Vet Zootec. 58:1001–1008. doi:[10.1590/S0102-09352006000600005](https://doi.org/10.1590/S0102-09352006000600005).

Luria BJ, Levy JK, Lappin MR, Breitschwerdt EB, Legendre AM, Hernandez JA, Gorman SP, Lee IT. 2004. Prevalence of infectious diseases in feral cats in Northern Florida. J Feline Med Surg. 6(5):287–296. doi:[10.1016/j.jfms.2003.11.005](https://doi.org/10.1016/j.jfms.2003.11.005).

Lyoo KS HTW Kim D, Jang HG, Lee SJ, Park MY. 2017. Prevalence of antibodies against *Coxiella burnetii* in Korean native cattle, dairy cattle, and dogs in South Korea. Vector Borne Zoonotic Dis. 17(3):213–216. doi:[10.1089/vbz.2016.1977](https://doi.org/10.1089/vbz.2016.1977).

Maden M, Doğan, Altintaş G, Yildiz E, Ekik M, Ince ME, Köse S. 2015. Prevalence of *Bartonella henselae* in pet and stray cats from the aspect of public health: a research sample in the concept of One Medicine -One Health. Kafkas Univ Vet Fak Derg. 21:313–317. doi:[10.9775/kvfd.2014.12371](https://doi.org/10.9775/kvfd.2014.12371).

Magalhães FJR, Ribeiro-Andrade M, Souza FM, Lima Filho CDF, Biondo AW, Vidotto O, Navarro IT, Mota RA. 2017. Seroprevalence and spatial distribution of *Toxoplasma gondii* infection in cats, dogs, pigs and equines of the Fernando de Noronha Island, Brazil. Parasitol Int. 66(2):43–46. doi:[10.1016/j.parint.2016.11.014](https://doi.org/10.1016/j.parint.2016.11.014).

Magalhaes J, Sicupira P, Munhoz A. 2009. Frequency of gastrointestinal parasites in dogs at municipality of Ilheus in the state of Bahia. Revista brasileira de medicina veterinaria. 31:151–156.

Mahdy MA SJ Lim YA, Ngui R, Siti Fatimah MR, Choy SH, Yap NJ, Al-Mekhlafi HM, Ibrahim J. 2012. Prevalence and zoonotic potential of canine hookworms in Malaysia. Parasit Vectors. 5:88. doi:[10.1186/1756-3305-5-88](https://doi.org/10.1186/1756-3305-5-88).

Maia C, Almeida B, Coimbra M, Fernandes MC, Cristóvão JM, Ramos C, Martins Â, Martinho F, Silva P, Neves N, et al. 2015. Bacterial and protozoal agents of canine vector-borne diseases in the blood of domestic and stray dogs from southern Portugal. Parasit Vectors. 8:138. doi:[10.1186/s13071-015-0759-8](https://doi.org/10.1186/s13071-015-0759-8).

Maia C CL Cortes H, Brancal H, Lopes AP, Pimenta P, Campino L. 2014. Prevalence and correlates of antibodies to *Neospora caninum* in dogs in Portugal. Parasite. 21:29. doi:[10.1051/parasite/2014031](https://doi.org/10.1051/parasite/2014031).

Maia C, Nunes M, Campino L. 2008. Importance of cats in zoonotic Leishmaniasis in Portugal. Vector Borne Zoonotic Dis. 8(4):555–559. doi:[10.1089/vbz.2007.0247](https://doi.org/10.1089/vbz.2007.0247).

Maia C, Ramos C, Coimbra M, Bastos F, Martins Â, Pinto P, Nunes M, Vieira ML, Cardoso L, Campino L. 2014. Bacterial and protozoal agents of feline vector-borne diseases in domestic and stray cats from southern Portugal. Parasites & Vectors. 7(1):115. doi:[10.1186/1756-3305-7-115](https://doi.org/10.1186/1756-3305-7-115).

Maleky F, Moradkhan M. 2000. Echinococcosis in the stray dogs of Tehran, Iran. Ann Trop Med Parasitol. 94(4):329–331. doi:[10.1080/00034983.2000.11813547](https://doi.org/10.1080/00034983.2000.11813547).

Malgor R, Nonaka N, Basmadjian I, Sakai H, Carámbula B, Oku Y, Carmona C, Kamiya M. 1997. Coproantigen detection in dogs experimentally and naturally infected with *Echinococcus granulosus* by a monoclonal antibody-based enzyme-linked immunosorbent assay. Int J Parasitol. 27(12):1605–1612. doi:[10.1016/s0020-7519(97)00127-6](https://doi.org/10.1016/s0020-7519(97)00127-6).

Malgor R, Oku Y, Gallardo R, Yarzábal I. 1996. High prevalence of *Ancylostoma* spp. infection in dogs, associated with endemic focus of human Cutaneous Larva Migrans, in Tacuarembo, Uruguay. Parasite. 3(2):131–134. doi:[10.1051/parasite/1996032131](https://doi.org/10.1051/parasite/1996032131).

Malmasi A, Janitabar S, Mohebali M, Akhoundi B, Maazi N, Aramoon M, Khorrami N, Seifi HA. 2014. Seroepidemiologic survey of canine Visceral Leishmaniasis in Tehran and Alborz Provinces of Iran. J Arthropod Borne Dis. 8(2):132–138.

Mancianti F, Nardoni S, Ariti G, Parlanti D, Giuliani G, Papini RA. 2010. Cross-sectional survey of *Toxoplasma gondii* infection in colony cats from urban Florence (Italy). J Feline Med Surg. 12(4):351–354. doi:[10.1016/j.jfms.2009.09.001](https://doi.org/10.1016/j.jfms.2009.09.001).

Manić M. 2015. Seroepidemiological survey of leptospiral infection in stray dogs in Serbia. Turkish Vet Anim Sci. 39:719–723.

Manyarara R, Tubbesing U, Soni M, Noden BH. 2015. Serodetection of *Ehrlichia canis* amongst dogs in central Namibia. J S Afr Vet Assoc. 86(1):1272. doi:[10.4102/jsava.v86i1.1272](https://doi.org/10.4102/jsava.v86i1.1272).

Mark-Carew MP, Adesiyun AA, Basu A, Georges KA, Pierre T, Tilitz S, Wade SE, Mohammed HO. 2013. Characterization of *Giardia duodenalis* infections in dogs in Trinidad and Tobago. Vet Parasitol. 196(1–2):199–202. doi:[10.1016/j.vetpar.2013.01.023](https://doi.org/10.1016/j.vetpar.2013.01.023).

Markovich JE, Ross L, McCobb E. 2012. The prevalence of leptospiral antibodies in free roaming cats in Worcester County, Massachusetts. J Vet Intern Med. 26(3):688–689. doi:[10.1111/j.1939-1676.2012.00900.x](https://doi.org/10.1111/j.1939-1676.2012.00900.x).

Martínez-Barbabosa I GE Quiroz MG, González LA, Cárdenas EM, Edubiel AA, Juárez JL. 2008. Prevalence of anti-*T. canis* antibodies in stray dogs in Mexico City. Vet Parasitol. 153(3–4):270–276. doi:[10.1016/j.vetpar.2008.02.011](https://doi.org/10.1016/j.vetpar.2008.02.011).

Matei IA, D’Amico G, Yao PK, Ionică AM, Kanyari PWN, Daskalaki AA, Dumitrache MO, Sándor AD, Gherman CM, Qablan M, et al. 2016. Molecular detection of *Anaplasma platys* infection in free-roaming dogs and ticks from Kenya and Ivory Coast. Parasites Vectors. 9(1):157. doi:[10.1186/s13071-016-1443-3](https://doi.org/10.1186/s13071-016-1443-3).

Matsuu A, Yokota S-I, Ito K, Masatani T. 2017. Seroprevalence of *Toxoplasma gondii* in free-ranging and feral cats on Amami Oshima Island, Japan. J Vet Med Sci. 79(11):1853–1856. doi:[10.1292/jvms.17-0359](https://doi.org/10.1292/jvms.17-0359).

McGlade TR, Robertson ID, Elliot AD, Read C, Thompson RCA. 2003. Gastrointestinal parasites of domestic cats in Perth, Western Australia. Vet Parasitol. 117(4):251–262. doi:[10.1016/j.vetpar.2003.08.010](https://doi.org/10.1016/j.vetpar.2003.08.010).

McManus CM TSJ Levy JK, Andersen LA, McGorray SP, Leutenegger CM, Gray LK, Hilligas J. 2014. Prevalence of upper respiratory pathogens in four management models for unowned cats in the Southeast United States. Vet J. 201(2):196–201. doi:[10.1016/j.tvjl.2014.05.015](https://doi.org/10.1016/j.tvjl.2014.05.015).

McQuiston JH, Guerra MA, Watts MR, Lawaczeck E, Levy C, Nicholson WL, Adjemian J, Swerdlow DL. 2011. Evidence of exposure to Spotted Fever Group Rickettsiae among Arizona dogs outside a previously documented outbreak area. Zoonoses Public Health. 58(2):85–92. doi:[10.1111/j.1863-2378.2009.01300.x](https://doi.org/10.1111/j.1863-2378.2009.01300.x).

Mehrabani D, Sadjjadi SM, Oryan A. 2002. Prevalence of gastrointestinal nematode parasites in stray dogs in Shiraz, southern Iran. J Appl Anim Res. 22(1):157–160. doi:[10.1080/09712119.2002.9706391](https://doi.org/10.1080/09712119.2002.9706391).

Mehrabani D SSM Oryan A, Oryan A. 1999. Prevalence of *Echinococcus granulosus* infection in stray dogs and herbivores in Shiraz, Iran. Vet Parasitol. 86(3):217–220. doi:[10.1016/s0304-4017(99)00151-x](https://doi.org/10.1016/s0304-4017(99)00151-x).

Meira CD, Wenceslau AA, Carvalho FS, Albuquerque GR, Dias RC. 2011. Diagnóstico molecular de Leptospirose no sangue de cães naturalmente infectados. Bras J Vet Med. 33(1):7–11.

Melo LC MJY Oresco C, Leigue L, Netto HM, Melville PA, Benites NR, Saras E, Haenni M, Lincopan N. 2018. Prevalence and molecular features of ESBL/pAmpC-producing Enterobacteriaceae in healthy and diseased companion animals in Brazil. Vet Microbiol. 221:59–66. doi:[10.1016/j.vetmic.2018.05.017](https://doi.org/10.1016/j.vetmic.2018.05.017).

Melo RPB, Almeida JC, Lima DCV, Pedrosa CM, Magalhães FJR, Alcântara AM, Barros LD, Vieira RFC, Garcia JL, Mota RA. 2016. Atypical *Toxoplasma gondii* genotype in feral cats from the Fernando de Noronha Island, northeastern Brazil. Vet Parasitol. 224:92–95. doi:[10.1016/j.vetpar.2016.05.023](https://doi.org/10.1016/j.vetpar.2016.05.023).

Melo SN BVS Teixeira-Neto RG, Werneck GL, Struchiner CJ, Ribeiro RAN, Sousa LR, de Melo MOG, Carvalho Júnior CG, Penaforte KM, Manhani MN, Aquino VV, Silva ES. 2018. Prevalence of Visceral Leishmaniasis in A population of free-roaming dogs as determined by multiple sampling efforts: A longitudinal study analyzing the effectiveness of euthanasia. Prev Vet Med. 161:19–24. doi:[10.1016/j.prevetmed.2018.10.010](https://doi.org/10.1016/j.prevetmed.2018.10.010).

Mendes-de-Almeida F, Labarthe N, Guerrero J, Faria MCF, Branco AS, Pereira CD, Barreira JD, Pereira MJS. 2007. Follow-up of the health conditions of an urban colony of free-roaming cats (*Felis catus* Linnaeus, 1758) in the city of Rio de Janeiro, Brazil. Vet Parasitol. 147(1):9–15. doi:[10.1016/j.vetpar.2007.03.035](https://doi.org/10.1016/j.vetpar.2007.03.035).

Meneses IDS de, Andrade MR, Uzêda RS, Bittencourt MV, Lindsay DS, Gondim LFP, Meneses IDS de, Andrade MR, Uzêda RS, Bittencourt MV, et al. 2014. Frequency of antibodies against *Sarcocystis neurona* and *Neospora caninum* in domestic cats in the state of Bahia, Brazil. Rev Bras Parasitol Vet. 23(4):526–529. doi:[10.1590/S1984-29612014080](https://doi.org/10.1590/S1984-29612014080).

Meshgi B AO. 2003. Prevalence of *Linguatula serrata* infestation in stray dogs of Shahrekord, Iran. J Vet Med B Infect Dis Vet Public Health. 50(9):466–467. doi:[10.1046/j.0931-1793.2003.00705.x](https://doi.org/10.1046/j.0931-1793.2003.00705.x).

Millán J, Cabezón O, Pabón M, Dubey JP, Almería S. 2009. Seroprevalence of *Toxoplasma gondii* and *Neospora caninum* in feral cats (*Felis silvestris catus*) in Majorca, Balearic Islands, Spain. Vet Parasitol. 165(3–4):323–326. doi:[10.1016/j.vetpar.2009.07.014](https://doi.org/10.1016/j.vetpar.2009.07.014).

Millán J, Candela MG, López-Bao JV, Pereira M, Jiménez MA, León-Vizcaíno L. 2009. Leptospirosis in wild and domestic carnivores in natural areas in Andalusia, Spain. Vector Borne Zoonotic Dis. 9(5):549–554. doi:[10.1089/vbz.2008.0081](https://doi.org/10.1089/vbz.2008.0081).

Millán J, Casanova JC. 2009. High prevalence of helminth parasites in feral cats in Majorca Island (Spain). Parasitol Res. 106(1):183–188. doi:[10.1007/s00436-009-1647-y](https://doi.org/10.1007/s00436-009-1647-y).

Millán J, Sobrino R, Rodríguez A, Oleaga A, Gortazar C, Schares G. 2012. Large-scale serosurvey of *Besnoitia besnoiti* in free-living carnivores in Spain. Vet Parasitol. 190(1–2):241–245. doi:[10.1016/j.vetpar.2012.06.014](https://doi.org/10.1016/j.vetpar.2012.06.014).

Millán J, Zanet S, Gomis M, Trisciuoglio A, Negre N, Ferroglio E. 2011. An investigation into alternative reservoirs of canine leishmaniasis on the endemic island of Mallorca (Spain). Transbound Emerg Dis. 58(4):352–357. doi:[10.1111/j.1865-1682.2011.01212.x](https://doi.org/10.1111/j.1865-1682.2011.01212.x).

Milstein TC, Goldsmid JM. 1997. Parasites of feral cats from southern Tasmania and their potential significance. Aust Vet J. 75(3):218–219. doi:[10.1111/j.1751-0813.1997.tb10072.x](https://doi.org/10.1111/j.1751-0813.1997.tb10072.x).

Minnaar WN, Krecek RC, Fourie LJ. 2002. Helminths in dogs from a peri-urban resource-limited community in Free State Province, South Africa. Vet Parasitol. 107(4):343–349. doi:[10.1016/s0304-4017(02)00155-3](https://doi.org/10.1016/s0304-4017(02)00155-3).

Mirbadie SR, Kamyabi H, Mohammadi MA, Shamsaddini S, Harandi MF. 2018. Copro-PCR prevalence of *Echinococcus granulosus* infection in dogs in Kerman, south-eastern Iran. J Helminthol. 92(1):17–21. doi:[10.1017/S0022149X17000074](https://doi.org/10.1017/S0022149X17000074).

Mircean V, Dumitrache MO, Györke A, Pantchev N, Jodies R, Mihalca AD, Cozma V. 2012. Seroprevalence and geographic distribution of *Dirofilaria immitis* and tick-borne infections (*Anaplasma phagocytophilum*, *Borrelia burgdorferi* sensu lato, and *Ehrlichia canis*) in dogs from Romania. Vector Borne Zoonotic Dis. 12(7):595–604. doi:[10.1089/vbz.2011.0915](https://doi.org/10.1089/vbz.2011.0915).

Miró G, Checa R, Montoya A, Hernández L, Dado D, Gálvez R. 2012. Current situation of *Leishmania infantum* infection in shelter dogs in northern Spain. Parasit Vectors. 5:60. doi:[10.1186/1756-3305-5-60](https://doi.org/10.1186/1756-3305-5-60).

Miró G FI Montoya A, Jiménez S, Frisuelos C, Mateo M. 2004. Prevalence of antibodies to *Toxoplasma gondii* and intestinal parasites in stray, farm and household cats in Spain. Vet Parasitol. 126(3):249–255. doi:[10.1016/j.vetpar.2004.08.015](https://doi.org/10.1016/j.vetpar.2004.08.015).

Miró G, Mateo M, Montoya A, Vela E, Calonge R. 2007. Survey of intestinal parasites in stray dogs in the Madrid area and comparison of the efficacy of three anthelmintics in naturally infected dogs. Parasitol Res. 100(2):317–320. doi:[10.1007/s00436-006-0258-0](https://doi.org/10.1007/s00436-006-0258-0).

Miró G, Müller A, Montoya A, Checa R, Marino V, Marino E, Fuster F, Escacena C, Descalzo MA, Gálvez R. 2017. Epidemiological role of dogs since the human Leishmaniosis outbreak in Madrid. Parasit Vectors. 10:209. doi:[10.1186/s13071-017-2147-z](https://doi.org/10.1186/s13071-017-2147-z).

Miró G, Rupérez C, Checa R, Gálvez R, Hernández L, García M, Canorea I, Marino V, Montoya A. 2014. Current status of *L. infantum* infection in stray cats in the Madrid region (Spain): implications for the recent outbreak of human Leishmaniosis? Parasit Vectors. 7:112. doi:[10.1186/1756-3305-7-112](https://doi.org/10.1186/1756-3305-7-112).

Mirzaei M. 2012. Epidemiological survey of *Cryptosporidium* spp. in companion and stray dogs in Kerman, Iran. Vet Ital. 48(3):291–296.

Mitrea IL, Enachescu V, Ionita M. 2013. *Neospora caninum* infection in dogs from Southern Romania: coproparasitological study and serological follow-up. J Parasitol. 99(2):365–367. doi:[10.1645/GE-3230.1](https://doi.org/10.1645/GE-3230.1).

Mohd Zain SN, Sahimin N, Pal P, Lewis JW. 2013. Macroparasite communities in stray cat populations from urban cities in Peninsular Malaysia. Vet Parasitol. 196(3–4):469–477. doi:[10.1016/j.vetpar.2013.03.030](https://doi.org/10.1016/j.vetpar.2013.03.030).

Mohsen A, Hossein H. 2009. Gastrointestinal parasites of stray cats in Kashan, Iran. Trop Biomed. 26(1):16–22.

Molan A. 1993. Epidemiology of Hydatidosis and Echinococcosis in Theqar Province, southern Iraq. Jpn J Med Sci Biol. 46(1):29–35. doi:[10.7883/yoken1952.46.29](https://doi.org/10.7883/yoken1952.46.29).

Molan A, Baban M. 1992. The prevalence of *Echinococcus granulosus* in stray dogs in Iraq. J Trop Med Hyg. 95(2):146–148.

Mtambo MM, Nash AS, Blewett DA, Smith HV, Wright S. 1991. *Cryptosporidium* infection in cats: prevalence of infection in domestic and feral cats in the Glasgow area. Vet Rec. 129(23):502–504.

Mtambo MM, Nash AS, Wright SE, Smith HV, Blewett DA, Jarrett O. 1995. Prevalence of specific anti-*Cryptosporidium* IgG, IgM and IgA antibodies in cat sera using an indirect immunofluorescence antibody test. Vet Parasitol. 60(1–2):37–43. doi:[10.1016/0304-4017(94)00749-3](https://doi.org/10.1016/0304-4017(94)00749-3).

Muirden A. 2002. Prevalence of Feline Leukaemia Virus and antibodies to Feline Immunodeficiency Virus and Feline Coronavirus in stray cats sent to an RSPCA hospital. Vet Rec. 150(20):621–625. doi:[10.1136/vr.150.20.621](https://doi.org/10.1136/vr.150.20.621).

Mukaratirwa S SVP. 2010. Prevalence of gastrointestinal parasites of stray dogs impounded by the Society for the Prevention of Cruelty to Animals (SPCA), Durban and Coast, South Africa. J S Afr Vet Assoc. 81(2):123–125. doi:[10.4102/jsava.v81i2.124](https://doi.org/10.4102/jsava.v81i2.124).

Müller S, Boulouis H, Viallard J, Beugnet F. 2004. Epidemiological survey of canine Bartonellosis to *Bartonella vinsonii subs. berkhoffii* and Canine Monocytic Ehrlichiosis in dogs on the Island of Reunion. Rev Med Vet. 155:377–380.

Mundim MJ CMC Rosa LA, Hortêncio SM, Faria ES, Rodrigues RM. 2007. Prevalence of *Giardia duodenalis* and *Cryptosporidium* spp. in dogs from different living conditions in Uberlândia, Brazil. Vet Parasitol. 144(3–4):356–359. doi:[10.1016/j.vetpar.2006.09.039](https://doi.org/10.1016/j.vetpar.2006.09.039).

Myers DM, Varela-Díaz VM. 1980. Serological and bacteriological detection of *Brucella canis* infection of stray dogs in Moreno, Argentina. Cornell Vet. 70(3):258–265.

Nava AFD, Cullen L, Sana DA, Nardi MS, Filho JDR, Lima TF, Abreu KC, Ferreira F. 2008. First evidence of Canine Distemper in Brazilian free-ranging felids. Ecohealth. 5(4):513–518. doi:[10.1007/s10393-008-0207-8](https://doi.org/10.1007/s10393-008-0207-8).

Nazir MM, Maqbool A, Akhtar M, Ayaz M, Ahmad AN, Ashraf K, Ali A, Alam MA, Ali MA, Khalid AR, et al. 2014. *Neospora caninum* prevalence in dogs raised under different living conditions. Vet Parasitol. 204(3–4):364–368. doi:[10.1016/j.vetpar.2014.05.041](https://doi.org/10.1016/j.vetpar.2014.05.041).

Ng KL, Lee EL, Sani RA. 2012. Low prevalence of *Dirofilaria immitis* in dogs in Johor Bahru, Malaysia as a reflection of vector availability? Trop Biomed. 29(1):187–190.

Nguyen TT-D, Choe S-E, Byun J-W, Koh H-B, Lee H-S, Kang S-W. 2012. Seroprevalence of *Toxoplasma gondii* and *Neospora caninum* in dogs from Korea. Acta Parasitol. 57(1):7–12. doi:[10.2478/s11686-012-0010-0](https://doi.org/10.2478/s11686-012-0010-0).

Nichol S SKR Ball SJ. 1981. Prevalence of intestinal parasites in feral cats in some urban areas of England. Vet Parasitol. 9(2):107–110. doi:[10.1016/0304-4017(81)90028-5](https://doi.org/10.1016/0304-4017(81)90028-5).

Nicholson WL, Gordon R, Demma LJ. 2006. Spotted fever group rickettsial infection in dogs from eastern Arizona: how long has it been there? Ann N Y Acad Sci. 1078:519–522. doi:[10.1196/annals.1374.102](https://doi.org/10.1196/annals.1374.102).

Nikolić I, Dimitrijević M, Đurković-Đaković M, Bobic B, Maksimović-Mihajlović O. 2002. Giardiasis in dogs and cats in the Belgrade area. Acta Vet. 52(1):43-48. doi:[10.2298/AVB0201043N](https://doi.org/10.2298/AVB0201043N).

Normand CM, Urbanek RE. 2017. Exurban feral cat seroprevalence of Feline Leukemia and Feline Immunodeficiency Viruses and adult survival. sena. 16(1):1–18. doi:[10.1656/058.016.0102](https://doi.org/10.1656/058.016.0102).

Norris JM, Bell ET, Hales L, Toribio J-ALML, White JD, Wigney DI, Baral RM, Malik R. 2007. Prevalence of Feline Immunodeficiency Virus infection in domesticated and feral cats in eastern Australia. J Feline Med Surg. 9(4):300–308. doi:[10.1016/j.jfms.2007.01.007](https://doi.org/10.1016/j.jfms.2007.01.007).

Nutter FB, Dubey JP, Levine JF, Breitschwerdt EB, Ford RB, Stoskopf MK. 2004. Seroprevalences of antibodies against *Bartonella henselae* and *Toxoplasma gondii* and fecal shedding of *Cryptosporidium* spp, *Giardia* spp, and *Toxocara cati* in feral and pet domestic cats. J Am Vet Med Assoc. 225(9):1394–1398. doi:[10.2460/javma.2004.225.1394](https://doi.org/10.2460/javma.2004.225.1394).

Obaidat MM, Alshehabat MA. 2018. Zoonotic *Anaplasma phagocytophilum*, *Ehrlichia canis*, *Dirofilaria immitis*, *Borrelia burgdorferi*, and Spotted Fever Group Rickettsiae (SFGR) in different types of dogs. Parasitol Res. 117(11):3407–3412. doi:[10.1007/s00436-018-6033-1](https://doi.org/10.1007/s00436-018-6033-1).

O’Callaghan M, Reddin J, Lehmann D. 2005. Helminth and protozoan parasites of feral cats from Kangaroo Island. Trans R Soc S Aust. 129:81–83.

O’Dwyer LH, Saito ME, Hasegawa MY, Kohayagawa A. 2006. Prevalence, hematology and serum biochemistry in stray dogs naturally infected by *Hepatozoon canis* in São Paulo. Arq Bras Med Vet Zootec. 58:688–690. doi:[10.1590/S0102-09352006000400039](https://doi.org/10.1590/S0102-09352006000400039).

Ogunkoya AB, Beran GW, Umoh JU, Gomwalk NE, Abdulkadir IA. 1990. Serological evidence of infection of dogs and man in Nigeria by lyssaviruses (family Rhabdoviridae). Trans R Soc Trop Med Hyg. 84(6):842–845. doi:[10.1016/0035-9203(90)90103-l](https://doi.org/10.1016/0035-9203(90)90103-l).

Ojha KC, Singh DK, Kaphle K, Shah Y, Pant DK. 2018. Sero-prevalence of Leptospirosis and differentiation in blood parameters between positive and negative cases in dogs of Kathmandu Valley. Trans R Soc Trop Med Hyg. 112(8):378–382. doi:[10.1093/trstmh/try065](https://doi.org/10.1093/trstmh/try065).

Okoye IC, Obiezue NR, Okorie CE, Ofoezie IE. 2011. Epidemiology of intestinal helminth parasites in stray dogs from markets in south-eastern Nigeria. J Helminthol. 85(4):415–420. doi:[10.1017/S0022149X10000738](https://doi.org/10.1017/S0022149X10000738).

Okumu TA, Munene JN, Wabacha J, Tsuma V, Leeuwen JV. 2016. Seroepidemiological survey of *Neospora caninum* and its risk factors in farm dogs in Nakuru district, Kenya. Vet World. 9(10):1162–1166. doi:[10.14202/vetworld.2016.1162-1166](https://doi.org/10.14202/vetworld.2016.1162-1166).

Oliveira-Sequeira TC NLC Amarante AF, Ferrari TB. 2002. Prevalence of intestinal parasites in dogs from São Paulo State, Brazil. Vet Parasitol. 103(1–2):19–27. doi:[10.1016/s0304-4017(01)00575-1](https://doi.org/10.1016/s0304-4017(01)00575-1).

O’Lorcain P. 1994. Epidemiology of *Toxocara* spp. in stray dogs and cats in Dublin, Ireland. J Helminthol. 68(4):331–336. doi:[10.1017/s0022149x00001590](https://doi.org/10.1017/s0022149x00001590).

Olsen CS, Willesen JL, Pipper CB, Mejer H. 2015. Occurrence of *Aelurostrongylus abstrusus* (Railliet, 1898) in Danish cats: A modified lung digestion method for isolating adult worms. Vet Parasitol. 210(1–2):32–39. doi:[10.1016/j.vetpar.2015.03.016](https://doi.org/10.1016/j.vetpar.2015.03.016).

Oncel T, Handemir E, Kamburgil K, Yurtalan S. 2007. Determination of Seropositivity for *Toxoplasma gondii* in Stray Dogs in Istanbul, Turkey. Rev Med Vet. 158:223-228.

Oncel T, Vural G. 2005. Seroprevalence of *Dirofilaria immitis* in Stray Dogs in İstanbul and Üzmir. Turkish J Vet Anim. 29:785–789.

Orkun O, Koc N, Sursal N, Cakmak A, Nalbantoğlu S, Karaer Z. 2018. Molecular characterization of tick-borne blood protozoa in stray dogs from Central Anatolia Region of Turkey with a High-Rate *Hepatozoon* Infection. Kafkas Univ Vet Fak Derg. 24:227–232. doi:[10.9775/kvfd.2017.18678](https://doi.org/10.9775/kvfd.2017.18678).

Ortega‐Pacheco A, Guzmán‐Marín E, Acosta‐Viana KY, Vado‐Solís I, Jiménez‐Delgadillo B, Cárdenas‐Marrufo M, Pérez‐Osorio C, Puerto‐Solís M, Jiménez‐Coello M. 2017. Serological survey of *Leptospira interrogans*, *Toxoplasma gondii* and *Trypanosoma cruzi* in free roaming domestic dogs and cats from a marginated rural area of Yucatan Mexico. Vet Med Sci. 3(1):40–47. doi:[10.1002/vms3.55](https://doi.org/10.1002/vms3.55).

Ortuño A, Gauss CBL, García F, Gutierrez JF. 2005. Serological evidence of *Ehrlichia* spp. exposure in cats from northeastern Spain. J Vet Med B Infect Dis Vet Public Health. 52(5):246–248. doi:[10.1111/j.1439-0450.2005.00849.x](https://doi.org/10.1111/j.1439-0450.2005.00849.x).

Oryan A, Sadjjadi S, Mehrabani D, Kargar M. 2008. Spirocercosis and its complications in stray dogs in Shiraz, Southern Iran. Veterinarni Medicina. 53(11):617-624. doi:[10.17221/1866-VETMED](https://doi.org/10.17221/1866-VETMED).

OSullivan EN. 1997. Helminth infections in owned and stray dogs in Co Cork, Ireland. Irish Vet J. 50(2):108–110.

Öter K, Bilgin Z, Tinar R, Tüzer E. 2011. Tapeworm infections in stray dogs and cats in İstanbul, Turkey. Kafkas Univ Vet Fak Derg. 17:595-599.

Palmer MV, Stoffregen WC, Carpenter JG, Stabel JR. 2005. Isolation of *Mycobacterium avium* subsp paratuberculosis (Map) from feral cats on a dairy farm with Map-infected cattle. J Wildl Dis. 41(3):629–635. doi:[10.7589/0090-3558-41.3.629](https://doi.org/10.7589/0090-3558-41.3.629).

Pandey VS, Dakkak A, Elmamoune M. 1987. Parasites of stray dogs in the Rabat region, Morocco. Ann Trop Med Parasitol. 81(1):53–55. doi:[10.1080/00034983.1987.11812090](https://doi.org/10.1080/00034983.1987.11812090).

Pardo A, Pérez C, Góngora A, Gómez L, Moreno A. 2009 May 1. Encuesta exploratoria de infección por *Brucella canis* en perros de Villavicencio - Colombia. Rev MVZ Córdoba. 14(2):1690-1696. doi:[10.21897/rmvz.352](https://doi.org/10.21897/rmvz.352).

Parkes JP, Heyward RP, Henning J, Motha MXJ. 2004. Antibody responses to rabbit haemorrhagic disease virus in predators, scavengers, and hares in New Zealand during epidemics in sympatric rabbit populations. N Z Vet J. 52(2):85–89. doi:[10.1080/00480169.2004.36410](https://doi.org/10.1080/00480169.2004.36410).

de Paula Dreer MK, Gonçalves DD, da Silva Caetano IC, Gerônimo E, Menegas PH, Bergo D, Ruiz Lopes-Mori FM, Benitez A, de Freitas JC, Evers F, et al. 2013. Toxoplasmosis, Leptospirosis and Brucellosis in stray dogs housed at the shelter in Umuarama municipality, Paraná, Brazil. J Venom Anim Toxins Incl Trop Dis. 19:23. doi:[10.1186/1678-9199-19-23](https://doi.org/10.1186/1678-9199-19-23).

Payo-Puente P R-VFA Botelho-Dinis M, Carvaja Urueña AM, Payo-Puente M, Gonzalo-Orden JM. 2008. Prevalence study of the lungworm *Aelurostrongylus abstrusus* in stray cats of Portugal. J Feline Med Surg. 10(3):242–246. doi:[10.1016/j.jfms.2007.12.002](https://doi.org/10.1016/j.jfms.2007.12.002).

Paz e Silva FM, E Silva P, Monobe MM, Lopes RS, Araujo JP. 2012. Molecular characterization of *Giardia duodenalis* in dogs from Brazil. Parasitol Res. 110(1):325–334. doi:[10.1007/s00436-011-2492-3](https://doi.org/10.1007/s00436-011-2492-3).

Pellizzonil S, Sicupira P, Carlos R, Lopes C, Albuquerque G. 2009. Ocorrence of antibodies anti-*Toxoplasma* *gondii* in dogs seized by the Ilheus Center For Zoonosis Control, BA, BRAZIL. Rev Bras Med Vet. 31:9–12.

Pereira PF, Barbosa A da S, Moura APP de, Vasconcellos ML, Uchôa CMA, Bastos OMP, Amendoeira MRR. 2017. Gastrointestinal parasites in stray and shelter cats in the municipality of Rio de Janeiro, Brazil. Rev Bras Parasitol Vet. 26(3):383–388. doi:[10.1590/S1984-29612017024](https://doi.org/10.1590/S1984-29612017024).

Perera P, Rajapakse R, Rajakaruna (Amarakoon) R. 2013. Gastrointestinal parasites of dogs in Hantana area in the Kandy District †. Journal of the National Science Foundation of Sri Lanka. 41:81.

Ponce-Macotela M, Peralta-Abarca GE, Martínez-Gordillo MN. 2005. *Giardia intestinalis* and other zoonotic parasites: prevalence in adult dogs from the southern part of Mexico City. Vet Parasitol. 131(1–2):1–4. doi:[10.1016/j.vetpar.2005.03.027](https://doi.org/10.1016/j.vetpar.2005.03.027).

Qablan MA, Kubelová M, Siroký P, Modrý D, Amr ZS. 2012. Stray dogs of northern Jordan as reservoirs of ticks and tick-borne hemopathogens. Parasitol Res. 111(1):301–307. doi:[10.1007/s00436-012-2839-4](https://doi.org/10.1007/s00436-012-2839-4).

Qi M, Dong H, Wang R, Li J, Zhao J, Zhang L, Luo J. 2016. Infection rate and genetic diversity of *Giardia duodenalis* in pet and stray dogs in Henan Province, China. Parasitol Int. 65(2):159–162. doi:[10.1016/j.parint.2015.11.008](https://doi.org/10.1016/j.parint.2015.11.008).

Qian W, Wang H, Su C, Shan D, Cui X, Yang N, Lv C, Liu Q. 2012. Isolation and characterization of *Toxoplasma gondii* strains from stray cats revealed a single genotype in Beijing, China. Vet Parasitol. 187(3–4):408–413. doi:[10.1016/j.vetpar.2012.01.026](https://doi.org/10.1016/j.vetpar.2012.01.026).

Qiu Y, Nakao R, Thu MJ, Akter S, Alam MZ, Kato S, Katakura K, Sugimoto C. 2016. Molecular evidence of Spotted Fever Group Rickettsiae and Anaplasmataceae from ticks and stray dogs in Bangladesh. Parasitol Res. 115(3):949–955. doi:[10.1007/s00436-015-4819-y](https://doi.org/10.1007/s00436-015-4819-y).

Raab O, Greenwood S, Vanderstichel R, Gelens H. 2016. A cross-sectional study of *Tritrichomonas foetus* infection in feral and shelter cats in Prince Edward Island, Canada. Can Vet J. 57(3):265–270.

Radwan N, Khalil A, Elmahy R. 2009. Morphology and Occurrence of Species of *Toxocara* in Wild Mammal Populations from Egypt. Comp Parasitol. 76:273–282. doi:[10.1654/4367.1](https://doi.org/10.1654/4367.1).

Ragg JR, Moller H, Waldrup KA. 1995. The prevalence of Bovine Tuberculosis (*Mycobacterium bovis*) infections in feral populations of cats (*Felis catus*), ferrets (*Mustela furo*) and stoats (*Mustela erminea*) in Otago and Southland, New Zealand. N Z Vet J. 43(7):333–337. doi:[10.1080/00480169./1995.35915](https://doi.org/10.1080/00480169./1995.35915).

Ramos D, Zocco B, Torres M, Braga Í, Pacheco R, Sinkoc A. 2015. Helminths parasites of stray dogs (*Canis lupus familiaris*) from Cuiabá, Midwestern of Brazil. Sem Cienc Agrar. 36:889. doi:[10.5433/1679-0359.2015v36n2p889](https://doi.org/10.5433/1679-0359.2015v36n2p889).

Ramos DG de S, Scheremeta RGA da C, Oliveira ACS de, Sinkoc AL, Pacheco R de C. 2013. Survey of helminth parasites of cats from the metropolitan area of Cuiabá, Mato Grosso, Brazil. Rev Bras Parasitol Vet. 22:201–206. doi:[10.1590/S1984-29612013000200040](https://doi.org/10.1590/S1984-29612013000200040).

Ranganathan S, Balajee SA, Raja SM. 1997. A survey of dermatophytosis in animals in Madras, India. Mycopathologia. 140(3):137–140.

Ranjbar-Bahadori S, Veshgini A, Shirani D, Eslami A, Mohieddin H, Shemshadi B, Masooleh R. 2011. Epidemiological aspects of canine Dirofilariasis in the north of Iran. Iran J Parasitol. 6(1):73–80.

Raoof P, Garedaghi Y. 2020 Mar 22. Investigation of infection with *Dirofilaria immitis* parasite in stray dogs in Tabriz city of Iran. Livest Sci. 8:38-42.

Ravagnan S CG Carli E, Piseddu E, Da Rold G, Porcellato E, Zanardello C, Carminato A, Vascellari M. 2017. Prevalence and molecular characterization of canine and feline Hemotropic Mycoplasmas (Hemoplasmas) in northern Italy. Parasit Vectors. 10(1):132. doi:[10.1186/s13071-017-2069-9](https://doi.org/10.1186/s13071-017-2069-9).

Regidor-Cerrillo J, Pedraza-Diaz S, Rojo-Montejo S, Vazquez-Moreno E, Arnaiz I, Gomez-Bautista M, Jimenez-Palacios S, Ortega-Mora LM, Collantes-Fernandez E. 2010. *Neospora caninum* infection in stray and farm dogs: seroepidemiological study and oocyst shedding. Vet Parasitol. 174(3–4):332–335. doi:[10.1016/j.vetpar.2010.08.033](https://doi.org/10.1016/j.vetpar.2010.08.033).

Rembiesa C, Richardson D. 2003. Helminth Parasites of the House Cat, *Felis catus*, in Connecticut, U.S.A. Comp Parasitol. 70:115-119.

Ribeiro CM, Matos AC, Azzolini T, Bones ER, Wasnieski EA, Richini-Pereira VB, Lucheis SB, Vidotto O. 2017. Molecular epidemiology of *Anaplasma platys*, *Ehrlichia canis* and *Babesia vogeli* in stray dogs in Paraná, Brazil. Pesq Vet Bras. 37:129–136. doi:[10.1590/S0100-736X2017000200006](https://doi.org/10.1590/S0100-736X2017000200006).

Rodríguez-Ponce E, González JF, Conde de Felipe M, Hernández JN, Raduan Jaber J. 2016. Epidemiological survey of zoonotic helminths in feral cats in Gran Canaria island (Macaronesian archipelago-Spain). Acta Parasitol. 61(3):443–450. doi:[10.1515/ap-2016-0059](https://doi.org/10.1515/ap-2016-0059).

Rojero-Vázquez E, Gordillo-Pérez G, Weber M. 2017. Infection of *Anaplasma phagocytophilum* and *Ehrlichia* spp. in opossums and dogs in Campeche, Mexico: The role of tick infestation. Front Ecol Evol. 5:161. doi:[10.3389/fevo.2017.00161](https://doi.org/10.3389/fevo.2017.00161).

Romano C, Valenti L, Barbara R. 1997. Dermatophytes isolated from asymptomatic stray cats. Mycoses. 40(11–12):471–472. doi:[10.1111/j.1439-0507.1997.tb00187.x](https://doi.org/10.1111/j.1439-0507.1997.tb00187.x).

Romero-Callejas E, Rendón-Franco E, Villanueva-García C, Osorio-Sarabia D, Muñoz-García CI. 2014. Risk of Cutaneous Larva Migrans and other zoonotic parasites infections due to feral cats from a touristic tropical park. Travel Med Infect Dis. 12(6 Pt A):684–686. doi:[10.1016/j.tmaid.2014.10.018](https://doi.org/10.1016/j.tmaid.2014.10.018).

Rondon FCM, Bevilaqua CML, Franke CR, Barros RS, Oliveira FR, Alcântara AC, Diniz AT. 2008. Cross-sectional serological study of canine *Leishmania* infection in Fortaleza, Ceará state, Brazil. Vet Parasitol. 155(1–2):24–31. doi:[10.1016/j.vetpar.2008.04.014](https://doi.org/10.1016/j.vetpar.2008.04.014).

Rosypal AC, Bowman SS, Epps SA, El Behairy AM, Hilali M, Dubey JP. 2013. Serological survey of dogs from Egypt for antibodies to *Leishmania* species. J Parasitol. 99(1):170–171. doi:[10.1645/GE-3242.1](https://doi.org/10.1645/GE-3242.1).

Ruiz J, Giraldo-Echeverri C, López L, Chica J. 2010. Brucella canis seroprevalence in stray dogs from "Centro de Bienestar Animal “La Perla”, Medellín (Colombia), 2008. Rev Colomb de Cienc Pecu. 23:166–172.

Ryan GE. 1976. Gastro-intestinal parasites of feral cats in New South Wales. Aust Vet J. 52(5):224–227. doi:[10.1111/j.1751-0813.1976.tb00072.x](https://doi.org/10.1111/j.1751-0813.1976.tb00072.x).

Sadighian A. 1969. Helminth parasites of stray dogs and jackals in Shahsavar area, caspian region, Iran. J Parasitol. 55(2):372–374.

Saeed I, Kapel C, Saida LA, Willingham L, Nansen P. 2000. Epidemiology of *Echinococcus granulosus* in Arbil province, northern Iraq, 1990-1998. J Helminthol. 74(1):83–88. doi:[10.1017/s0022149x00000111](https://doi.org/10.1017/s0022149x00000111).

Santana CC, Vassallo J, De Freitas LAR, Oliveira GGS, Pontes-De-Carvalho LC, Dos-Santos WLC. 2008. Inflammation and structural changes of splenic lymphoid tissue in Visceral Leishmaniasis: A study on naturally infected dogs. Parasite Immunol. 30(10):515–524. doi:[10.1111/j.1365-3024.2008.01051.x](https://doi.org/10.1111/j.1365-3024.2008.01051.x).

Sardarian K ZAH Maghsood AH, Ghiasian SA. 2015. Prevalence of zoonotic intestinal parasites in household and stray dogs in rural areas of Hamadan, Western Iran. Trop Biomed. 32(2):240–246.

Saridomichelakis MN, Apostolidis K, Chatzis M, Petanides T, Kokkinaki K, Athanasiou L, Kasabalis D, Leontides L. 2018. Are stray dogs confined in animal shelters at increased risk of seropositivity to *Leishmania infantum*? A case control study. Rev Med Vet. 1(169):12-23.

Savani ES, Galati EA, Camargo MC, D’Auria SR, Damaceno JT, Balduino SA. 1999. [Serological survey for American cutaneous leishmaniasis in stray dogs in the S. Paulo State, Brazil]. Rev Saude Publica. 33(6):629–631. doi:[10.1590/s0034-89101999000600017](https://doi.org/10.1590/s0034-89101999000600017).

Scanziani E, Origgi F, Giusti AM, Iacchia G, Vasino A, Pirovano G, Scarpa P, Tagliabue S. 2002. Serological survey of leptospiral infection in kennelled dogs in Italy. J Small Anim Pract. 43(4):154–157. doi:[10.1111/j.1748-5827.2002.tb00048.x](https://doi.org/10.1111/j.1748-5827.2002.tb00048.x).

Seaman RL, Kania SA, Hegarty BC, Legendre AM, Breitschwerdt EB. 2004. Comparison of results for serologic testing and a polymerase chain reaction assay to determine the prevalence of stray dogs in eastern Tennessee seropositive to *Ehrlichia canis*. Am J Vet Res. 65(9):1200–1203. doi:[10.2460/ajvr.2004.65.1200](https://doi.org/10.2460/ajvr.2004.65.1200).

Segura F, Pons I, Miret J, Pla J, Ortuño A, Nogueras M-M. 2014. The role of cats in the eco-epidemiology of spotted fever group diseases. Parasit Vectors. 7:353. doi:[10.1186/1756-3305-7-353](https://doi.org/10.1186/1756-3305-7-353).

Shalaby HA, Abdel-Shafy S, Derbala AA. 2010. The role of dogs in transmission of *Ascaris lumbricoides* for humans. Parasitol Res. 106(5):1021–1026. doi:[10.1007/s00436-010-1755-8](https://doi.org/10.1007/s00436-010-1755-8).

Shamsi S, McSpadden K, Baker S, Jenkins DJ. 2017. Occurrence of tongue worm, *Linguatula cf. serrata* (Pentastomida: Linguatulidae) in wild canids and livestock in south-eastern Australia. Int J Parasitol. 6(3):271–277. doi:[10.1016/j.ijppaw.2017.08.008](https://doi.org/10.1016/j.ijppaw.2017.08.008).

Shapiro AJ, Bosward KL, Heller J, Norris JM. 2015. Seroprevalence of *Coxiella burnetii* in domesticated and feral cats in eastern Australia. Vet Microbiol. 177(1–2):154–161. doi:[10.1016/j.vetmic.2015.02.011](https://doi.org/10.1016/j.vetmic.2015.02.011).

Shapiro AJ, Brown G, Norris JM, Bosward KL, Marriot DJ, Balakrishnan N, Breitschwerdt EB, Malik R. 2017. Vector-borne and zoonotic diseases of dogs in North-west New South Wales and the Northern Territory, Australia. BMC Vet Res. 13(1):238. doi:[10.1186/s12917-017-1169-2](https://doi.org/10.1186/s12917-017-1169-2).

Shapiro AJ, Norris JM, Heller J, Brown G, Malik R, Bosward KL. 2016. Seroprevalence of *Coxiella burnetii* in Australian dogs. Zoonoses Public Health. 63(6):458–466. doi:[10.1111/zph.12250](https://doi.org/10.1111/zph.12250).

Shariatzadeh SA, Spotin A, Gholami S, Fallah E, Hazratian T, Mahami-Oskouei M, Montazeri F, Moslemzadeh HR, Shahbazi A. 2015. The first morphometric and phylogenetic perspective on molecular epidemiology of *Echinococcus granulosus* sensu lato in stray dogs in a hyperendemic Middle East focus, northwestern Iran. Parasit Vectors. 8:409. doi:[10.1186/s13071-015-1025-9](https://doi.org/10.1186/s13071-015-1025-9).

Sharif M, Nasrolahei M, Ziapour SP, Gholami S, Ziaei H, Daryani A, Khalilian A. 2007. *Toxocara cati* infections in stray cats in northern Iran. J Helminthol. 81(1):63–66. doi:[10.1017/S0022149X07214117](https://doi.org/10.1017/S0022149X07214117).

Sharma R, Kimmitt T, Tiwari K, Chikweto A, Thomas D, Lanza Perea M, Bhaiyat MI. 2015. Serological evidence of antibodies to *Neospora caninum* in stray and owned Grenadian dogs. Trop Biomed. 32(2):286–290.

el-Shehabi FS, Kamhawi SA, Schantz PM, Craig PS, Abdel-Hafez SK. 2000. Diagnosis of canine echinococcosis: comparison of coproantigen detection with necropsy in stray dogs and red foxes from northern Jordan. Parasite. 7(2):83–90. doi:[10.1051/parasite/2000072083](https://doi.org/10.1051/parasite/2000072083).

Shin J-C, Reyes AWB, Kim S-H, Kim S, Park H-J, Seo K-W, Song K-H. 2015. Molecular Detection of *Giardia intestinalis* from Stray Dogs in Animal Shelters of Gyeongsangbuk-do (Province) and Daejeon, Korea. Korean J Parasitol. 53(4):477–481. doi:[10.3347/kjp.2015.53.4.477](https://doi.org/10.3347/kjp.2015.53.4.477).

Shin S-S, Oh D-S, Ahn K-S, Cho S-H, Lee W-J, Na B-K, Sohn W-M. 2015. Zoonotic intestinal trematodes in stray cats (*Felis catus*) from riverside areas of the Republic of Korea. Korean J Parasitol. 53(2):209–213. doi:[10.3347/kjp.2015.53.2.209](https://doi.org/10.3347/kjp.2015.53.2.209).

Silaghi C, Knaus M, Rapti D, Kusi I, Shukullari E, Hamel D, Pfister K, Rehbein S. 2014. Survey of *Toxoplasma gondii* and *Neospora caninum*, haemotropic mycoplasmas and other arthropod-borne pathogens in cats from Albania. Parasit Vectors. 7(1):62. doi:[10.1186/1756-3305-7-62](https://doi.org/10.1186/1756-3305-7-62).

da Silva JR, Maciel BM, de Santana Souza Santos LKN, Carvalho FS, de Santana Rocha D, Lopes CWG, Albuquerque GR. 2017. Isolation and genotyping of *Toxoplasma gondii* in Brazilian dogs. Korean J Parasitol. 55(3):239–246. doi:[10.3347/kjp.2017.55.3.239](https://doi.org/10.3347/kjp.2017.55.3.239).

Silva RC da, Souza LC de, Langoni H, Tanaka EM, Lima VY de, Silva AV da. 2010. Risk factors and presence of antibodies to *Toxoplasma gondii* in dogs from the coast of São Paulo State, Brazil. Pesq Vet Bras. 30:161–166. doi:[10.1590/S0100-736X2010000200011](https://doi.org/10.1590/S0100-736X2010000200011).

Silva RC, Richini-Pereira VB, Kikuti M, Marson PM, Langoni H. 2017. Detection of *Leishmania (L.) infantum* in stray dogs by molecular techniques with sensitive species-specific primers. Vet Q. 37(1):23–30. doi:[10.1080/01652176.2016.1252073](https://doi.org/10.1080/01652176.2016.1252073).

Simking P, Wongnakphet S, Stich RW, Jittapalapong S. 2010. Detection of *Babesia vogeli* in stray cats of metropolitan Bangkok, Thailand. Vet Parasitol. 173(1–2):70–75. doi:[10.1016/j.vetpar.2010.06.025](https://doi.org/10.1016/j.vetpar.2010.06.025).

Simsek S, Ciftci A. 2016. Serological and Molecular Detection of *Dirofilaria* Species in Stray Dogs and Investigation of *Wolbachia* DNA by PCR in Turkey. J Arthropod Borne Dis. 10(4):445–453.

Simsek S, Ozkanlar Y, Balkaya I, Aktas MS. 2011. Microscopic, serologic and molecular surveys on *Dirofilaria immitis* in stray dogs, Turkey. Vet Parasitol. 183(1–2):109–113. doi:[10.1016/j.vetpar.2011.06.012](https://doi.org/10.1016/j.vetpar.2011.06.012).

Simsek S, Utuk AE, Koroglu E, Rishniw M. 2008. Serological and molecular studies on *Dirofilaria immitis* in dogs from Turkey. J Helminthol. 82(2):181–186. doi:[10.1017/S0022149X0896079X](https://doi.org/10.1017/S0022149X0896079X).

Singh B, Sharma R, Sharma J, Singh Gill J. 2014. Molecular detection of *E. granulosus* sheep strain (G1) infections in naturally infected dogs in Punjab (India). Helminthologia. 51(4):269–272. doi:[10.2478/s11687-014-0240-9](https://doi.org/10.2478/s11687-014-0240-9).

Smout FA, Skerratt LF, Butler JRA, Johnson CN, Congdon BC, Thompson RCA. 2017. The hookworm *Ancylostoma ceylanicum*: An emerging public health risk in Australian tropical rainforests and Indigenous communities. One Health. 3:66–69. doi:[10.1016/j.onehlt.2017.04.002](https://doi.org/10.1016/j.onehlt.2017.04.002).

Smout FA, Skerratt LF, Johnson CN, Butler JRA, Congdon BC. 2018. Zoonotic Helminth Diseases in Dogs and Dingoes Utilising Shared Resources in an Australian Aboriginal Community. Trop Med Infect Dis. 3(4):110. doi:[10.3390/tropicalmed3040110](https://doi.org/10.3390/tropicalmed3040110).

Spada E PD Canzi I, Baggiani L, Perego R, Vitale F, Migliazzo A. 2016. Prevalence of *Leishmania infantum* and co-infections in stray cats in northern Italy. Comp Immunol Microbiol Infect Dis. 45:53–58. doi:[10.1016/j.cimid.2016.03.001](https://doi.org/10.1016/j.cimid.2016.03.001).

Spada E, Proverbio D, Galluzzo P, Della Pepa A, Perego R, Bagnagatti De Giorgi G, Ferro E. 2014. Molecular study on selected vector-borne infections in urban stray colony cats in northern Italy. J Feline Med Surg. 16(8):684–688. doi:[10.1177/1098612X13514422](https://doi.org/10.1177/1098612X13514422).

Spolidorio MG, Minervino AHH, Valadas SYOB, Soares HS, Neves KAL, Labruna MB, Ribeiro MFB, Gennari SM. 2013. Serosurvey for tick-borne diseases in dogs from the Eastern Amazon, Brazil. Rev Bras Parasitol Vet. 22:214–219. doi:[10.1590/S1984-29612013005000023](https://doi.org/10.1590/S1984-29612013005000023).

Stanek JF, Stich RW, Dubey JP, Reed SM, Njoku CJ, Lindsay DS, Schmall LM, Johnson GK, LaFave BM, Saville WJA. 2003. Epidemiology of *Sarcocystis neurona* infections in domestic cats (*Felis domesticus*) and its association with equine protozoal myeloencephalitis (EPM) case farms and feral cats from a mobile spay and neuter clinic. Vet Parasitol. 117(4):239–249. doi:[10.1016/j.vetpar.2003.09.002](https://doi.org/10.1016/j.vetpar.2003.09.002).

Stojanovic V, Foley P. 2011. Infectious disease prevalence in a feral cat population on Prince Edward Island, Canada. Can Vet J. 52(9):979–982.

Su S, Qi W, Zhou P, Xiao C, Yan Z, Cui J, Jia K, Zhang G, Gray GC, Liao M, et al. 2014. First evidence of H10N8 Avian Influenza Virus infections among feral dogs in live poultry markets in Guangdong Province, China. Clin Infect Dis. 59(5):748–750. doi:[10.1093/cid/ciu345](https://doi.org/10.1093/cid/ciu345).

Su S, Zhou P, Fu X, Wang L, Hong M, Lu G, Sun L, Qi W, Ning Z, Jia K, et al. 2014. Virological and epidemiological evidence of Avian Influenza Virus infections among feral dogs in live poultry markets, China: a threat to human health? Clin Infect Dis. 58(11):1644–1646. doi:[10.1093/cid/ciu154](https://doi.org/10.1093/cid/ciu154).

Suepaul SM, Carrington CV, Campbell M, Borde G, Adesiyun AA. 2014. Seroepidemiology of Leptospirosis in dogs and rats in Trinidad. Trop Biomed. 31(4):853–861.

Suepaul SM, Carrington CVF, Campbell M, Borde G, Adesiyun AA. 2010. Serovars of *Leptospira* isolated from dogs and rodents. Epidemiol Infect. 138(7):1059–1070. doi:[10.1017/S0950268809990902](https://doi.org/10.1017/S0950268809990902).

Sultanov A, Abdybekova A, Abdibaeva A, Shapiyeva Z, Yeshmuratov T, Torgerson PR. 2014. Epidemiology of fishborne trematodiasis in Kazakhstan. Acta Trop. 138:60–66. doi:[10.1016/j.actatropica.2014.04.030](https://doi.org/10.1016/j.actatropica.2014.04.030).

Switzer AD, McMillan-Cole AC, Kasten RW, Stuckey MJ, Kass PH, Chomel BB. 2013. *Bartonella* and *Toxoplasma* infections in stray cats from Iraq. Am J Trop Med Hyg. 89(6):1219–1224. doi:[10.4269/ajtmh.13-0353](https://doi.org/10.4269/ajtmh.13-0353).

Szwabe K, Blaszkowska J. 2017. Stray dogs and cats as potential sources of soil contamination with zoonotic parasites. Ann Agric Environ Med. 24(1):39–43. doi:[10.5604/12321966.1234003](https://doi.org/10.5604/12321966.1234003).

Talebkhan Garoussi M, Mehravaran M, Abdollahpour G, Khoshnegah J. 2015. Seroprevalence of leptospiral infection in feline population in urban and dairy cattle herds in Mashhad, Iran. Vet Res Forum. 6(4):301–304.

Talvik H, Moks E, Mägi E, Järvis T, Miller I. 2006. Distribution of *Toxocara* infection in the environment and in definitive and paratenic hosts in Estonia. Acta Vet Hung. 54(3):399–406. doi:[10.1556/AVet.54.2006.3.10](https://doi.org/10.1556/AVet.54.2006.3.10).

Tarish JH, Al-Saqur IM, Al-Abbassy SN, Kadhim FS. 1986. The prevalence of parasitic helminths in stray dogs in the Baghdad area, Iraq. Ann Trop Med Parasitol. 80(3):329–331. doi:[10.1080/00034983.1986.11812024](https://doi.org/10.1080/00034983.1986.11812024).

Terao M, Akter S, Yasin MG, Nakao R, Kato H, Alam MZ, Katakura K. 2015. Molecular detection and genetic diversity of *Babesia gibsoni* in dogs in Bangladesh. Infect Genet Evol. 31:53–60. doi:[10.1016/j.meegid.2015.01.011](https://doi.org/10.1016/j.meegid.2015.01.011).

Thomas JE, Staubus L, Goolsby JL, Reichard MV. 2016. Ectoparasites of free-roaming domestic cats in the central United States. Vet Parasitol. 228:17–22. doi:[10.1016/j.vetpar.2016.07.034](https://doi.org/10.1016/j.vetpar.2016.07.034).

Tian Y-M, Cao J-F, Zhou D-H, Zou F-C, Miao Q, Liu Z-L, Li B-F, Lv R-Q, Du X-P, Zhu X-Q. 2014. Seroprevalence and risk factors of *Chlamydia* infection in dogs in Southwestern China. Acta Tropica. 130:67–70. doi:[10.1016/j.actatropica.2013.09.027](https://doi.org/10.1016/j.actatropica.2013.09.027).

Tiao N, Darrington C, Molla B, Saville WJA, Tilahun G, Kwok OCH, Gebreyes WA, Lappin MR, Jones JL, Dubey JP. 2013. An investigation into the seroprevalence of *Toxoplasma gondii*, *Bartonella* spp., Feline Immunodeficiency Virus (FIV), and Feline Leukaemia Virus (FeLV) in cats in Addis Ababa, Ethiopia. Epidemiol Infect. 141(5):1029–1033. doi:[10.1017/S0950268812001707](https://doi.org/10.1017/S0950268812001707).

Tiawsirisup S, Thanapaisarnkit T, Varatorn E, Apichonpongsa T, Bumpenkiattikun N, Rattanapuchpong S, Chungpiwat S, Sanprasert V, Nuchprayoon S. 2010. Canine Heartworm (*Dirofilaria immitis*) infection and immunoglobulin G antibodies against *Wolbachia* (Rickettsiales: Rickettsiaceae) in stray dogs in Bangkok, Thailand. Thai J Vet Med. 40(2): 165-170.

Torkan S, Ghandehari-Alavijeh MR, Khamesipour F. 2017. Survey of the prevalence of *Toxocara cati* in stray cats in Isfahan city, Iran by PCR method. Trop Biomed. 34(3):550–555.

Trasviña-Muñoz E, Valencia G, Centeno P, Cueto Gonzalez SA, Monge F, Tinoco-Gracia L, Núñez-Castro K, Pérez-Ortiz P, Medina-Basulto G, Tamayo A, et al. 2017. Prevalence and distribution of intestinal parasites in stray dogs in the northwest area of Mexico. Arch Vet Med. 49:105–111. doi:[10.4067/S0719-81322017000200105](https://doi.org/10.4067/S0719-81322017000200105).

Tsai H-J, Huang H-C, Lin C-M, Lien Y-Y, Chou C-H. 2007. Salmonellae and campylobacters in household and stray dogs in northern Taiwan. Vet Res Commun. 31(8):931–939. doi:[10.1007/s11259-007-0009-4](https://doi.org/10.1007/s11259-007-0009-4).

Tseng YC PSY Ho GD, Chen T TW, Huang BF, Cheng PC, Chen JL. 2014. Prevalence and genotype of *Giardia duodenalis* from faecal samples of stray dogs in Hualien city of eastern Taiwan. Trop Biomed. 31(2):305–311.

Tsukada R, Osaka Y, Takano T, Sasaki M, Inose M, Ikadai H. 2016. Serological survey of *Encephalitozoon cuniculi* infection in cats in Japan. J Vet Med Sci. 78(10):1615–1617. doi:[10.1292/jvms.15-0545](https://doi.org/10.1292/jvms.15-0545).

Tuemmers C CC Lüders C, Rojas C, Serri M, Espinoza R. 2013. [Prevalence of Leptospirosis in vague dogs captured in Temuco City, 2011]. Rev Chilena Infectol. 30(3):252–257. doi:[10.4067/s0716-10182013000300003](https://doi.org/10.4067/s0716-10182013000300003).

Tun S, Ithoi I, Mahmud R, Samsudin NI, Kek Heng C, Ling LY. 2015. Detection of helminth eggs and identification of hookworm species in stray cats, dogs and soil from Klang Valley, Malaysia. PLoS One. 10(12):e0142231. doi:[10.1371/journal.pone.0142231](https://doi.org/10.1371/journal.pone.0142231).

Umar Y. 2009. Intestinal Helminthoses in Dogs in Kaduna Metropolis, Kaduna State, Nigeria. Iran J Parasitol. 4(1):34–39.

Valadas S, Minervino AHH, Lima VMF, Soares RM, Ortolani EL, Gennari SM. 2010. Occurrence of antibodies anti-*Neospora caninum*, anti-*Toxoplasma gondii*, and anti-*Leishmania chagasi* in serum of dogs from Pará State, Amazon, Brazil. Parasitol Res. 107(2):453–457. doi:[10.1007/s00436-010-1890-2](https://doi.org/10.1007/s00436-010-1890-2).

Vanparijs O, Hermans L, van der Flaes L. 1991. Helminth and protozoan parasites in dogs and cats in Belgium. Vet Parasitol. 38(1):67–73. doi:[10.1016/0304-4017(91)90010-s](https://doi.org/10.1016/0304-4017(91)90010-s).

Vanwormer E, Conrad PA, Miller MA, Melli AC, Carpenter TE, Mazet JAK. 2013. *Toxoplasma gondii*, source to sea: higher contribution of domestic felids to terrestrial parasite loading despite lower infection prevalence. Ecohealth. 10(3):277–289. doi:[10.1007/s10393-013-0859-x](https://doi.org/10.1007/s10393-013-0859-x).

Vascellari M, Ravagnan S, Carminato A, Cazzin S, Carli E, Da Rold G, Lucchese L, Natale A, Otranto D, Capelli G. 2016. Exposure to vector-borne pathogens in candidate blood donor and free-roaming dogs of northeast Italy. Parasit Vectors. 9(1):369. doi:[10.1186/s13071-016-1639-6](https://doi.org/10.1186/s13071-016-1639-6).

Waap H, Gomes J, Nunes T. 2014. Parasite communities in stray cat populations from Lisbon, Portugal. J Helminthol. 88(4):389–395. doi:[10.1017/S0022149X1300031X](https://doi.org/10.1017/S0022149X1300031X).

Wang L. 1997. Canine filarial infections in north Taiwan. Acta Trop. 68(1):115–120. doi:[10.1016/s0001-706x(97)00081-8](https://doi.org/10.1016/s0001-706x(97)00081-8).

Wang L, Zheng Y, Fu C, Huang S, Hong M, Yan Z, Jia K, Zhou P, Li S. 2016. Seroprevalence of Hepatitis E virus infection among dogs in several developed cities in the Guangdong province of China. J Med Virol. 88(8):1404–1407. doi:[10.1002/jmv.24468](https://doi.org/10.1002/jmv.24468).

Wang LC. 1998. Comparison of a whole-blood agglutination test and an ELISA for the detection of the antigens of *Dirofilaria immitis* in dogs. Ann Trop Med Parasitol. 92(1):73–77. doi:[10.1080/00034989860193](https://doi.org/10.1080/00034989860193).

Wang W, Cuttell L, Bielefeldt-Ohmann H, Inpankaew T, Owen H, Traub RJ. 2013. Diversity of *Blastocystis* subtypes in dogs in different geographical settings. Parasit Vectors. 6(1):215. doi:[10.1186/1756-3305-6-215](https://doi.org/10.1186/1756-3305-6-215).

Winkler IG, Löchelt M, Flower RL. 1999. Epidemiology of Feline Foamy Virus and Feline Immunodeficiency Virus infections in domestic and feral cats: a seroepidemiological study. J Clin Microbiol. 37(9):2848–2851. doi:[10.1128/JCM.37.9.2848-2851.1999](https://doi.org/10.1128/JCM.37.9.2848-2851.1999).

Wu C, Fan P. 2003. Prevalence of canine dirofilariasis in Taiwan. J Helminthol. 77(1):83–88. doi:[10.1079/joh2002150](https://doi.org/10.1079/joh2002150).

Yaglom HD, Nicholson WL, Casal M, Nieto NC, Adams L. 2018. Serologic assessment for exposure to Spotted Fever Group Rickettsiae in dogs in the Arizona-Sonora border region. Zoonoses Public Health. 65(8):984–992. doi:[10.1111/zph.12517](https://doi.org/10.1111/zph.12517).

Yagoob G. 2012. Seroprevalence of *Neospora caninum* in stray dogs of Tabriz, Iran. J Anim Vet Adv. 11(6):723-726.

Yakhchali M, Hajipour N, Malekzadeh-Viayeh R, Esmaeilnejad B, Nemati-Haravani T, Fathollahzadeh M, Jafari R. 2017. Gastrointestinal helminths and ectoparasites in the stray cats (Felidae: *Felis catus*) of Ahar Municipality, Northwestern Iran. Iran J Parasitol. 12(2):298–304.

Yakhchali M MA Javadi S. 2010. Prevalence of antibodies to *Neospora caninum* in stray dogs of Urmia, Iran. Parasitol Res. 106(6):1455–1458. doi:[10.1007/s00436-010-1824-z](https://doi.org/10.1007/s00436-010-1824-z).

Yan C, Fu L-L, Yue C-L, Tang R-X, Liu Y-S, Lv L, Shi N, Zeng P, Zhang P, Wang D-H, et al. 2012. Stray dogs as indicators of *Toxoplasma gondii* distributed in the environment: the first report across an urban-rural gradient in China. Parasit Vectors. 5(1):5. doi:[10.1186/1756-3305-5-5](https://doi.org/10.1186/1756-3305-5-5).

Yildirim A IA Ica A, Atalay O, Duzlu O. 2007. Prevalence and epidemiological aspects of *Dirofilaria immitis* in dogs from Kayseri Province, Turkey. Res Vet Sci. 82(3):358–363. doi:[10.1016/j.rvsc.2006.08.006](https://doi.org/10.1016/j.rvsc.2006.08.006).

Yoak AJ, Reece JF, Gehrt SD, Hamilton IM. 2014. Disease control through fertility control: Secondary benefits of animal birth control in Indian street dogs. Prev Vet Med. 113(1):152–156. doi:[10.1016/j.prevetmed.2013.09.005](https://doi.org/10.1016/j.prevetmed.2013.09.005).

Yu D-H, Kim H-W, Desai AR, Han I-A, Li Y-H, Lee M-J, Kim I-S, Chae J-S, Park J. 2007. Molecular detection of feline hemoplasmas in feral cats in Korea. J Vet Med Sci. 69(12):1299–1301. doi:[10.1292/jvms.69.1299](https://doi.org/10.1292/jvms.69.1299).

Zaidi S, Bouam A, Bessas A, Hezil D, Ghaoui H, Ait-Oudhia K, Drancourt M, Bitam I. 2018. Urinary shedding of pathogenic *Leptospira* in stray dogs and cats, Algiers: A prospective study. PLoS One. 13(5):e0197068. doi:[10.1371/journal.pone.0197068](https://doi.org/10.1371/journal.pone.0197068).

Zhang H, Zhou D-H, Chen Y-Z, Lin R-Q, Yuan Z-G, Song H-Q, Li S-J, Zhu X-Q. 2010. Antibodies to *Toxoplasma gondii* in stray and household dogs in Guangzhou, China. J Parasitol. 96(3):671–672. doi:[10.1645/GE-2352.1](https://doi.org/10.1645/GE-2352.1).

Zhang X-X, Cai Y-N, Wang C-F, Jiang J, Xu Y-T, Yang G-L, Zhao Q. 2015. Seroprevalence and risk factors of *Toxoplasma gondii* infection in stray dogs in northern China. Parasitol Res. 114(12):4725–4729. doi:[10.1007/s00436-015-4746-y](https://doi.org/10.1007/s00436-015-4746-y).

Zhang Y, Zhong Z, Deng L, Wang M, Li W, Gong C, Fu H, Cao S, Shi X, Wu K, et al. 2017. Detection and multilocus genotyping of *Giardia duodenalis* in dogs in Sichuan province, China. Parasite. 24:31. doi:[10.1051/parasite/2017032](https://doi.org/10.1051/parasite/2017032).

Zhang Y-B, Chen J-D, Xie J-X, Zhu W-J, Wei C-Y, Tan L-K, Cao N, Chen Y, Zhang M-Z, Zhang G-H, et al. 2013. Serologic reports of H3N2 Canine Influenza Virus infection in dogs in northeast China. J Vet Med Sci. 75(8):1061–1062. doi:[10.1292/jvms.13-0022](https://doi.org/10.1292/jvms.13-0022).

Zheng G, Hu W, Liu Y, Luo Q, Tan L, Li G. 2015. Occurrence and molecular identification of *Giardia duodenalis* from stray cats in Guangzhou, southern China. Korean J Parasitol. 53(1):119–124. doi:[10.3347/kjp.2015.53.1.119](https://doi.org/10.3347/kjp.2015.53.1.119).

Zhou H, He S, Sun L, He H, Ji F, Sun Y, Jia K, Ning Z, Wang H, Yuan L, et al. 2015. Serological evidence of Avian Influenza Virus and Canine Influenza Virus infections among stray cats in live poultry markets, China. Vet Microbiol. 175(2–4):369–373. doi:[10.1016/j.vetmic.2014.12.018](https://doi.org/10.1016/j.vetmic.2014.12.018).

Zibaei M SB Sadjjadi SM. 2007. Prevalence of *Toxocara cati* and other intestinal helminths in stray cats in Shiraz, Iran. Trop Biomed. 24(2):39–43.

Σπανακος) GS (γ, Παπαδογιαννακης) EP (ε, Κοντος) VK (β, Μενουνος) PM (π, Βελονακης) EV (ε, Κουτης) CK (χ, Βακαλης) NCV (ν. 2011. Molecular screening for *Blastocystìs* sp. in canine faecal samples in Greece. J Hell Vet Medical Soc. 62(3):216–220. doi:[10.12681/jhvms.14852](https://doi.org/10.12681/jhvms.14852).
