## Supplementary Figure 4 for "Parasite and pathogen prevalence in our closest animal companions is determined by accessibility of sanitation services"

**Supplementary Figure 4: Socioeconomic and Environmental Variable Database information**

**GDP, Lindgren 2019**

Per capita gross domestic product (GDP) for all countries in the free-roaming companion animal infectious agent database was obtained from the World Bank database (Lindgren 2019). This database provides per capita gross domestic product values expressed as current international dollars (purchasing power of the U.S. dollar at a given time (The World Bank 2022)) and converted by purchasing power parity (PPP). This value is obtained using the total gross value by all producers in each country (minus subsidies) and divided by total population of the country (all residents regardless of citizenship). This indicator was attributed to the database of free-roaming companion animal infectious agent prevalence by matching the country in which each study was conducted, using the year of publication to match the date.

**NBI, Convention on Biological Diversity 2011**

The National Biodiversity Index (Convention on Biological Diversity 2011) is based on an index from 0.000 to 1.000 reflecting minimum (i.e., Greenland) and maximum (i.e. Indonesia) country richness and endemism across four terrestrial vertebrate classes and vascular plants.

For regions where free-roaming companion animal infectious agents were sampled that did not have a national biodiversity index, the index for a country within the same geographic world region and of a similar land mass was used (Table 1).

**Table 1:** National biodiversity index (NBI) replacements from countries of similar land mass within the same world region

| **Country/ Land Mass** | **Area (km^2^)** | **Country NBI Used** | **Country NBI Used Area (km^2)** | **NBI** | **Region** |
| --- | --- | --- | --- | --- | --- |
| Puerto Rico | 9104 | Dominican Republic | 15,545 | 0.661 | Caribbean |
| Turks and Caicos | 948 | Bahamas | 1458 | 0.443 | Caribbean |

**Absolute latitude**

Absolute latitude was obtained from the approximate center-point of each country where each study was conducted using Google Earth version 7.3.2.5776.

**Gini index, World Bank 2020a**

Most Gini index values was obtained using data from the World Bank (2020a). The Gini index of income inequality (World Bank 2020a) is based on a scale from 0 (perfect equality) to 1 (perfect inequality) and reflects the deviation from equal distribution of income within individuals or households in an economy.

There were thirteen countries for which The World Bank (2020a) did not have data, and gini coefficients for these countries were thus obtained separately (see Table X for countries, obtained gini index values, and citation).

**Table 2:** List of gini coefficients and sources for countries with gini coefficients not available in The World Bank database

| **Country** | **Gini** | **Source** |
| --- | --- | --- |
| Grenada | 37.0 | Government of Grenada 2008 |
| Kuwait | 47.5 | Al-Qudsi 1981 |
| Libya | 30.7 | Hassine 2015 |
| Puerto Rico | 46.6 | Data USA 2019 |
| Qatar | 29.3 | Ministry of Development Planning and Statistics 2015 |
| Saudi Arabia | 45.9 | Kittaneh 2019 |
| New Zealand | 34.2 | Ministry of Social Development 2016 |

**Sanitation**

Sanitation access in the free-roaming companion animal infectious agent database is expressed as a percentage of the population with access to basic and/or safely managed sanitation services. Basic sanitation services are defined as sanitation facilities that are not shared between households, and safely managed sanitation services include flush/pour flush piped sewage systems, septic tanks, pit latrines, and composting toilets (The World Bank 2020b). This data is from a World Health Organization (WHO)/United Nations Children’s Fund (UNICEF) Joint Monitoring Program (JMP) for Water Supply, Sanitation, and Hygiene (United Nation's Childrens Fund (UNICEF) and World Health Organization 2019). Most of this data was obtained from the World Bank (2020b), but sanitation data for La Reunion was not included in the World Bank database, and was instead obtained directly from the World Health Organization 2019 JMP wash data report (United Nation's Childrens Fund (UNICEF) and World Health Organization 2019).

**Human Population**

Human population data for most countries in the free-roaming companion animal infectious agent database was obtained from The World Bank (2020c) and is based on the total resident population of the country, regardless of citizenship. Population estimates for Reunion and Taiwan were not available in the World Bank population database (The World Bank 2020c) and were thus obtained from separately. Population estimates from the United Nations Department of Economic and Social Affairs Population Division (2019) were used for Reunion, and population estimates from the Republic of China (2021) were used for Taiwan.

United Nations, Department of Economic and Social Affairs, Population Division. 2019. World Population Prospects 2019, Online Edition. Rev. 1.
