## Supplementary Figure 2 for "Parasite and pathogen prevalence in our closest animal companions is determined by accessibility of sanitation services": SuppFig2_LeaveOneOut_8MAR2022.html

Leave One Out


### Leave One Out

#### Purpose

The purpose of this script is to run through a leave-one-out bias check for two analyses investigating global associations between parasite prevalence of free-roaming dogs and socioeconomic, geographical, and ecological variables. In a leave-one-out bias check, individual studies are removed one at a time and differences in the overall effect size and significance of effect (i.e., p value) between the full data set and the data set with one study removed, are recorded. This is done to determine if the effect observed is heavily influenced by any one particular study.

#### Setup

*Set root directory*

*Load the libraries*

```
library(lme4)
```

```
## Loading required package: Matrix
```

```
library(ggplot2)
library(dplyr)
```

```
## 
## Attaching package: 'dplyr'
```

```
## The following objects are masked from 'package:stats':
## 
##     filter, lag
```

```
## The following objects are masked from 'package:base':
## 
##     intersect, setdiff, setequal, union
```

*Load the databases*

```
feral <- read.csv(file="feral_afterdb_04OCT2021.csv")
country <- read.csv(file="country_afterdb_27SEP2021.csv")
survey <- read.csv(file="survey_total_27SEP2021.csv")
zoonotic <- read.csv(file="zoonotic_27SEP2021.csv")
```

```
#View(feral)
#Need to remove added NBI for the following countries:
#Cayman Islands, New Caledonia, Cape Verde, Reunion, Serbia, Saint Kitts, Taiwan

feral[which(feral$country=="GRAND CAYMAN ISLAND"),which(colnames(feral)=="nbii" )]<-NA
feral[which(feral$country=="NEW CALEDONIA"),which(colnames(feral)=="nbii" )]<-NA
feral[which(feral$country=="CAPE VERDE"),which(colnames(feral)=="nbii" )]<-NA
feral[which(feral$country=="REUNION"),which(colnames(feral)=="nbii" )]<-NA
feral[which(feral$country=="SERBIA"),which(colnames(feral)=="nbii" )]<-NA
feral[which(feral$country=="ST KITTS"),which(colnames(feral)=="nbii" )]<-NA
feral[which(feral$country=="TAIWAN"),which(colnames(feral)=="nbii" )]<-NA
feral[which(feral$country=="HONG KONG"),which(colnames(feral)=="nbii" )]<-NA
```

```
nrow(feral) #1539
```

```
## [1] 1539
```

```
length(unique(feral$study)) #470
```

```
## [1] 470
```

**Housekeeping**

```
#Make data frame with variables needed for this analysis
feral<-data.frame(pathogen=feral$pathogen, year=feral$year, N=feral$N, prev=feral$prev, country=feral$country, gdp=feral$gdp, gini=feral$gini, nbii=feral$nbii, zoonotic=feral$zoonotic, uniq=feral$uniq, pathogen=feral$pathogen, study=feral$study, ablat=feral$ablat, sanitation=feral$sanitation, feral=feral$feral, island=feral$island, method=feral$method, population=feral$population, land_sq_km=feral$land_sq_km, prevalence=feral$prevalence)

#Make column for transformed gdp values
feral$gdp_10000<-feral$gdp/10000

#Remove rows that have NAs for any value
feral<-feral[complete.cases(feral), ]

#Make transformed yi in feral dataset for later graphical visualization of meta-regression and trends
#feral <- as.data.frame(escalc(measure="PLO", xi=prev, ni=N, data=feral))
#zoonotic <- as.data.frame(escalc(measure="PLO", xi=prev, ni=N, data=zoonotic))

################


#Need add these to databasing script
```

##### Leave one out bias check for first analysis: effect of socioeconomic and environmental gradients on parasite prevalence in free-roaming companion animals

For this leave-one-out analysis, only changes in effect for sanitation will be observed. This is because this was the most important variable, according to the AIC values. Additionally, the values for sanitation included in this analyses are not evenly distributed, such that there are few data points in countries with very low sanitation access.

Leave it out will remove one study at a time, and reports the results of the glmer model without each study

```
# function to run model
run_model <- function(feral) {
  feral$san_10<-feral$sanitation/10
  res_san<-glmer(prevalence ~ san_10 + (1 | pathogen) + (1 | method) + (1 | study) + (1 | study:uniq), data=feral, family=binomial, weights=N, na.action=na.fail, control = glmerControl(optimizer = "optimx", calc.derivs = FALSE, optCtrl = list(method = "nlminb", starttests = FALSE, kkt = FALSE)))
  return(res_san)
}

# function to get stats of interest 
get_stats <- function(res, studyID) {
  Vcov <- vcov(res, useScale = FALSE)
  betas<-fixef(res)
  se <- sqrt(diag(Vcov))
  zval <- betas / se
  pval <- 2 * pnorm(abs(zval), lower.tail = FALSE)
    
  # store these in a data frame
  df<-data.frame(row.names=NULL,"studyID removed"=studyID, 
                 "beta"=betas[2], "se"=se[2], "zval"=zval[2], "pval"=pval[2])
  return(df)
}

# for exploration here's a subset of the dataset 
#subFeral <- feral[1:30,]

# function to run model and gather stats of interest under leave-one-out (looping through removing a study at a time)
# take as input `feral` 
leaveOneOut <- function(feral) {
  # initialize empty data frame
  df<-data.frame("studyID removed"=numeric(), 
                 "beta"=numeric(), 
                 "se"=numeric(), 
                 "zval"=numeric(), 
                 "pval"=numeric())
  
  # get unique study IDs
  studies <- unique(feral$study)
  
  # loop over unique studies, removing one at a time
  for (val in 1:length(studies)) {
    studyID <- studies[val]
    # remove study of interest
    feralMinusOne <- feral[feral$study!=studyID,]
    # run model on dataset minus study of interest
    res <- run_model(feralMinusOne)
    df.dat <- get_stats(res=res,studyID=studyID)
    df <- rbind(df, df.dat)
  }
  return(df)
}
```

Run the model

```
df_test <- leaveOneOut(feral)

#Add Effect Column
df_test$odds_ratio<-exp(df_test$beta)

head(df_test)
```

ABCDEFGHIJ0123456789

|  | studyID.removed  <int> | beta  <dbl> | se  <dbl> | zval  <dbl> | pval  <dbl> | odds\_ratio  <dbl> |
| --- | --- | --- | --- | --- | --- | --- |
| 1 | 227 | -0.1143875 | 0.03921855 | -2.916669 | 0.003537914 | 0.8919123 |
| 2 | 102 | -0.1136463 | 0.03908445 | -2.907710 | 0.003640853 | 0.8925736 |
| 3 | 310 | -0.1131706 | 0.03924718 | -2.883535 | 0.003932387 | 0.8929983 |
| 4 | 463 | -0.1128226 | 0.03911013 | -2.884741 | 0.003917364 | 0.8933091 |
| 5 | 472 | -0.1115393 | 0.03919933 | -2.845438 | 0.004435040 | 0.8944563 |
| 6 | 106 | -0.1139108 | 0.03930360 | -2.898227 | 0.003752783 | 0.8923376 |

6 rows

```
max(df_test$studyID.removed)
```

```
## [1] 472
```

```
df_test$studyID.removed<-as.numeric(df_test$studyID.removed)
```

**Model 1 (san\_10 only) p value LeaveOneOut plot**

```
pval <- ggplot(df_test, aes(x=studyID.removed, y=pval)) + 
  geom_point() + # plot data
  geom_line(aes(y=0.05), color="red", linetype="dotted") + # add red line for pval of interest
  labs(x="Study Removed", y="P-value", title="Leave-one-out cross-validation to test for publication bias") + # add labels
  geom_text(
    data=df_test %>% filter(pval>0.05), #label studies for p-val > 0.01
    aes(label=studyID.removed),
    nudge_x = 15, nudge_y = 0) # label data points above pval line of interest

pval + theme_classic() + 
  scale_x_discrete(limits=c(1,99)) + # discretize x axis
  scale_y_continuous()
```

```
ggsave("../feral_analysis_output/LeaveOneOut_pvalue.png")
```

**Model 1 (san\_10 only) Effect Size LeaveOneOut plot**

```
effect <- ggplot(df_test, aes(x=studyID.removed, y=odds_ratio)) + 
  geom_point() + # plot data
  #geom_line(aes(y=0.01), color="red", linetype="dotted") + # add red line for pval of interest
  labs(x="Study Removed", y="Prevalence", title="Leave-one-out cross-validation to test for publication bias") + # add labels
  geom_text(
    data=df_test %>% filter(pval>0.05), #label studies for p-val > 0.01
    aes(label=studyID.removed),
    nudge_x = 15, nudge_y = 0) # label data points above pval line of interest

effect + theme_classic() + 
  scale_x_discrete(limits=c(1,481)) + # discretize x axis
  scale_y_continuous()
```

```
ggsave("../feral_analysis_output/LeaveOneOut_effect.png")
```
