## Supplementary Figure 5 for "Parasite and pathogen prevalence in our closest animal companions is determined by accessibility of sanitation services": SuppFig5_Correlation_Supplement.html

Correlation\_Supplement.knit


### Independent variable correlations and model selection

#### Purpose

The purpose of this document is to demonstrate the process by which the list of models in the present manuscript were chosen.

**This was done through the following steps:**  
1- **Tests for collinearity** among independant variables with variance inflation factor (VIF), and for correlation with a Pearson/Polyserial correlation matrix  
2- **Redundancy/Confounding** Variables which were correlated were compared between univariable and multivariable models to identify redundancy or confounding  
3- **Model selection** Variables which were correlated and redundant were not included in the same model (because one of the variables was not explanatory when included in the multivariable model). Variables that were correlated and confounding were analyzed only in a multivariable model together (because univariable modeling with confounding variables can lead to biased estimates). Variables which were neither redundant nor confounding were compared in both univariable and multivariable models together.

#### Testing for collinearity among independent variables

**Variance Inflation Factor (VIF)**  
The variance inflation factor (VIF) is used to identify collinearity among independent variables. Commonly, if VIF > 4, variables are considered collinear (Craney and Surles 2007).

```
##        RMSE       R2
## 1 0.2449186 0.032618
```

```
##                GVIF Df GVIF^(1/(2*Df))
## gdp        1.659511  1        1.288220
## sanitation 1.703542  1        1.305198
## nbii       2.037698  1        1.427480
## island     1.666599  2        1.136208
## gini       2.036584  1        1.427089
## ablat      2.017478  1        1.420379
```

All VIF < 4, therefore there is no collinearity between independent variables.

**Correlation Matrix**  
In addition to the VIF, a series of Polyserial and Pearson correlations was run in order to be conservative about correlations in the model leading to biased estimates.

*Scale for Pearson and Polyserial correlations*

High correlation: between +/- 0.50 and +/- 1  
Moderate correlation: +/- 0.30 and +/- 0.49  
Low correlation: between 0 and +/- 0.29

**Summary of correlation plot findings:**

1. `gini` correlates moderately with `island` and `ablat`, highyl with `nbii`
2. `gdp` correlates highly with `island` and `sanitation` and moderately with `ablat`
3. `nbii` correlates moderately with `ablat` and `gini` and moderately with `island`
4. `sanitation` correlates moderately with`ablat` and `island`, highly with `gdp`
5. `ablat` correlates moderately with `gini`, `sanitation`, and `nbii` and moderately with `gdp`
6. `island` correlates highly with `gdp` and moderately with `nbii`, `gini`, and `san`

#### Testing for redundant/confounding variables

Next, multivariable models will be tested among significant correlated variables to identify redundancy/confounding. Redundant variables are variables that do not add new information to a model (Song et al. 2018), and can be identified when the effect of one variable changes between the univariable and multivariable models. Thus, inclusion of the less important redundant variable only takes away power and does not need to be included in the multivariable model. Confounding is when there is an effect on the model that is not accounted for, resulting in biased estimates of effect (Kaplan and Berry 1990). Confounding variables are variables which are correlated with both another independent variable, and the dependent variable (Kaplan and Berry 1990). Variables which are correlated with other variables and redundant will not be considered in multivariable models together. Confounding variables must be kept in the same model to avoid biased estimates (Kaplan and Berry 1990).

##### `gdp` Potential Confounding/Redundancy

**`gdp` and `sanitation`**

*`gdp` univariable model*

```
## Generalized linear mixed model fit by maximum likelihood (Laplace
##   Approximation) [glmerMod]
##  Family: binomial  ( logit )
## Formula: prevalence ~ gdp_10000 + (1 | pathogen) + (1 | study) + (1 |  
##     study:uniq)
##    Data: feral
## Weights: N
## Control: 
## glmerControl(optimizer = "optimx", calc.derivs = FALSE, optCtrl = list(method = "nlminb",  
##     starttests = FALSE, kkt = FALSE))
## 
##      AIC      BIC   logLik deviance df.resid 
##  11712.3  11738.8  -5851.1  11702.3     1475 
## 
## Scaled residuals: 
##      Min       1Q   Median       3Q      Max 
## -1.25126 -0.18344 -0.00793  0.07806  1.30181 
## 
## Random effects:
##  Groups     Name        Variance Std.Dev.
##  study:uniq (Intercept) 2.0972   1.4482  
##  study      (Intercept) 0.7302   0.8545  
##  pathogen   (Intercept) 0.7210   0.8491  
## Number of obs: 1480, groups:  study:uniq, 1480; study, 445; pathogen, 361
## 
## Fixed effects:
##             Estimate Std. Error z value Pr(>|z|)    
## (Intercept) -1.95015    0.12986 -15.017   <2e-16 ***
## gdp_10000   -0.09953    0.04992  -1.994   0.0462 *  
## ---
## Signif. codes:  0 '***' 0.001 '**' 0.01 '*' 0.05 '.' 0.1 ' ' 1
## 
## Correlation of Fixed Effects:
##           (Intr)
## gdp_10000 -0.709
```

*`sanitation` univariable model*

```
## Generalized linear mixed model fit by maximum likelihood (Laplace
##   Approximation) [glmerMod]
##  Family: binomial  ( logit )
## Formula: prevalence ~ san_10 + (1 | pathogen) + (1 | study) + (1 | study:uniq)
##    Data: feral
## Weights: N
## Control: 
## glmerControl(optimizer = "optimx", calc.derivs = FALSE, optCtrl = list(method = "nlminb",  
##     starttests = FALSE, kkt = FALSE))
## 
##      AIC      BIC   logLik deviance df.resid 
##  11708.5  11735.0  -5849.3  11698.5     1475 
## 
## Scaled residuals: 
##      Min       1Q   Median       3Q      Max 
## -1.26353 -0.18191 -0.00903  0.07830  1.30843 
## 
## Random effects:
##  Groups     Name        Variance Std.Dev.
##  study:uniq (Intercept) 2.0986   1.4487  
##  study      (Intercept) 0.7140   0.8450  
##  pathogen   (Intercept) 0.7087   0.8419  
## Number of obs: 1480, groups:  study:uniq, 1480; study, 445; pathogen, 361
## 
## Fixed effects:
##             Estimate Std. Error z value Pr(>|z|)   
## (Intercept) -1.13516    0.36840  -3.081  0.00206 **
## san_10      -0.11316    0.04044  -2.798  0.00514 **
## ---
## Signif. codes:  0 '***' 0.001 '**' 0.01 '*' 0.05 '.' 0.1 ' ' 1
## 
## Correlation of Fixed Effects:
##        (Intr)
## san_10 -0.969
```

*`gdp` and `sanitation` multivariable model*

```
## Generalized linear mixed model fit by maximum likelihood (Laplace
##   Approximation) [glmerMod]
##  Family: binomial  ( logit )
## Formula: prevalence ~ gdp_10000 + san_10 + (1 | pathogen) + (1 | method) +  
##     (1 | study) + (1 | study:uniq)
##    Data: feral
## Weights: N
## Control: 
## glmerControl(optimizer = "optimx", calc.derivs = FALSE, optCtrl = list(method = "nlminb",  
##     starttests = FALSE, kkt = FALSE))
## 
##      AIC      BIC   logLik deviance df.resid 
##  11679.0  11716.1  -5832.5  11665.0     1473 
## 
## Scaled residuals: 
##      Min       1Q   Median       3Q      Max 
## -1.29411 -0.18626 -0.01398  0.08101  1.31405 
## 
## Random effects:
##  Groups     Name        Variance Std.Dev.
##  study:uniq (Intercept) 2.0346   1.4264  
##  study      (Intercept) 0.5797   0.7614  
##  pathogen   (Intercept) 0.7157   0.8460  
##  method     (Intercept) 0.3000   0.5477  
## Number of obs: 1480, groups:  
## study:uniq, 1480; study, 445; pathogen, 361; method, 83
## 
## Fixed effects:
##             Estimate Std. Error z value Pr(>|z|)  
## (Intercept) -0.99138    0.39218  -2.528   0.0115 *
## gdp_10000   -0.03715    0.05682  -0.654   0.5132  
## san_10      -0.09779    0.04664  -2.097   0.0360 *
## ---
## Signif. codes:  0 '***' 0.001 '**' 0.01 '*' 0.05 '.' 0.1 ' ' 1
## 
## Correlation of Fixed Effects:
##           (Intr) g_1000
## gdp_10000  0.305       
## san_10    -0.899 -0.543
```

`san_10` appears to be a better predictor of parasite prevalence than `gdp`, suggesting that this is a more important factor. GDP is redundant because its effect size is changed quite a bit when sanitation is included in the model. Sanitation effect size changes only slightly. Loss of significance for both in multivar model, but this makes sense as including a redundant variable that takes away power.

**`gdp` and `ablat`**

*`gdp` univariable model*

```
## Generalized linear mixed model fit by maximum likelihood (Laplace
##   Approximation) [glmerMod]
##  Family: binomial  ( logit )
## Formula: prevalence ~ gdp_10000 + (1 | pathogen) + (1 | study) + (1 |  
##     study:uniq)
##    Data: feral
## Weights: N
## Control: 
## glmerControl(optimizer = "optimx", calc.derivs = FALSE, optCtrl = list(method = "nlminb",  
##     starttests = FALSE, kkt = FALSE))
## 
##      AIC      BIC   logLik deviance df.resid 
##  11712.3  11738.8  -5851.1  11702.3     1475 
## 
## Scaled residuals: 
##      Min       1Q   Median       3Q      Max 
## -1.25126 -0.18344 -0.00793  0.07806  1.30181 
## 
## Random effects:
##  Groups     Name        Variance Std.Dev.
##  study:uniq (Intercept) 2.0972   1.4482  
##  study      (Intercept) 0.7302   0.8545  
##  pathogen   (Intercept) 0.7210   0.8491  
## Number of obs: 1480, groups:  study:uniq, 1480; study, 445; pathogen, 361
## 
## Fixed effects:
##             Estimate Std. Error z value Pr(>|z|)    
## (Intercept) -1.95015    0.12986 -15.017   <2e-16 ***
## gdp_10000   -0.09953    0.04992  -1.994   0.0462 *  
## ---
## Signif. codes:  0 '***' 0.001 '**' 0.01 '*' 0.05 '.' 0.1 ' ' 1
## 
## Correlation of Fixed Effects:
##           (Intr)
## gdp_10000 -0.709
```

*`ablat` univariable model*

```
## Generalized linear mixed model fit by maximum likelihood (Laplace
##   Approximation) [glmerMod]
##  Family: binomial  ( logit )
## Formula: prevalence ~ ablat + (1 | pathogen) + (1 | study) + (1 | study:uniq)
##    Data: feral
## Weights: N
## Control: 
## glmerControl(optimizer = "optimx", calc.derivs = FALSE, optCtrl = list(method = "nlminb",  
##     starttests = FALSE, kkt = FALSE))
## 
##      AIC      BIC   logLik deviance df.resid 
##  11714.4  11740.9  -5852.2  11704.4     1475 
## 
## Scaled residuals: 
##     Min      1Q  Median      3Q     Max 
## -1.2554 -0.1845 -0.0094  0.0782  1.2996 
## 
## Random effects:
##  Groups     Name        Variance Std.Dev.
##  study:uniq (Intercept) 2.1003   1.4492  
##  study      (Intercept) 0.7375   0.8588  
##  pathogen   (Intercept) 0.7098   0.8425  
## Number of obs: 1480, groups:  study:uniq, 1480; study, 445; pathogen, 361
## 
## Fixed effects:
##              Estimate Std. Error z value Pr(>|z|)    
## (Intercept) -1.925342   0.178898  -10.76   <2e-16 ***
## ablat       -0.006712   0.005011   -1.34     0.18    
## ---
## Signif. codes:  0 '***' 0.001 '**' 0.01 '*' 0.05 '.' 0.1 ' ' 1
## 
## Correlation of Fixed Effects:
##       (Intr)
## ablat -0.860
```

*`gdp` and `ablat` multivariable model*

```
## Generalized linear mixed model fit by maximum likelihood (Laplace
##   Approximation) [glmerMod]
##  Family: binomial  ( logit )
## Formula: prevalence ~ gdp_10000 + ablat + (1 | pathogen) + (1 | study) +  
##     (1 | study:uniq)
##    Data: feral
## Weights: N
## Control: 
## glmerControl(optimizer = "optimx", calc.derivs = FALSE, optCtrl = list(method = "nlminb",  
##     starttests = FALSE, kkt = FALSE))
## 
##      AIC      BIC   logLik deviance df.resid 
##  11713.9  11745.7  -5851.0  11701.9     1474 
## 
## Scaled residuals: 
##      Min       1Q   Median       3Q      Max 
## -1.25670 -0.18380 -0.00700  0.07861  1.30485 
## 
## Random effects:
##  Groups     Name        Variance Std.Dev.
##  study:uniq (Intercept) 2.0984   1.4486  
##  study      (Intercept) 0.7270   0.8527  
##  pathogen   (Intercept) 0.7183   0.8476  
## Number of obs: 1480, groups:  study:uniq, 1480; study, 445; pathogen, 361
## 
## Fixed effects:
##              Estimate Std. Error z value Pr(>|z|)    
## (Intercept) -1.874506   0.181562 -10.324   <2e-16 ***
## gdp_10000   -0.086530   0.054429  -1.590    0.112    
## ablat       -0.003249   0.005450  -0.596    0.551    
## ---
## Signif. codes:  0 '***' 0.001 '**' 0.01 '*' 0.05 '.' 0.1 ' ' 1
## 
## Correlation of Fixed Effects:
##           (Intr) g_1000
## gdp_10000 -0.184       
## ablat     -0.700 -0.400
```

`ablat` effect reduced by third and `gdp` only slightly reduced effect in multivar model compared to univar model, ablat likely redundant

##### `gini` Potential Confounding/Redundancy

**`gini` and `nbii`**

*`gini` univariable model*

```
## Generalized linear mixed model fit by maximum likelihood (Laplace
##   Approximation) [glmerMod]
##  Family: binomial  ( logit )
## Formula: prevalence ~ gini + (1 | pathogen) + (1 | study) + (1 | study:uniq)
##    Data: feral
## Weights: N
## Control: 
## glmerControl(optimizer = "optimx", calc.derivs = FALSE, optCtrl = list(method = "nlminb",  
##     starttests = FALSE, kkt = FALSE))
## 
##      AIC      BIC   logLik deviance df.resid 
##  11715.0  11741.5  -5852.5  11705.0     1475 
## 
## Scaled residuals: 
##      Min       1Q   Median       3Q      Max 
## -1.26068 -0.18657 -0.00938  0.07853  1.29737 
## 
## Random effects:
##  Groups     Name        Variance Std.Dev.
##  study:uniq (Intercept) 2.1002   1.4492  
##  study      (Intercept) 0.7400   0.8602  
##  pathogen   (Intercept) 0.7125   0.8441  
## Number of obs: 1480, groups:  study:uniq, 1480; study, 445; pathogen, 361
## 
## Fixed effects:
##              Estimate Std. Error z value Pr(>|z|)    
## (Intercept) -2.502036   0.347485  -7.200    6e-13 ***
## gini         0.009437   0.008535   1.106    0.269    
## ---
## Signif. codes:  0 '***' 0.001 '**' 0.01 '*' 0.05 '.' 0.1 ' ' 1
## 
## Correlation of Fixed Effects:
##      (Intr)
## gini -0.965
```

*`nbi` univariable model*

```
## Generalized linear mixed model fit by maximum likelihood (Laplace
##   Approximation) [glmerMod]
##  Family: binomial  ( logit )
## Formula: prevalence ~ nbii + (1 | pathogen) + (1 | study) + (1 | study:uniq)
##    Data: feral
## Weights: N
## Control: 
## glmerControl(optimizer = "optimx", calc.derivs = FALSE, optCtrl = list(method = "nlminb",  
##     starttests = FALSE, kkt = FALSE))
## 
##      AIC      BIC   logLik deviance df.resid 
##  11716.0  11742.5  -5853.0  11706.0     1475 
## 
## Scaled residuals: 
##      Min       1Q   Median       3Q      Max 
## -1.23797 -0.18479 -0.00783  0.07829  1.29654 
## 
## Random effects:
##  Groups     Name        Variance Std.Dev.
##  study:uniq (Intercept) 2.0968   1.4480  
##  study      (Intercept) 0.7499   0.8660  
##  pathogen   (Intercept) 0.7151   0.8456  
## Number of obs: 1480, groups:  study:uniq, 1480; study, 445; pathogen, 361
## 
## Fixed effects:
##             Estimate Std. Error z value Pr(>|z|)    
## (Intercept)  -2.0431     0.2339  -8.734   <2e-16 ***
## nbii         -0.1410     0.3478  -0.405    0.685    
## ---
## Signif. codes:  0 '***' 0.001 '**' 0.01 '*' 0.05 '.' 0.1 ' ' 1
## 
## Correlation of Fixed Effects:
##      (Intr)
## nbii -0.920
```

*`gini` and `nbi` multivariable model*

```
## Generalized linear mixed model fit by maximum likelihood (Laplace
##   Approximation) [glmerMod]
##  Family: binomial  ( logit )
## Formula: prevalence ~ gini + nbii + (1 | pathogen) + (1 | method) + (1 |  
##     study) + (1 | study:uniq)
##    Data: feral
## Weights: N
## Control: 
## glmerControl(optimizer = "optimx", calc.derivs = FALSE, optCtrl = list(method = "nlminb",  
##     starttests = FALSE, kkt = FALSE))
## 
##      AIC      BIC   logLik deviance df.resid 
##  11685.6  11722.7  -5835.8  11671.6     1473 
## 
## Scaled residuals: 
##      Min       1Q   Median       3Q      Max 
## -1.28680 -0.18512 -0.01226  0.08018  1.29974 
## 
## Random effects:
##  Groups     Name        Variance Std.Dev.
##  study:uniq (Intercept) 2.0325   1.4256  
##  study      (Intercept) 0.6117   0.7821  
##  pathogen   (Intercept) 0.7207   0.8489  
##  method     (Intercept) 0.2975   0.5454  
## Number of obs: 1480, groups:  
## study:uniq, 1480; study, 445; pathogen, 361; method, 83
## 
## Fixed effects:
##              Estimate Std. Error z value Pr(>|z|)    
## (Intercept) -2.170486   0.360941  -6.013 1.82e-09 ***
## gini         0.013937   0.009984   1.396    0.163    
## nbii        -0.462347   0.405394  -1.140    0.254    
## ---
## Signif. codes:  0 '***' 0.001 '**' 0.01 '*' 0.05 '.' 0.1 ' ' 1
## 
## Correlation of Fixed Effects:
##      (Intr) gini  
## gini -0.697       
## nbii -0.090 -0.563
```

Both increase in effect size when in model together, decrease in p value, possible confounding

**`gini` and `ablat`**

*`gini` univariable model*

```
## Generalized linear mixed model fit by maximum likelihood (Laplace
##   Approximation) [glmerMod]
##  Family: binomial  ( logit )
## Formula: prevalence ~ gini + (1 | pathogen) + (1 | study) + (1 | study:uniq)
##    Data: feral
## Weights: N
## Control: 
## glmerControl(optimizer = "optimx", calc.derivs = FALSE, optCtrl = list(method = "nlminb",  
##     starttests = FALSE, kkt = FALSE))
## 
##      AIC      BIC   logLik deviance df.resid 
##  11715.0  11741.5  -5852.5  11705.0     1475 
## 
## Scaled residuals: 
##      Min       1Q   Median       3Q      Max 
## -1.26068 -0.18657 -0.00938  0.07853  1.29737 
## 
## Random effects:
##  Groups     Name        Variance Std.Dev.
##  study:uniq (Intercept) 2.1002   1.4492  
##  study      (Intercept) 0.7400   0.8602  
##  pathogen   (Intercept) 0.7125   0.8441  
## Number of obs: 1480, groups:  study:uniq, 1480; study, 445; pathogen, 361
## 
## Fixed effects:
##              Estimate Std. Error z value Pr(>|z|)    
## (Intercept) -2.502036   0.347485  -7.200    6e-13 ***
## gini         0.009437   0.008535   1.106    0.269    
## ---
## Signif. codes:  0 '***' 0.001 '**' 0.01 '*' 0.05 '.' 0.1 ' ' 1
## 
## Correlation of Fixed Effects:
##      (Intr)
## gini -0.965
```

*`ablat` univariable model*

```
## Generalized linear mixed model fit by maximum likelihood (Laplace
##   Approximation) [glmerMod]
##  Family: binomial  ( logit )
## Formula: prevalence ~ ablat + (1 | pathogen) + (1 | study) + (1 | study:uniq)
##    Data: feral
## Weights: N
## Control: 
## glmerControl(optimizer = "optimx", calc.derivs = FALSE, optCtrl = list(method = "nlminb",  
##     starttests = FALSE, kkt = FALSE))
## 
##      AIC      BIC   logLik deviance df.resid 
##  11714.4  11740.9  -5852.2  11704.4     1475 
## 
## Scaled residuals: 
##     Min      1Q  Median      3Q     Max 
## -1.2554 -0.1845 -0.0094  0.0782  1.2996 
## 
## Random effects:
##  Groups     Name        Variance Std.Dev.
##  study:uniq (Intercept) 2.1003   1.4492  
##  study      (Intercept) 0.7375   0.8588  
##  pathogen   (Intercept) 0.7098   0.8425  
## Number of obs: 1480, groups:  study:uniq, 1480; study, 445; pathogen, 361
## 
## Fixed effects:
##              Estimate Std. Error z value Pr(>|z|)    
## (Intercept) -1.925342   0.178898  -10.76   <2e-16 ***
## ablat       -0.006712   0.005011   -1.34     0.18    
## ---
## Signif. codes:  0 '***' 0.001 '**' 0.01 '*' 0.05 '.' 0.1 ' ' 1
## 
## Correlation of Fixed Effects:
##       (Intr)
## ablat -0.860
```

*`gini` and `ablat` multivariable model*

```
## Generalized linear mixed model fit by maximum likelihood (Laplace
##   Approximation) [glmerMod]
##  Family: binomial  ( logit )
## Formula: prevalence ~ gini + ablat + (1 | pathogen) + (1 | method) + (1 |  
##     study) + (1 | study:uniq)
##    Data: feral
## Weights: N
## Control: 
## glmerControl(optimizer = "optimx", calc.derivs = FALSE, optCtrl = list(method = "nlminb",  
##     starttests = FALSE, kkt = FALSE))
## 
##      AIC      BIC   logLik deviance df.resid 
##  11685.3  11722.4  -5835.7  11671.3     1473 
## 
## Scaled residuals: 
##      Min       1Q   Median       3Q      Max 
## -1.28692 -0.18227 -0.01319  0.07913  1.30379 
## 
## Random effects:
##  Groups     Name        Variance Std.Dev.
##  study:uniq (Intercept) 2.0356   1.4267  
##  study      (Intercept) 0.6020   0.7759  
##  pathogen   (Intercept) 0.7135   0.8447  
##  method     (Intercept) 0.3166   0.5626  
## Number of obs: 1480, groups:  
## study:uniq, 1480; study, 445; pathogen, 361; method, 83
## 
## Fixed effects:
##              Estimate Std. Error z value Pr(>|z|)    
## (Intercept) -1.723692   0.517858  -3.329 0.000873 ***
## gini         0.001142   0.009601   0.119 0.905286    
## ablat       -0.007267   0.005656  -1.285 0.198919    
## ---
## Signif. codes:  0 '***' 0.001 '**' 0.01 '*' 0.05 '.' 0.1 ' ' 1
## 
## Correlation of Fixed Effects:
##       (Intr) gini  
## gini  -0.911       
## ablat -0.720  0.516
```

`gini` appears redundant with ablat, reduced in effect size quite a bit while `ablat` does not change much.

##### `nbii` Potential Confounding/Redundancy

**`nbii` and `ablat`**

*`nbi` univariable model*

```
## Generalized linear mixed model fit by maximum likelihood (Laplace
##   Approximation) [glmerMod]
##  Family: binomial  ( logit )
## Formula: prevalence ~ nbii + (1 | pathogen) + (1 | study) + (1 | study:uniq)
##    Data: feral
## Weights: N
## Control: 
## glmerControl(optimizer = "optimx", calc.derivs = FALSE, optCtrl = list(method = "nlminb",  
##     starttests = FALSE, kkt = FALSE))
## 
##      AIC      BIC   logLik deviance df.resid 
##  11716.0  11742.5  -5853.0  11706.0     1475 
## 
## Scaled residuals: 
##      Min       1Q   Median       3Q      Max 
## -1.23797 -0.18479 -0.00783  0.07829  1.29654 
## 
## Random effects:
##  Groups     Name        Variance Std.Dev.
##  study:uniq (Intercept) 2.0968   1.4480  
##  study      (Intercept) 0.7499   0.8660  
##  pathogen   (Intercept) 0.7151   0.8456  
## Number of obs: 1480, groups:  study:uniq, 1480; study, 445; pathogen, 361
## 
## Fixed effects:
##             Estimate Std. Error z value Pr(>|z|)    
## (Intercept)  -2.0431     0.2339  -8.734   <2e-16 ***
## nbii         -0.1410     0.3478  -0.405    0.685    
## ---
## Signif. codes:  0 '***' 0.001 '**' 0.01 '*' 0.05 '.' 0.1 ' ' 1
## 
## Correlation of Fixed Effects:
##      (Intr)
## nbii -0.920
```

*`ablat` univariable model*

```
## Generalized linear mixed model fit by maximum likelihood (Laplace
##   Approximation) [glmerMod]
##  Family: binomial  ( logit )
## Formula: prevalence ~ ablat + (1 | pathogen) + (1 | study) + (1 | study:uniq)
##    Data: feral
## Weights: N
## Control: 
## glmerControl(optimizer = "optimx", calc.derivs = FALSE, optCtrl = list(method = "nlminb",  
##     starttests = FALSE, kkt = FALSE))
## 
##      AIC      BIC   logLik deviance df.resid 
##  11714.4  11740.9  -5852.2  11704.4     1475 
## 
## Scaled residuals: 
##     Min      1Q  Median      3Q     Max 
## -1.2554 -0.1845 -0.0094  0.0782  1.2996 
## 
## Random effects:
##  Groups     Name        Variance Std.Dev.
##  study:uniq (Intercept) 2.1003   1.4492  
##  study      (Intercept) 0.7375   0.8588  
##  pathogen   (Intercept) 0.7098   0.8425  
## Number of obs: 1480, groups:  study:uniq, 1480; study, 445; pathogen, 361
## 
## Fixed effects:
##              Estimate Std. Error z value Pr(>|z|)    
## (Intercept) -1.925342   0.178898  -10.76   <2e-16 ***
## ablat       -0.006712   0.005011   -1.34     0.18    
## ---
## Signif. codes:  0 '***' 0.001 '**' 0.01 '*' 0.05 '.' 0.1 ' ' 1
## 
## Correlation of Fixed Effects:
##       (Intr)
## ablat -0.860
```

\*`nbi` and `ablat` multivariable model

```
## Generalized linear mixed model fit by maximum likelihood (Laplace
##   Approximation) [glmerMod]
##  Family: binomial  ( logit )
## Formula: prevalence ~ nbii + ablat + (1 | pathogen) + (1 | method) + (1 |  
##     study) + (1 | study:uniq)
##    Data: feral
## Weights: N
## Control: 
## glmerControl(optimizer = "optimx", calc.derivs = FALSE, optCtrl = list(method = "nlminb",  
##     starttests = FALSE, kkt = FALSE))
## 
##      AIC      BIC   logLik deviance df.resid 
##  11683.0  11720.1  -5834.5  11669.0     1473 
## 
## Scaled residuals: 
##      Min       1Q   Median       3Q      Max 
## -1.28128 -0.18272 -0.01218  0.07997  1.30315 
## 
## Random effects:
##  Groups     Name        Variance Std.Dev.
##  study:uniq (Intercept) 2.0319   1.4254  
##  study      (Intercept) 0.5974   0.7729  
##  pathogen   (Intercept) 0.7160   0.8462  
##  method     (Intercept) 0.3234   0.5687  
## Number of obs: 1480, groups:  
## study:uniq, 1480; study, 445; pathogen, 361; method, 83
## 
## Fixed effects:
##              Estimate Std. Error z value Pr(>|z|)   
## (Intercept) -1.126537   0.410655  -2.743  0.00608 **
## nbii        -0.613837   0.398570  -1.540  0.12354   
## ablat       -0.012509   0.005785  -2.163  0.03058 * 
## ---
## Signif. codes:  0 '***' 0.001 '**' 0.01 '*' 0.05 '.' 0.1 ' ' 1
## 
## Correlation of Fixed Effects:
##       (Intr) nbii  
## nbii  -0.853       
## ablat -0.776  0.549
```

Both `nbii` and `ablat` change quite a bit between univar and multivar, and increase in effect size when included together in multivar model. Potential confounding, keep in model together.

##### `san_10` Potential Confounding/Redundancy

**`san` and `ablat`**

*`san` univariable model*

```
## Generalized linear mixed model fit by maximum likelihood (Laplace
##   Approximation) [glmerMod]
##  Family: binomial  ( logit )
## Formula: prevalence ~ san_10 + (1 | pathogen) + (1 | study) + (1 | study:uniq)
##    Data: feral
## Weights: N
## Control: 
## glmerControl(optimizer = "optimx", calc.derivs = FALSE, optCtrl = list(method = "nlminb",  
##     starttests = FALSE, kkt = FALSE))
## 
##      AIC      BIC   logLik deviance df.resid 
##  11708.5  11735.0  -5849.3  11698.5     1475 
## 
## Scaled residuals: 
##      Min       1Q   Median       3Q      Max 
## -1.26353 -0.18191 -0.00903  0.07830  1.30843 
## 
## Random effects:
##  Groups     Name        Variance Std.Dev.
##  study:uniq (Intercept) 2.0986   1.4487  
##  study      (Intercept) 0.7140   0.8450  
##  pathogen   (Intercept) 0.7087   0.8419  
## Number of obs: 1480, groups:  study:uniq, 1480; study, 445; pathogen, 361
## 
## Fixed effects:
##             Estimate Std. Error z value Pr(>|z|)   
## (Intercept) -1.13516    0.36840  -3.081  0.00206 **
## san_10      -0.11316    0.04044  -2.798  0.00514 **
## ---
## Signif. codes:  0 '***' 0.001 '**' 0.01 '*' 0.05 '.' 0.1 ' ' 1
## 
## Correlation of Fixed Effects:
##        (Intr)
## san_10 -0.969
```

*`ablat` univariable model*

```
## Generalized linear mixed model fit by maximum likelihood (Laplace
##   Approximation) [glmerMod]
##  Family: binomial  ( logit )
## Formula: prevalence ~ ablat + (1 | pathogen) + (1 | study) + (1 | study:uniq)
##    Data: feral
## Weights: N
## Control: 
## glmerControl(optimizer = "optimx", calc.derivs = FALSE, optCtrl = list(method = "nlminb",  
##     starttests = FALSE, kkt = FALSE))
## 
##      AIC      BIC   logLik deviance df.resid 
##  11714.4  11740.9  -5852.2  11704.4     1475 
## 
## Scaled residuals: 
##     Min      1Q  Median      3Q     Max 
## -1.2554 -0.1845 -0.0094  0.0782  1.2996 
## 
## Random effects:
##  Groups     Name        Variance Std.Dev.
##  study:uniq (Intercept) 2.1003   1.4492  
##  study      (Intercept) 0.7375   0.8588  
##  pathogen   (Intercept) 0.7098   0.8425  
## Number of obs: 1480, groups:  study:uniq, 1480; study, 445; pathogen, 361
## 
## Fixed effects:
##              Estimate Std. Error z value Pr(>|z|)    
## (Intercept) -1.925342   0.178898  -10.76   <2e-16 ***
## ablat       -0.006712   0.005011   -1.34     0.18    
## ---
## Signif. codes:  0 '***' 0.001 '**' 0.01 '*' 0.05 '.' 0.1 ' ' 1
## 
## Correlation of Fixed Effects:
##       (Intr)
## ablat -0.860
```

*`san` and `ablat` multivariable model*

```
## Generalized linear mixed model fit by maximum likelihood (Laplace
##   Approximation) [glmerMod]
##  Family: binomial  ( logit )
## Formula: prevalence ~ san_10 + ablat + (1 | pathogen) + (1 | method) +  
##     (1 | study) + (1 | study:uniq)
##    Data: feral
## Weights: N
## Control: 
## glmerControl(optimizer = "optimx", calc.derivs = FALSE, optCtrl = list(method = "nlminb",  
##     starttests = FALSE, kkt = FALSE))
## 
##      AIC      BIC   logLik deviance df.resid 
##  11679.3  11716.4  -5832.7  11665.3     1473 
## 
## Scaled residuals: 
##      Min       1Q   Median       3Q      Max 
## -1.29549 -0.18476 -0.01360  0.08098  1.31347 
## 
## Random effects:
##  Groups     Name        Variance Std.Dev.
##  study:uniq (Intercept) 2.0351   1.4266  
##  study      (Intercept) 0.5807   0.7620  
##  pathogen   (Intercept) 0.7106   0.8430  
##  method     (Intercept) 0.3049   0.5522  
## Number of obs: 1480, groups:  
## study:uniq, 1480; study, 445; pathogen, 361; method, 83
## 
## Fixed effects:
##              Estimate Std. Error z value Pr(>|z|)  
## (Intercept) -0.910696   0.373670  -2.437   0.0148 *
## san_10      -0.107607   0.043532  -2.472   0.0134 *
## ablat       -0.001897   0.005339  -0.355   0.7223  
## ---
## Signif. codes:  0 '***' 0.001 '**' 0.01 '*' 0.05 '.' 0.1 ' ' 1
## 
## Correlation of Fixed Effects:
##        (Intr) san_10
## san_10 -0.824       
## ablat  -0.002 -0.436
```

`ablat` is reduced to 1/7 of effect size in univar model, probably redundant because `san_10` doesn’t change much.

##### `island` Potential Confounding/Redundancy

**`island` and `gdp`**

*`island` univariable results*

```
## Generalized linear mixed model fit by maximum likelihood (Laplace
##   Approximation) [glmerMod]
##  Family: binomial  ( logit )
## Formula: prevalence ~ island + (1 | pathogen) + (1 | study) + (1 | study:uniq)
##    Data: feral
## Weights: N
## Control: 
## glmerControl(optimizer = "optimx", calc.derivs = FALSE, optCtrl = list(method = "nlminb",  
##     starttests = FALSE, kkt = FALSE))
## 
##      AIC      BIC   logLik deviance df.resid 
##  11717.7  11749.5  -5852.8  11705.7     1474 
## 
## Scaled residuals: 
##      Min       1Q   Median       3Q      Max 
## -1.24150 -0.18463 -0.00770  0.07771  1.29642 
## 
## Random effects:
##  Groups     Name        Variance Std.Dev.
##  study:uniq (Intercept) 2.0955   1.4476  
##  study      (Intercept) 0.7473   0.8645  
##  pathogen   (Intercept) 0.7154   0.8458  
## Number of obs: 1480, groups:  study:uniq, 1480; study, 445; pathogen, 361
## 
## Fixed effects:
##             Estimate Std. Error z value Pr(>|z|)    
## (Intercept) -2.15176    0.09681 -22.226   <2e-16 ***
## islandQ     -0.31739    1.70334  -0.186    0.852    
## islandY      0.12203    0.17774   0.687    0.492    
## ---
## Signif. codes:  0 '***' 0.001 '**' 0.01 '*' 0.05 '.' 0.1 ' ' 1
## 
## Correlation of Fixed Effects:
##         (Intr) islndQ
## islandQ -0.016       
## islandY -0.320  0.020
```

*`gdp` univariable results*

```
## Generalized linear mixed model fit by maximum likelihood (Laplace
##   Approximation) [glmerMod]
##  Family: binomial  ( logit )
## Formula: prevalence ~ gdp_10000 + (1 | pathogen) + (1 | study) + (1 |  
##     study:uniq)
##    Data: feral
## Weights: N
## Control: 
## glmerControl(optimizer = "optimx", calc.derivs = FALSE, optCtrl = list(method = "nlminb",  
##     starttests = FALSE, kkt = FALSE))
## 
##      AIC      BIC   logLik deviance df.resid 
##  11712.3  11738.8  -5851.1  11702.3     1475 
## 
## Scaled residuals: 
##      Min       1Q   Median       3Q      Max 
## -1.25126 -0.18344 -0.00793  0.07806  1.30181 
## 
## Random effects:
##  Groups     Name        Variance Std.Dev.
##  study:uniq (Intercept) 2.0972   1.4482  
##  study      (Intercept) 0.7302   0.8545  
##  pathogen   (Intercept) 0.7210   0.8491  
## Number of obs: 1480, groups:  study:uniq, 1480; study, 445; pathogen, 361
## 
## Fixed effects:
##             Estimate Std. Error z value Pr(>|z|)    
## (Intercept) -1.95015    0.12986 -15.017   <2e-16 ***
## gdp_10000   -0.09953    0.04992  -1.994   0.0462 *  
## ---
## Signif. codes:  0 '***' 0.001 '**' 0.01 '*' 0.05 '.' 0.1 ' ' 1
## 
## Correlation of Fixed Effects:
##           (Intr)
## gdp_10000 -0.709
```

*`island` and `gdp` multivariable results*

```
## Generalized linear mixed model fit by maximum likelihood (Laplace
##   Approximation) [glmerMod]
##  Family: binomial  ( logit )
## Formula: prevalence ~ island + gdp_10000 + (1 | pathogen) + (1 | method) +  
##     (1 | study) + (1 | study:uniq)
##    Data: feral
## Weights: N
## Control: 
## glmerControl(optimizer = "optimx", calc.derivs = FALSE, optCtrl = list(method = "nlminb",  
##     starttests = FALSE, kkt = FALSE))
## 
##      AIC      BIC   logLik deviance df.resid 
##  11683.1  11725.5  -5833.5  11667.1     1472 
## 
## Scaled residuals: 
##      Min       1Q   Median       3Q      Max 
## -1.28382 -0.18166 -0.01390  0.07868  1.31303 
## 
## Random effects:
##  Groups     Name        Variance Std.Dev.
##  study:uniq (Intercept) 2.0305   1.4250  
##  study      (Intercept) 0.5848   0.7647  
##  pathogen   (Intercept) 0.7303   0.8546  
##  method     (Intercept) 0.3033   0.5507  
## Number of obs: 1480, groups:  
## study:uniq, 1480; study, 445; pathogen, 361; method, 83
## 
## Fixed effects:
##             Estimate Std. Error z value Pr(>|z|)    
## (Intercept) -1.73282    0.17259 -10.040   <2e-16 ***
## islandQ     -0.73705    1.64902  -0.447    0.655    
## islandY      0.25184    0.17593   1.431    0.152    
## gdp_10000   -0.12244    0.04984  -2.457    0.014 *  
## ---
## Signif. codes:  0 '***' 0.001 '**' 0.01 '*' 0.05 '.' 0.1 ' ' 1
## 
## Correlation of Fixed Effects:
##           (Intr) islndQ islndY
## islandQ   -0.018              
## islandY   -0.013  0.020       
## gdp_10000 -0.472  0.023 -0.281
```

`island` effect doubles in multivar model. `gdp` doesn’t change much. potential confounding.

**`island` and `gini`**

*`island` univariable results*

```
## Generalized linear mixed model fit by maximum likelihood (Laplace
##   Approximation) [glmerMod]
##  Family: binomial  ( logit )
## Formula: prevalence ~ island + (1 | pathogen) + (1 | study) + (1 | study:uniq)
##    Data: feral
## Weights: N
## Control: 
## glmerControl(optimizer = "optimx", calc.derivs = FALSE, optCtrl = list(method = "nlminb",  
##     starttests = FALSE, kkt = FALSE))
## 
##      AIC      BIC   logLik deviance df.resid 
##  11717.7  11749.5  -5852.8  11705.7     1474 
## 
## Scaled residuals: 
##      Min       1Q   Median       3Q      Max 
## -1.24150 -0.18463 -0.00770  0.07771  1.29642 
## 
## Random effects:
##  Groups     Name        Variance Std.Dev.
##  study:uniq (Intercept) 2.0955   1.4476  
##  study      (Intercept) 0.7473   0.8645  
##  pathogen   (Intercept) 0.7154   0.8458  
## Number of obs: 1480, groups:  study:uniq, 1480; study, 445; pathogen, 361
## 
## Fixed effects:
##             Estimate Std. Error z value Pr(>|z|)    
## (Intercept) -2.15176    0.09681 -22.226   <2e-16 ***
## islandQ     -0.31739    1.70334  -0.186    0.852    
## islandY      0.12203    0.17774   0.687    0.492    
## ---
## Signif. codes:  0 '***' 0.001 '**' 0.01 '*' 0.05 '.' 0.1 ' ' 1
## 
## Correlation of Fixed Effects:
##         (Intr) islndQ
## islandQ -0.016       
## islandY -0.320  0.020
```

*`gini` univariable results*

```
## Generalized linear mixed model fit by maximum likelihood (Laplace
##   Approximation) [glmerMod]
##  Family: binomial  ( logit )
## Formula: prevalence ~ gini + (1 | pathogen) + (1 | study) + (1 | study:uniq)
##    Data: feral
## Weights: N
## Control: 
## glmerControl(optimizer = "optimx", calc.derivs = FALSE, optCtrl = list(method = "nlminb",  
##     starttests = FALSE, kkt = FALSE))
## 
##      AIC      BIC   logLik deviance df.resid 
##  11715.0  11741.5  -5852.5  11705.0     1475 
## 
## Scaled residuals: 
##      Min       1Q   Median       3Q      Max 
## -1.26068 -0.18657 -0.00938  0.07853  1.29737 
## 
## Random effects:
##  Groups     Name        Variance Std.Dev.
##  study:uniq (Intercept) 2.1002   1.4492  
##  study      (Intercept) 0.7400   0.8602  
##  pathogen   (Intercept) 0.7125   0.8441  
## Number of obs: 1480, groups:  study:uniq, 1480; study, 445; pathogen, 361
## 
## Fixed effects:
##              Estimate Std. Error z value Pr(>|z|)    
## (Intercept) -2.502036   0.347485  -7.200    6e-13 ***
## gini         0.009437   0.008535   1.106    0.269    
## ---
## Signif. codes:  0 '***' 0.001 '**' 0.01 '*' 0.05 '.' 0.1 ' ' 1
## 
## Correlation of Fixed Effects:
##      (Intr)
## gini -0.965
```

*`island` and `gini` multivariable results*

```
## Generalized linear mixed model fit by maximum likelihood (Laplace
##   Approximation) [glmerMod]
##  Family: binomial  ( logit )
## Formula: prevalence ~ island + gini + (1 | pathogen) + (1 | method) +  
##     (1 | study) + (1 | study:uniq)
##    Data: feral
## Weights: N
## Control: 
## glmerControl(optimizer = "optimx", calc.derivs = FALSE, optCtrl = list(method = "nlminb",  
##     starttests = FALSE, kkt = FALSE))
## 
##      AIC      BIC   logLik deviance df.resid 
##  11687.8  11730.2  -5835.9  11671.8     1472 
## 
## Scaled residuals: 
##      Min       1Q   Median       3Q      Max 
## -1.28831 -0.18378 -0.01283  0.07987  1.30413 
## 
## Random effects:
##  Groups     Name        Variance Std.Dev.
##  study:uniq (Intercept) 2.0329   1.4258  
##  study      (Intercept) 0.6046   0.7776  
##  pathogen   (Intercept) 0.7188   0.8478  
##  method     (Intercept) 0.3072   0.5543  
## Number of obs: 1480, groups:  
## study:uniq, 1480; study, 445; pathogen, 361; method, 83
## 
## Fixed effects:
##              Estimate Std. Error z value Pr(>|z|)    
## (Intercept) -2.308422   0.373791  -6.176 6.59e-10 ***
## islandQ     -0.620235   1.655651  -0.375    0.708    
## islandY      0.174369   0.174810   0.997    0.319    
## gini         0.009429   0.008454   1.115    0.265    
## ---
## Signif. codes:  0 '***' 0.001 '**' 0.01 '*' 0.05 '.' 0.1 ' ' 1
## 
## Correlation of Fixed Effects:
##         (Intr) islndQ islndY
## islandQ -0.014              
## islandY -0.278  0.030       
## gini    -0.913  0.012  0.229
```

Neither `gini` nor `island` change that much between univar and multivar models.

**`island` and `san_10`**

*`island` univariable results*

```
## Generalized linear mixed model fit by maximum likelihood (Laplace
##   Approximation) [glmerMod]
##  Family: binomial  ( logit )
## Formula: prevalence ~ island + (1 | pathogen) + (1 | study) + (1 | study:uniq)
##    Data: feral
## Weights: N
## Control: 
## glmerControl(optimizer = "optimx", calc.derivs = FALSE, optCtrl = list(method = "nlminb",  
##     starttests = FALSE, kkt = FALSE))
## 
##      AIC      BIC   logLik deviance df.resid 
##  11717.7  11749.5  -5852.8  11705.7     1474 
## 
## Scaled residuals: 
##      Min       1Q   Median       3Q      Max 
## -1.24150 -0.18463 -0.00770  0.07771  1.29642 
## 
## Random effects:
##  Groups     Name        Variance Std.Dev.
##  study:uniq (Intercept) 2.0955   1.4476  
##  study      (Intercept) 0.7473   0.8645  
##  pathogen   (Intercept) 0.7154   0.8458  
## Number of obs: 1480, groups:  study:uniq, 1480; study, 445; pathogen, 361
## 
## Fixed effects:
##             Estimate Std. Error z value Pr(>|z|)    
## (Intercept) -2.15176    0.09681 -22.226   <2e-16 ***
## islandQ     -0.31739    1.70334  -0.186    0.852    
## islandY      0.12203    0.17774   0.687    0.492    
## ---
## Signif. codes:  0 '***' 0.001 '**' 0.01 '*' 0.05 '.' 0.1 ' ' 1
## 
## Correlation of Fixed Effects:
##         (Intr) islndQ
## islandQ -0.016       
## islandY -0.320  0.020
```

*`sanitation` univariable results*

```
## Generalized linear mixed model fit by maximum likelihood (Laplace
##   Approximation) [glmerMod]
##  Family: binomial  ( logit )
## Formula: prevalence ~ san_10 + (1 | pathogen) + (1 | study) + (1 | study:uniq)
##    Data: feral
## Weights: N
## Control: 
## glmerControl(optimizer = "optimx", calc.derivs = FALSE, optCtrl = list(method = "nlminb",  
##     starttests = FALSE, kkt = FALSE))
## 
##      AIC      BIC   logLik deviance df.resid 
##  11708.5  11735.0  -5849.3  11698.5     1475 
## 
## Scaled residuals: 
##      Min       1Q   Median       3Q      Max 
## -1.26353 -0.18191 -0.00903  0.07830  1.30843 
## 
## Random effects:
##  Groups     Name        Variance Std.Dev.
##  study:uniq (Intercept) 2.0986   1.4487  
##  study      (Intercept) 0.7140   0.8450  
##  pathogen   (Intercept) 0.7087   0.8419  
## Number of obs: 1480, groups:  study:uniq, 1480; study, 445; pathogen, 361
## 
## Fixed effects:
##             Estimate Std. Error z value Pr(>|z|)   
## (Intercept) -1.13516    0.36840  -3.081  0.00206 **
## san_10      -0.11316    0.04044  -2.798  0.00514 **
## ---
## Signif. codes:  0 '***' 0.001 '**' 0.01 '*' 0.05 '.' 0.1 ' ' 1
## 
## Correlation of Fixed Effects:
##        (Intr)
## san_10 -0.969
```

*`island` and `sanitation` multivariable results*

```
## Generalized linear mixed model fit by maximum likelihood (Laplace
##   Approximation) [glmerMod]
##  Family: binomial  ( logit )
## Formula: prevalence ~ island + san_10 + (1 | pathogen) + (1 | method) +  
##     (1 | study) + (1 | study:uniq)
##    Data: feral
## Weights: N
## Control: 
## glmerControl(optimizer = "optimx", calc.derivs = FALSE, optCtrl = list(method = "nlminb",  
##     starttests = FALSE, kkt = FALSE))
## 
##      AIC      BIC   logLik deviance df.resid 
##  11678.5  11720.9  -5831.2  11662.5     1472 
## 
## Scaled residuals: 
##      Min       1Q   Median       3Q      Max 
## -1.29639 -0.18431 -0.01231  0.08047  1.32057 
## 
## Random effects:
##  Groups     Name        Variance Std.Dev.
##  study:uniq (Intercept) 2.0311   1.4252  
##  study      (Intercept) 0.5667   0.7528  
##  pathogen   (Intercept) 0.7125   0.8441  
##  method     (Intercept) 0.3087   0.5556  
## Number of obs: 1480, groups:  
## study:uniq, 1480; study, 445; pathogen, 361; method, 83
## 
## Fixed effects:
##             Estimate Std. Error z value Pr(>|z|)    
## (Intercept) -0.78375    0.37977  -2.064 0.039041 *  
## islandQ     -0.45885    1.64412  -0.279 0.780177    
## islandY      0.29846    0.17505   1.705 0.088197 .  
## san_10      -0.13430    0.04073  -3.298 0.000975 ***
## ---
## Signif. codes:  0 '***' 0.001 '**' 0.01 '*' 0.05 '.' 0.1 ' ' 1
## 
## Correlation of Fixed Effects:
##         (Intr) islndQ islndY
## islandQ  0.028              
## islandY  0.200  0.036       
## san_10  -0.916 -0.034 -0.290
```

effect size of `island` changes a bit.. nearly doubles. p value gets quite a bit lower so potentially some confounding with `sanitation`. `sanitation` doesn’t change much in effect size, but more significant in the multivariable model.

**`island` and `nbii`**

*`island` univariable results*

```
## Generalized linear mixed model fit by maximum likelihood (Laplace
##   Approximation) [glmerMod]
##  Family: binomial  ( logit )
## Formula: prevalence ~ island + (1 | pathogen) + (1 | study) + (1 | study:uniq)
##    Data: feral
## Weights: N
## Control: 
## glmerControl(optimizer = "optimx", calc.derivs = FALSE, optCtrl = list(method = "nlminb",  
##     starttests = FALSE, kkt = FALSE))
## 
##      AIC      BIC   logLik deviance df.resid 
##  11717.7  11749.5  -5852.8  11705.7     1474 
## 
## Scaled residuals: 
##      Min       1Q   Median       3Q      Max 
## -1.24150 -0.18463 -0.00770  0.07771  1.29642 
## 
## Random effects:
##  Groups     Name        Variance Std.Dev.
##  study:uniq (Intercept) 2.0955   1.4476  
##  study      (Intercept) 0.7473   0.8645  
##  pathogen   (Intercept) 0.7154   0.8458  
## Number of obs: 1480, groups:  study:uniq, 1480; study, 445; pathogen, 361
## 
## Fixed effects:
##             Estimate Std. Error z value Pr(>|z|)    
## (Intercept) -2.15176    0.09681 -22.226   <2e-16 ***
## islandQ     -0.31739    1.70334  -0.186    0.852    
## islandY      0.12203    0.17774   0.687    0.492    
## ---
## Signif. codes:  0 '***' 0.001 '**' 0.01 '*' 0.05 '.' 0.1 ' ' 1
## 
## Correlation of Fixed Effects:
##         (Intr) islndQ
## islandQ -0.016       
## islandY -0.320  0.020
```

*`nbi` univariable results*

```
## Generalized linear mixed model fit by maximum likelihood (Laplace
##   Approximation) [glmerMod]
##  Family: binomial  ( logit )
## Formula: prevalence ~ nbii + (1 | pathogen) + (1 | study) + (1 | study:uniq)
##    Data: feral
## Weights: N
## Control: 
## glmerControl(optimizer = "optimx", calc.derivs = FALSE, optCtrl = list(method = "nlminb",  
##     starttests = FALSE, kkt = FALSE))
## 
##      AIC      BIC   logLik deviance df.resid 
##  11716.0  11742.5  -5853.0  11706.0     1475 
## 
## Scaled residuals: 
##      Min       1Q   Median       3Q      Max 
## -1.23797 -0.18479 -0.00783  0.07829  1.29654 
## 
## Random effects:
##  Groups     Name        Variance Std.Dev.
##  study:uniq (Intercept) 2.0968   1.4480  
##  study      (Intercept) 0.7499   0.8660  
##  pathogen   (Intercept) 0.7151   0.8456  
## Number of obs: 1480, groups:  study:uniq, 1480; study, 445; pathogen, 361
## 
## Fixed effects:
##             Estimate Std. Error z value Pr(>|z|)    
## (Intercept)  -2.0431     0.2339  -8.734   <2e-16 ***
## nbii         -0.1410     0.3478  -0.405    0.685    
## ---
## Signif. codes:  0 '***' 0.001 '**' 0.01 '*' 0.05 '.' 0.1 ' ' 1
## 
## Correlation of Fixed Effects:
##      (Intr)
## nbii -0.920
```

*`nbi` and `island` multivariable results*

```
## Generalized linear mixed model fit by maximum likelihood (Laplace
##   Approximation) [glmerMod]
##  Family: binomial  ( logit )
## Formula: prevalence ~ island + nbii + (1 | pathogen) + (1 | method) +  
##     (1 | study) + (1 | study:uniq)
##    Data: feral
## Weights: N
## Control: 
## glmerControl(optimizer = "optimx", calc.derivs = FALSE, optCtrl = list(method = "nlminb",  
##     starttests = FALSE, kkt = FALSE))
## 
##      AIC      BIC   logLik deviance df.resid 
##  11688.7  11731.1  -5836.4  11672.7     1472 
## 
## Scaled residuals: 
##      Min       1Q   Median       3Q      Max 
## -1.26832 -0.18429 -0.01371  0.08011  1.30688 
## 
## Random effects:
##  Groups     Name        Variance Std.Dev.
##  study:uniq (Intercept) 2.0286   1.4243  
##  study      (Intercept) 0.6129   0.7829  
##  pathogen   (Intercept) 0.7240   0.8509  
##  method     (Intercept) 0.3148   0.5611  
## Number of obs: 1480, groups:  
## study:uniq, 1480; study, 445; pathogen, 361; method, 83
## 
## Fixed effects:
##             Estimate Std. Error z value Pr(>|z|)    
## (Intercept)  -1.8090     0.2599  -6.960 3.41e-12 ***
## islandQ      -0.6672     1.6573  -0.403    0.687    
## islandY       0.1431     0.1725   0.830    0.407    
## nbii         -0.1877     0.3388  -0.554    0.579    
## ---
## Signif. codes:  0 '***' 0.001 '**' 0.01 '*' 0.05 '.' 0.1 ' ' 1
## 
## Correlation of Fixed Effects:
##         (Intr) islndQ islndY
## islandQ -0.023              
## islandY  0.015  0.024       
## nbii    -0.807  0.023 -0.144
```

Neither `nbii` nor `island` change that much between univar and multivar models

##### Summary of redundancy/confounding results

1. `gini` correlates moderately with `island` and `ablat`, `nbii`  
   **redundant/confounding?:** `gini` is redundant with `san_10` and `ablat`, may be confounding with `nbii`
2. `gdp` correlates highly with `island` and `sanitation` and moderately with `ablat`  
   **redundant/confounding?:** `gdp` redundant with sanitation, `ablat` redundant with `gdp`
3. `nbii` correlates moderately with `ablat` and `gini` and moderately with `island`  
   **redundant/confounding?:** `nbii` is confounding with `ablat`
4. `sanitation` correlates highly with `gdp`, `ablat` and `island`  
   **redundant/confounding?:** `gdp` and `ablat` redundant with `sanitation`
5. `ablat` correlates highly with `gini`, `sanitation`, and `nbii` and moderately with `gdp`  
   **redundant/confounding?:** confounding with `nbii`, redundant with `gdp` and `sanitation`
6. `island` correlates highly with `gini` and `gdp`, `san`, and moderately with `nbii`  
   **redundant/confounding?:** potential confounding with `gdp` and `sanitation`

##### Summarized Rules for model selection

-do not consider `gdp` in models with `san_10` or `ablat`, because it is redundant with both  
-`nbii` needs to be in same model as `ablat`, because they are confounding variables  
-do not consider `san_10` in models with `gdp` or `ablat`, because the latter two are redundant with sanitation  
-`island` increases slightly in effect size with `gdp` and with `san_10`, and both of these variables change somewhat in effect size in multivariable model– consider in multivariable models due to possible confounding.

**List of models to consider based on the above results:**  
0. Null model baseline  
1. san\_10  
2. san\_10 + island  
3. gdp  
4. gdp + island  
5. nbii + ablat  
6. gini
